# Seagrass genomes reveal a hexaploid ancestry facilitating adaptation to the marine environment

**DOI:** 10.1101/2023.03.05.531170

**Authors:** Xiao Ma, Steffen Vanneste, Jiyang Chang, Luca Ambrosino, Kerrie Barry, Till Bayer, Alexander A. Bobrov, LoriBeth Boston, Justin E Campbell, Hengchi Chen, Maria Luisa Chiusano, Emanuela Dattolo, Jane Grimwood, Guifen He, Jerry Jenkins, Marina Khachaturyan, Lázaro Marín-Guirao, Attila Mesterházy, Danish-Daniel Muhd, Jessica Pazzaglia, Chris Plott, Shanmugam Rajasekar, Stephane Rombauts, Miriam Ruocco, Alison Scott, Min Pau Tan, Jozefien Van de Velde, Bartel Vanholme, Jenell Webber, Li Lian Wong, Mi Yan, Yeong Yik Sung, Polina Novikova, Jeremy Schmutz, Thorsten Reusch, Gabriele Procaccini, Jeanine Olsen, Yves Van de Peer

**Affiliations:** Department of Plant Biotechnology and Bioinformatics, Ghent University, 9052 Ghent, Belgium; VIB Center for Plant Systems Biology, VIB, 9052 Ghent, Belgium; Department Plants and Crops, Faculty of Bioscience Engineering, Ghent University, Coupure Links 653, 9000 Ghent, Belgium; Department of Research Infrastructure for Marine Biological Resources, Stazione Zoologica Anton Dohrn, Villa Comunale, 80121, Naples, Italy; DOE Joint Genome Institute, Lawrence Berkeley National Laboratory, Mail Stop: 91R183, 1 Cyclotron Road, Berkeley, CA 94720, USA; GEOMAR Helmholtz-Zentrum für Ozeanforschung Kiel, Düsternbrooker Weg 20, 24105 Kiel, Germany; Papanin Institute for Biology of Inland Waters RAS, Borok, Nekouz Distr., Yaroslavl Reg., 152742, Russia; Genome Sequencing Center, Hudson Alpha Institute for Biotechnology, 601 Genome Way, Huntsville, AL 35806, USA; Coastlines and Oceans Division, Institute of Environment, Florida International University-Biscayne Bay Campus, 3000 NE 151^st^ St., Miami, Florida 33181, USA; Department of Agricultural Sciences, University Federico II of Naples, Naples, Italy; Department of Integrative Marine Ecology, Stazione Zoologica Anton Dohrn, Villa Comunale, 80121 Naples, Italy; Institute of General Microbiology, University of Kiel, Kiel, Germany; Seagrass Ecology Group, Oceanographic Center of Murcia, Spanish Institute of Oceanography (IEO-CSIC), Murcia, Spain; Centre for Ecological Research, Wetland Ecology Research Group, Bem tér 18/C, Debrecen, H-4026, Hungary; Institute of Marine Biotechnology, Universiti Malaysia Terengganu, Terengganu, Malaysia; Arizona Genomics Institute, University of Arizona, 1657 E. Helen St, Tucson, AZ 85721, USA; Max Planck Institute for Plant Breeding Research (MPIPZ), Department of Chromosome Biology, Carl-von-Linne-Weg 10, 50829 Köln, Germany; Marine Evolutionary Ecology, GEOMAR Helmholtz Centre for Ocean Research Kiel, Düsternbrooker Weg 20, D-24105 Kiel, Germany; Groningen Institute for Evolutionary Life Sciences (GELIFES), University of Groningen, Nijenborgh 7, 9747AG, Groningen, Netherlands; Centre for Microbial Ecology and Genomics, Department of Biochemistry, Genetics and Microbiology, University of Pretoria, Pretoria 0028, South Africa; College of Horticulture, Academy for Advanced Interdisciplinary Studies, Nanjing Agricultural University, Nanjing, China

**Keywords:** Alismatales, seagrasses, *Posidonia oceanica*, *Thalassia testudinum*, *Cymodocea nodosa*, *Potamogeton acutifolium*, *Zostera marina*, whole genome duplication (WGD), whole genome triplication (WGT) hexaploidy, convergent evolution, salinity, light, carbon acquisition, temperature, volatiles, defense, lignification

## Abstract

Seagrasses comprise the only submerged marine angiosperms, a feat of adaptation from three independent freshwater lineages within the Alismatales. These three parallel lineages offer the unique opportunity to study convergent versus lineage-specific adaptation to a fully marine lifestyle. Here, we present chromosome-level genome assemblies from a representative species of each of the seagrass lineages - *Posidonia oceanica* (Posidoniaceae), *Cymodocea nodosa* (Cymodoceaceae), and *Thalassia testudinum* (Hydrocharitaceae) *-* along with an improved assembly for *Zostera marina* (Zosteraceae). We also include a draft genome of *Potamogeton acutifolius*, a representative of Potamogetonaceae, the freshwater sister lineage to the Zosteraceae. Genome analysis reveals that all seagrasses share an ancient whole genome triplication (WGT) event, dating to the early evolution of the Alismatales. An additional whole genome duplication (WGD) event was uncovered for *C. nodosa* and *P. acutifolius*. Dating of ancient WGDs and more recent bursts of transposable elements correlate well with major geological and recent climatic events, supporting their role as rapid generators of genetic variation. Comparative analysis of selected gene families suggests that the transition from the submerged-freshwater to submerged-marine environment did not require revolutionary changes. Major gene losses related to, e.g., stomata, volatiles, defense, and lignification, are likely a consequence of the submerged lifestyle rather than the cause (‘use it or lose it’). Likewise, genes, often retained from the WGD and WGT, were co-opted for functions requiring the alignment of many small adaptations (‘tweaking’), e.g., osmoregulation, salinity, light capture, carbon acquisition, and temperature. Our ability to manage and conserve seagrass ecosystems depends on our understanding of the fundamental processes underpinning their resilience. These new genomes will accelerate functional studies and are expected to contribute to transformative solutions — as continuing worldwide losses of the ‘savannas of the sea’ are of major concern in times of climate change and loss of biodiversity.

## INTRODUCTION

Seagrasses are unique flowering plants, adapted to a fully submerged existence in the highly saline environment of the ocean, where they must root in reducing sediments and endure chronic light limitation. In spite of these obstacles, the 60 or so species are among the most widely distributed flowering plants ^1-3^ with current estimates of coverage ranging from 600,000 km^2 4^ to a modeled value of 1,6 million km^2 5,6^. Seagrasses fulfill critical ecosystem functions and services including coastal nurseries, nutrient cycling, bacterial suppression, and coastal erosion protection ^7-9^. Along with mangroves, saltmarshes, and coral reefs, seagrass meadows are among the most biologically productive ecosystems on Earth. They act as breeding and nursery grounds for a huge variety of organisms including juvenile and adult fish, epiphytic and free-living algae, mollusks, bristle worms, nematodes, and other invertebrates such as scallops, crabs, and shrimp. Their importance for marine megafauna is unrivalled and their disappearance an important driver of megafauna decline ^10^. Seagrasses also rank amongst the most efficient natural carbon sinks on Earth, sequestering CO_2_ through photosynthesis and storing organic carbon in sediments for millennia ^11^. While occupying only 0.1% of the ocean surface, seagrasses have been estimated to bury 27–44 Tg C_org_ per year globally, accounting for 10 - 18% of the total C burial in the oceans and being up to 40 times more efficient at capturing organic carbon than land forests soils ^12^.

Previous work in *Zostera marina* ^13,14^ uncovered several unique gene family losses, as well as metabolic pathway losses and gains, underlying novel structural and physiological traits, along with evidence for ancient polyploidy. Here, we expand on this work utilizing new chromosome-scale, high-quality reference genomes to understand the specific morphological and physiological adaptations that have enabled their global success from the tropics to the poles, except Antarctica ^1^. These included *Posidonia oceanica* (L.) Delile (Posidoniaceae), *Cymodocea nodosa* (Ucria) Ascherson (Cymodoceaceae), and *Thalassia testudinum* K. D. Koenig (Hydrocharitaceae) to chromosome level, and a closely related freshwater-submerged alismatid, *Potamogeton acutifolius* Link (Potamogetonaceae), to draft level. Representative species within families (Supplementary Fig. S1.1) were chosen based on importance and susceptibility to anthropogenic pressure, and the availability of an extensive ecological literature.

Briefly, *P. oceanica* is the iconic Mediterranean seagrass and largest in terms of plant size and physical biomass. It is a climax species characterized by extreme longevity and carbon storage capacity. *T. testudium* (turtle grass) is a climax tropical species unique to the greater Caribbean region, with a single sister species endemic to the Indo-Pacific. *C. nodosa* is restricted to the Mediterranean, Black and Caspian Seas, with an Atlantic extension along the Canary Island archipelago and along the subtropical Atlantic coast of Africa. It is the only temperate species of an otherwise disjunct tropical genus from the Indo-Pacific. The curly pondweed *P. acutifolius* belongs to the sister family of the Zosteraceae and was chosen as the closest submerged freshwater sister taxon. We also included the recently upgraded genome of *Zostera marina* L. ^15^, which is found throughout the northern hemisphere and arguably the most widespread species on the planet. To distinguish between adaptations to an aquatic lifestyle, and those to the unique ocean environment, our comparative analysis also included genomes of two additional recently sequenced emergent freshwater alismatids, along with the genomes of two salt-water tolerant mangrove species.

Having transitioned from a freshwater environment to a submerged saline environment on only three independent occasions is rare, we therefore assumed convergent evolution. To test this, we compared gene family evolution across species, considering gene loss, as well as gene birth through small and large-scale gene duplication events, and their effect on plant body structure (cell walls, stomata, hypolignification), as well as physiological adaptations (hypoxia, plant defense, secondary metabolites, light perception, carbon acquisition, heat shock factors and especially salt tolerance mechanisms).

## RESULTS AND DISCUSSION

### Genome assemblies and gene annotations

We assembled the genomes of *T. testudinum, P. oceanica*, and *C. nodosa* to chromosomal level using a combination of short sequence reads, PacBio HiFi, PacBio long reads, and Hi-C chromosome mapping. The novel seagrass genomes varied in haploid chromosome number from 6 to 18 and were very different in size, while containing approximately the same number of gene models (Table 1). Further details of genome assembly and annotation, based on a combination of *ab initio* prediction, homology searches, RNA-aided evidence, and manual curation can be found in Methods, Table 1, Supplementary Note S2.1, and Supplementary Table S2.1.3. BUSCO scores of >95% demonstrate the high level of completeness in the genomes. The prediction of non-protein coding RNA families (i.e., rRNAs, tRNAs, snoRNAs) for *Z. marina, C. nodosa, P. oceanica, T. testudinum*, and *P. acutifolius* can be found in Suppl. Note S3.1 and Suppl. Table S3.1. Figure 1 shows the distribution of different genomic features along the reconstructed pseudochromosomes for the different seagrasses. Information on plastid and mitochondrial genomes can be found respectively in Suppl. Note S2.2 and Suppl. Note S2.3.

**Table 1.** Primary genome assembly, annotation statistics and BUSCO completeness assessment of protein coding sequences. See Suppl. Table S2.1.3 for additional details for the alternate haplotypes.

| Statistics | *T. testudinum* | *P. oceanica* | *C. nodosa* | *Z. marina* v 3.1 <sup>15</sup> | *P. acutifolius* |
| --- | --- | --- | --- | --- | --- |
| **Assembly** |  |  |  |  |  |
| Haploid-chromosome number | 9 | 10 | 18 | 6 | 13 |
| Genome size (Mb) | 4261.9 | 2963.0 | 379.5 | 260.5 | 612 |
| Contig N50 (Mb) | 371.6 | 355.8 | 21.9 | 7.0 | 3.09 |
| Scaffold N50 (Mb) | 523.9 | 355.8 | 22.6 | 34.6 | 4.45 |
| **Annotation** |  |  |  |  |  |
| Protein coding genes | 25,665 | 23,306 | 20,563 | 22,256 | 21,277 |
| Mean gene length, bp | 19,151 | 7,017 | 5,866 | 3,237 | 3,448 |
| Mean CDS length, bp | 1,077 | 1,210 | 1,207 | 1,241 | 1,290 |
| Mean exon length, bp | 218 | 222 | 214 | 248 | 243 |
| Mean exon per gene | 4.95 | 5.46 | 5.65 | 5.01 | 5.3 |
| Mean intron length, bp | 4,576 | 1,303 | 1,003 | 499 | 503 |
| Number of introns > 1 kb % | 36.2 | 24.2 | 22.5 | 9.5 | 9.7 |
| Number of introns > 10 kb % | 13.5 | 2.6 | 1.4 | 0.6 | 0.3 |
| Number of introns > 20 kb % | 7.1 | 0.7 | 0.2 | 0.06 | 0.01 |
| Longest intron, bp | 283,604 | 224,280 | 89,280 | 46,497 | 34,817 |
| **Transcriptome support** |  |  |  |  |  |
| TPM > 0 % | 87.4 | 84.7 | 97.1 | 91.5 | 82.7 |
| TPM > 1 % | 72.9 | 70.1 | 86.6 | 80.2 | 75.6 |
| **BUSCO** |  |  |  |  |  |
| Complete % | **94.2** | **97.4** | **95.8** | **95.7** | **97.5** |
| single-copy % | 92.1 | 95.2 | 94.4 | 93.2 | 94.6 |
| duplicated % | 2.1 | 2.2 | 1.4 | 2.5 | 2.9 |
| Fragmented % | 1.1 | 0.2 | 0.3 | 80.5 | 0.2 |
| Missing % | 4.7 | 2.4 | 3.9 | 3.8 | 2.3 |
| **Functional annotation** |  |  |  |  |  |
|  | 92.3% | 96.8% | 91.8% | 96.2% | 98.9% |

**Figure 1.**
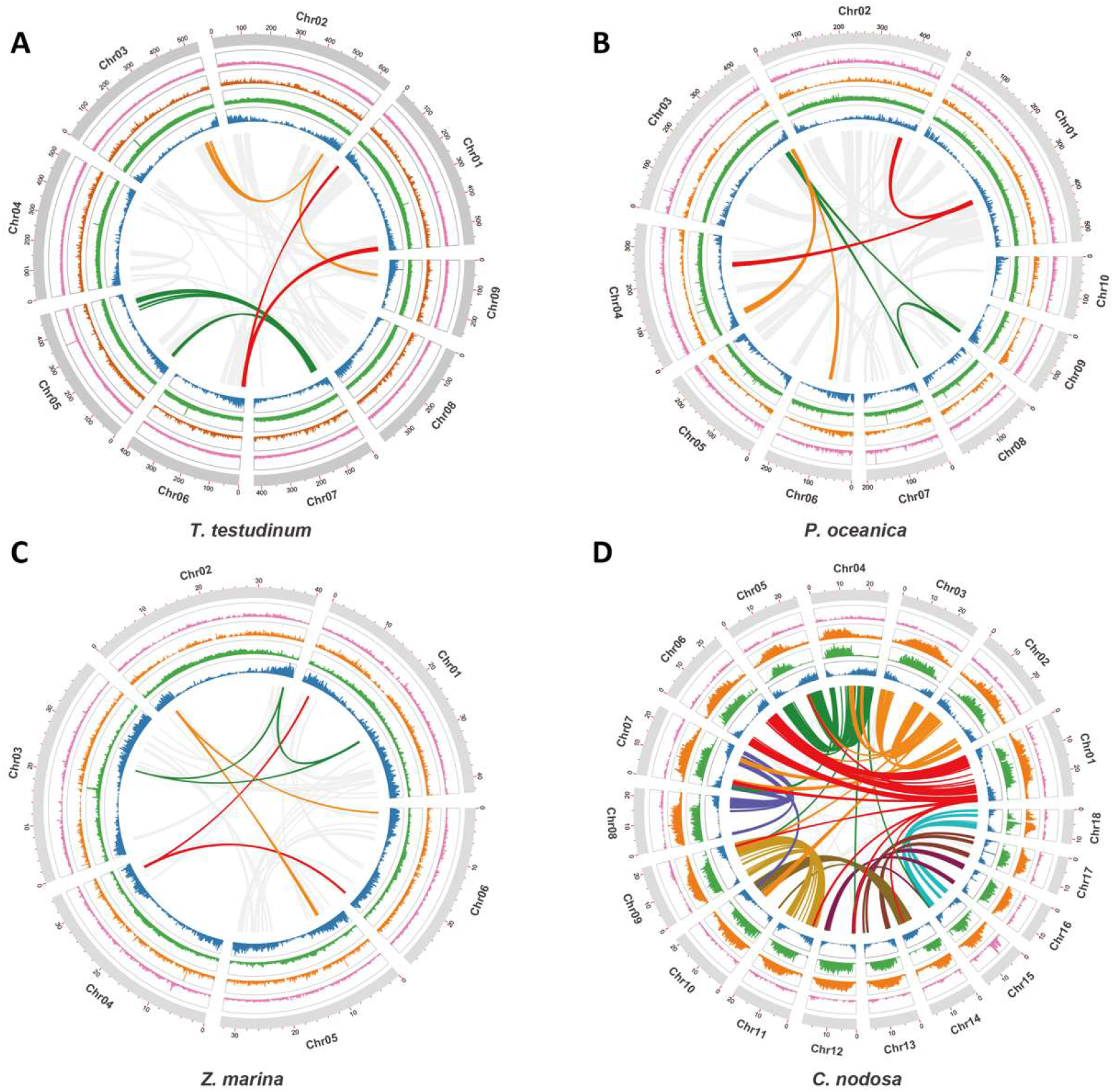
Distribution of the genomic features for *T. testudinum, P. oceanica, Z. marina* and *C. nodosa*. Tracks from the inner to outer side correspond to gene density (blue); LTR/Gypsy density (green); LTR/Copia (orange); DNA transposable elements (pink) and chromosomes (with length in Mb). Curved lines through the center denote synteny between different genomic regions. Grey lines in A, B and C reflect synteny involving the WGD, whereas the three colored lines represent synteny with WGTs. Colored lines in D represent synteny and strong intragenomic conservation and should not be compared with colors in A, B and C (see text for further details). The distribution of the genomic features for the longest scaffolds of *P. acutifolius*, can be found in Suppl. Fig. S2.1.2.

Information on Nuclear-mitochondria (NUMTs) and nuclear-chloroplast (NUPTs) integrants can be found in Suppl. Note S2.4 and Suppl. Table S2.4.

### Genome Evolution

#### Transposable elements

Transposable elements make up more than 85% of the genomes of *T. testudinum* and *P. oceanica*, as compared with 65% for *C. nodosa* and *Z. marina*, and 40% for *P. acutifolius* (Suppl. Table S4.1). Long terminal-repeat retrotransposons (LTR-REs) are the major class of TEs and account for 72.27%, 65.89%, 45.72% and 41.72% in *T. testudinum, P. oceanica, C. nodosa* and *Z. marina*, respectively. LTR/Gypsy elements account for 63.18% in *T. testudinum*, 57.8% in *P. oceanica* and 32.11% in *Z. marina*, whereas the proportion of LTR/Copia elements was higher than that of LTR/Gypsy in *C. nodosa* and *P. acutifolius*. Bursts of TEs (especially LTRs) create new genetic variation that may be adaptive under conditions of stress, and over evolutionary time different TE loads and distributions among species provide clues related to habitat differences and stress resistance ^16,17^. The insertion times of LTRs in the seagrass genomes (Methods) indicates a massive LTR/Gypsy burst around 200 thousand years ago (Kya) in *T. testudinum* (see y-axis), a moderate burst around 400 Kya in *P. oceanica* and *Z. marina*, but not in *C. nodosa*. By contrast, an expansion in Copia-elements happened around 2 Mya in *C. nodosa* but was weaker in *P. oceanica*, and nearly absent in *T. testudinum* and *Z. marina*. The recent TE gypsy burst (200 Kya) and older Copia burst (2 Mya median) correlate well with Pleistocene ice ages (Suppl. Fig. S4.1). The Gypsy bursts at 400 and 200 Kya correspond to Marine Isotope Stage MIS12 and MIS6, two heavy glaciations followed by rapid warming ^18^.

#### Whole genome duplication, ancient (hexa)polyploidy and dating

Next, we revisited the established WGD in *Z. marina* ^13^ and investigated whether evidence for ancient polyploidy could be found in the other seagrasses. To this end, we used inferred age distributions of synonymous substitution rate (K_S_), along with gene tree - species tree reconciliation methods (see Methods, Suppl. Note S4.2.1 and Suppl. Note S4.2.2). First, K_S_ distributions of all seagrass species showed peaks indicative of ancient WGDs (Suppl. Fig. S4.2.1)^14^. This was supported by intra- and inter-genomic collinearity analysis (see Suppl. Note S4.2.1.). Furthermore, comparison of *Z. marina, P. oceanica*, and *T. testudinum* with *Aristolochia fimbriata* - a magnoliid devoid of recent WGDs and therefore an excellent reference for the inference of angiosperm genome evolution ^19^ - shows a clear 3:1 synteny relationship (Suppl. Fig. S4.2.2). This implies, together with evidence from triplicated genomic blocks in intra-genome comparisons that *Z. marina, P. oceanica*, and *T. testudinum* (Figure 1) likely experienced an ancient hexaploidy. Interestingly, *C. nodosa* was found to show a 6:1 relationship compared to *A. fimbriata*, while showing a 2:1 relationship with its sister species *P. oceanica* (Suppl. Fig. S4.2.3), providing strong support for an additional WGD in *C. nodosa* after diverging from the *P. oceanica* lineage. Likewise, the freshwater species *P. acutifolius* was found to show a collinear relationship of 6:1 with *A. fimbriata* and a 2:1 relationship with *Z. marina* and especially *P. oceanica*, as well as a 2:2 relationship with *C. nodosa* (Suppl. Fig. S4.2.4). This provides evidence that also *P. acutifolius* experienced a lineage specific WGD event after its divergence with *Z. marina*.

Second, based on a K_S_ analysis using ksrates ^20^, we were able to confirm that the paleohexaploidy is shared by *P. oceanica, C. nodosa, Z. marina*, and *P. acutifolius*, while the analysis was inconclusive in *T. testudinum* (Suppl. Fig. S4.2.5). To resolve this issue, we applied a gene tree - species tree reconciliation approach using WHALE ^21^, which confirmed that the ancient WGT is shared by all seagrasses, as well as *P. acutifolius* (Suppl. Note S4.2.2 and Suppl. Fig. S4.2.6). Phylogenomic dating of the WGT (see Methods and Suppl. Note S4.2.3) further shows that most gene duplicates are reconciled on the branch leading to the most recent common ancestor (MRCA) of Potamogetonaceae, Zosteraceae, Posidoniaceae, Cymodoceaceae and Hydrocharitaceae, at approximately 86.96 (89.89 - 79.81) Mya (Figure 2 and Suppl. Fig. S4.2.7).

**Figure 2.**
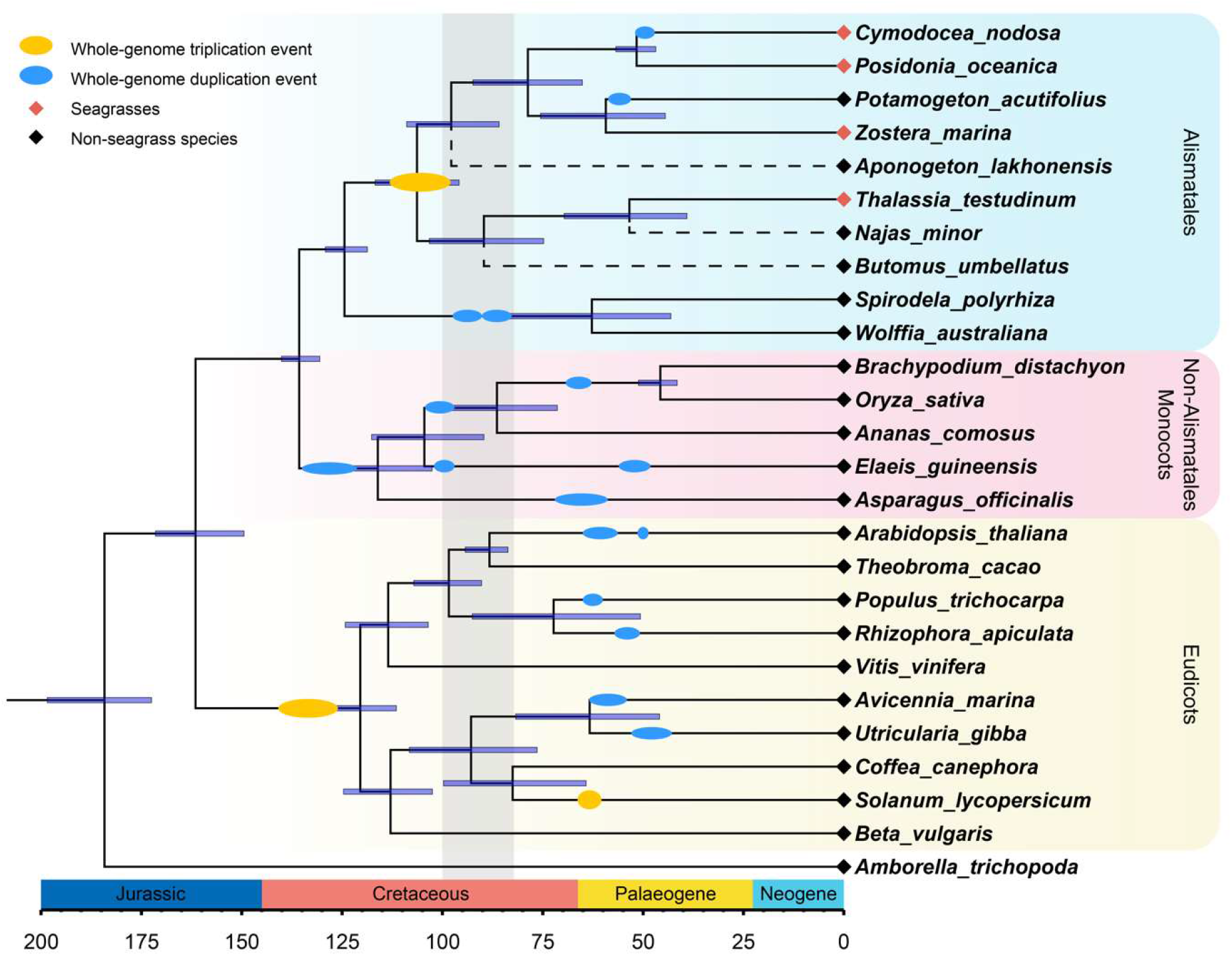
Time-calibrated phylogeny and WGT/WGD events across flowering plants with chromosome-level assembly. The tree was inferred from 146 single-copy genes and showing WGDs and WGTs based on inferences from the current study and previous analyses (Suppl. Table S4.2 and Suppl. Fig. S4.2.8). For a more comprehensive tree showing the phylogenetic position of seagrasses within Alismatales, see Suppl. Fig. S1.1. The dashed lines represent additional freshwater Alismatales species (phylogenetic position inferred using transcriptome data), mainly added for illustrative purposes to show non-monophyly of seagrass species. All branches have bootstrap support >98%. The light grey background denotes the Cenomanian-Turonian anoxic event (∼91± 8.6 Mya). See text and Methods for details.

### Adaptation to the Marine Environment

All three seagrass lineages characterized in this study share many specific morphological and physiological adaptations to their marine environment. Using a common set of species for which full genomes are available (four seagrasses, three freshwater alismatids, and 16 other angiosperms, Figure 2 and Suppl. Note S4.3), we broadly assessed commonalities and differences in gains and losses across gene families (further referred to as orthogroups, see Methods and Extended Data Table 1-13).

#### Use it or lose it

Stomata are not required and may even be harmful for a submerged lifestyle. Hence, seagrasses and to a limited extent also freshwater alismatids, e.g., *P. acutifolius*, have reduced the number of genes involved in their development. Specifically, out of 30 orthogroups containing guard cell toolkit genes ^22^, eleven have been convergently and completely lost in seagrasses, while six others were significantly contracted compared to non-seagrass genomes (Extended Data Table 1). Lost gene families include positive (SMF transcription factors), negative (EPIDERMAL PATTERNING FACTOR1 AND 2 (*EPF1, EPF2*), and TOO MANY MOUTHS (*TMM*)) regulators of stomatal development, as well as stomatal function (*BLUS1, KAT1/2* and *CHX20*) (Figures 3a and 3b). Gene losses and contractions in the guard cell toolkit are also seen in the submerged freshwater alismatid *P. acutifolius* studied here, but less extreme in the floating alismatid *S. polyrhiza* (Extended Data Table 1). See also ^14^.

**Figure 3.**
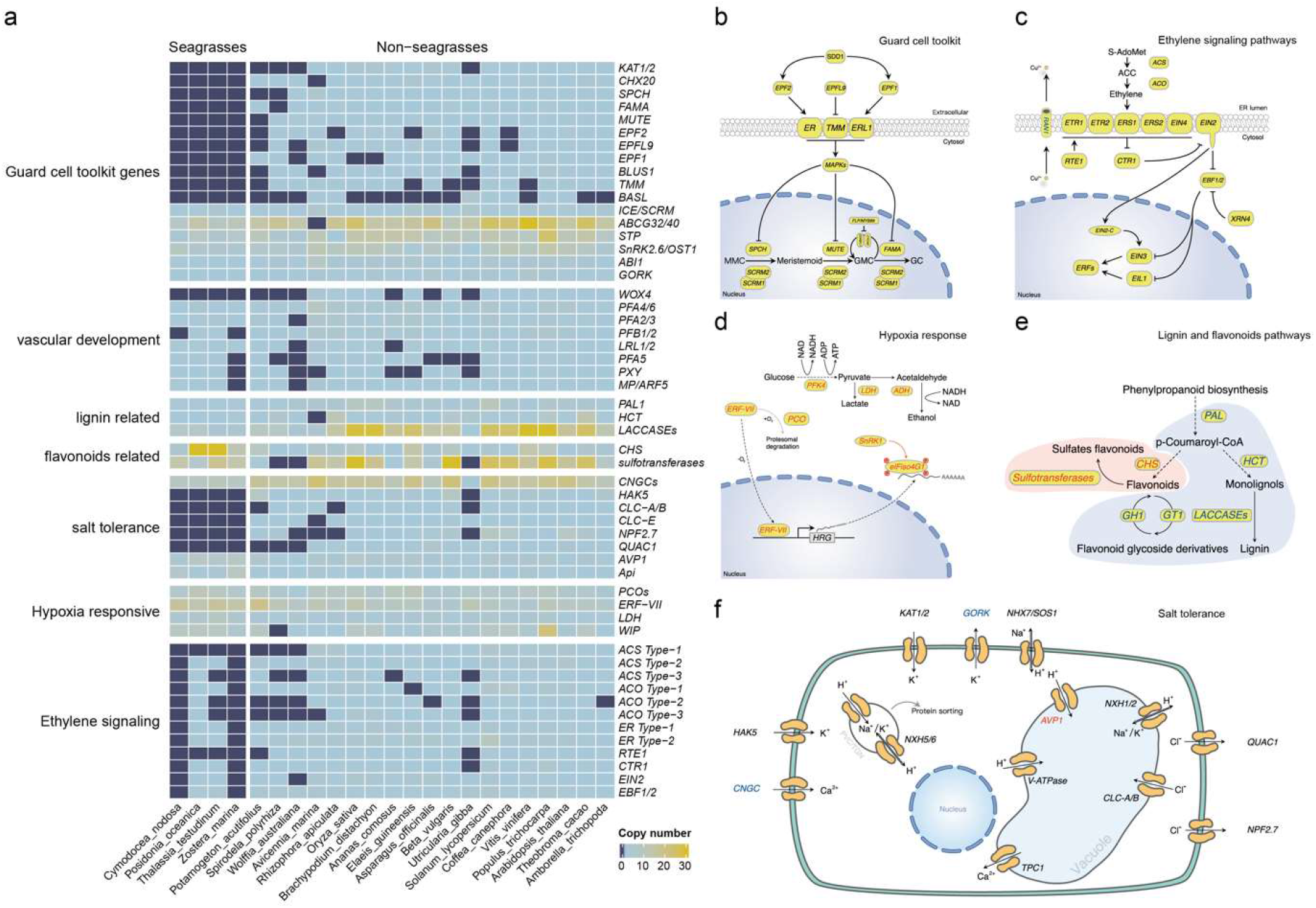
The loss, contraction, and expansion of gene families involved in the adaption to a marine environment. **a)** Gene copy numbers for 4 seagrasses and 19 representative non-seagrass species. **b)** Stomata differentiation from meristemoid mother cells (MMC) to guard mother cell (GMC), to guard cells. **c)** Ethylene synthesis and signaling. From panel a, we learn that all genes up to *EBF1/2* have been lost in *C. nodosa* and *Z. marina*, whereas *T. testudinum* and *P. oceanica* retained some genes. **d)** The hypoxia-responsive signaling in which the direct (*ERF-VII*) and indirect responsive (*SnRK1*) pathway are expanded. The rate-limiting enzyme (*PFK4*) in the glycolysis pathway, along with Lactate dehydrogenase (*LDH*), a rate-limiting enzyme in fermentation, are also expanded. **e)** Simplified schematic of the lignin and flavonoids biosynthesis pathways. Only steps that have significantly changed are shown. *PAL*, phenylalanine ammonialyase, is the gateway enzyme of the general phenylpropanoid pathway; *CHS*, chalcone synthase, the first enzyme of flavonoid biosynthesis, which directs the metabolic flux to flavonoid biosynthesis; *GT1*, flavonoid glycosyltransferases, is the final step of flavonoid biosynthesis to generate various flavonoid glycoside derivatives; *GH1*, flavonoid beta-glucosidase & myrosinase, is recycling of carbohydrate-based flavonoids; *HCT*, Hydroxycinnamoyl-CoA shikimate/quinate hydroxycinnamoyl transferase, channels phenylpropanoids via the “esters” pathway to monolignols”; *LACCASE*s are the last enzymes, which oxidize monolignols to facilitate their polymerization into lignin. **f)** Salt stress signaling implies different ion channels. HIGH-AFFINITY POTASSIUM TRANSPORTER 5 (*HAK5*) and selectivity filter are lost in seagrasses, allowing the efficient uptake of the essential ion K+ in seawater. Also, the Cl-transporter repertoire and CYCLIC NUCLEOTIDE GATE CATION CHANNELs are greatly reduced, however, Na^+^/H^+^ antiporters show no significant gene gains or losses. Vacuolar H^+^-PPases (*AVP1*) are expanded in all seagrasses. Panel d) e) f): genes in red are expanded; blue means contracted; The dashed line in the pathway means multiple metabolic steps.

The aqueous habitat of seagrasses, and their lack of stomata is also not conducive to emitting volatile substances as signals. Accordingly, we observed a convergent loss of orthogroups associated with volatile metabolites and signals. This includes the biosynthesis of triterpenes, and the volatile systemic acquired resistance signal, methyl salicylate ^23^ (Extended Data Table 2). Probably a more dramatic gene loss relates to ethylene biosynthesis and signaling (Extended Data Table 2). Two species, *C. nodosa* and *Z. marina*, do not contain *ACS* or *ACO* genes and are not expected to produce ACC or ethylene. Moreover, they seem to have lost the ability to respond to ethylene, as indicated by a severe contraction of the early ethylene signal transduction components (Figure 3a and 3c) ^13,14,24^. In contrast, the downstream ethylene transcription factors (*EIN3*/*EIL1/2*) have been retained in all seagrasses, suggesting they can still exert ethylene-independent functions. Remarkably, and unlike *C. nodosa* and *Z. marina, T. testudinum* and *P. oceanica* reduced, but did not entirely lose, the components for functional ethylene biosynthesis and signaling. One possible mechanism that may prevent the accumulation of deleterious levels of ethylene, and thus explain its retention in *T. testudinum* and *P. oceanica*, is via epiphytic and endophytic bacteria that express ACC deaminases. This hypothesis is supported by the presence of multiple ACC deaminases in the metagenome of *P. oceanica* sediments ^25^.

Seagrasses increase their morphological flexibility to withstand hydrodynamic wave and current forces by a reduction in vascular tissues, the main site of lignification ^26^. This is reflected in the absence of vascular proliferation factor *WOX4*, and a contraction of the number of pericycle cell identity transcription factors (Figure 3a and Extended Data Table 3). The most severe reduction of the vascular bundle is seen in *Z. marina* which even lacks a pericycle^27^, a finding that correlates with the loss and divergence of the vascular proliferation regulators *PXY* and *MONOPTEROS/ARF5* (Extended Data Table 3). Notably, the lack of *MONOPTEROS/ARF5* in *Z. marina* is further reflected in its inability to form an embryonic primary root ^28^. The general cellular hypolignification in seagrasses is reflected in the reduction in their number of *LACCASE*s, which oxidize monolignols to facilitate their polymerization into lignin ^29,30^ (Figures 3a, 3e and Extended Data Table 4). The reduced need for the monolignol production is matched by a reduction of respectively PHENYLALANINE AMMONIA LYASE (*PAL*), and HYDROXYCINNAMOYL-COA SHIKIMATE/QUINATE HYDROXYCINNAMOYL TRANSFERASE (*HCT*) genes, which constitute entrance points into phenylpropanoid biosynthesis ^31^ (Figures 3a, 3e and Extended Data Table 4).

The pathogen landscape of the marine environment is associated with a different composition of plant resistance (R-genes) genes. In the seagrasses, there are fewer genes containing nucleotide-binding leucine-rich repeat receptors (NLRs), as compared to most other plants (Extended Data Table 5, Suppl. Note S5.1, Suppl. Table S5.1 and Suppl. Fig. S5.1.1). Interestingly, NLRs with a TIR domain are completely absent in all seagrass lineages while a few such genes are missing the LRR domain. Lower counts of disease resistance genes have also been observed for other aquatic plants ^32,33^. Temperature fluctuations are much slower and show a lower amplitude in the marine compared to terrestrial environment ^34^. Accordingly, we observed a reduction in the number of plant heat shock transcription factors (*HSF*s) that are involved in the rapid activation of stress-responsive genes, and which have been linked to the evolutionary adaptation of plants to the terrestrial environment ^34^. Seagrasses contain only about half the number of *HSF*s as compared with terrestrial plants (Extended Data Table 5, Suppl. Note S5.2 and Suppl. Table S5.2). Interestingly, only seagrasses belonging to the tropical genera retained some of the key heat stress-related *HSF*s from WGD and WGT events (Extended Data Table 5), which could reflect their warmer native environment and higher heat stress tolerance compared to temperate seagrasses (*P. oceanica* and *Z. marina*).

### Multi-level tweaking to adapt to the marine environment

#### Protective flavonoids and phenolics

Most seagrasses, except *C. nodosa*, seem to have greatly expanded the number of CHALCON SYNTHASEs, which channel *p*-coumaroyl-CoA into flavonoid biosynthesis at the expense of monolignol biosynthesis (Extended Data Table 6). Flavonoids provide protection against UV and fungi, while enhancing recruitment of N-fixing bacteria ^25,35,36^. Flavonoids and other phenolics in seagrasses can be sulphated by the activity of cytosolic sulphotransferases to increase their water solubility, and bioactivity in the marine environment ^37,38^. For example, the sulphated monolignol, Zosteric Acid (*O*-sulfonated *p*-coumaric acid) is an antifouling agent which prevents biofilm formation at the leaf surface ^39^. In seagrasses, the orthogroup of cytosolic sulphotransferases was expanded, while flavonoid glycosyltransferases and flavonoid beta-glucosidases were contracted (Figure 3e and Extended Data Table 6). Jointly, these data illustrate how rerouting precursors of the lignin biosynthesis pathway facilitated two traits (reduced rigidity and protection) that contributed to the evolution of the marine lifestyle of seagrasses ^25,38^. In the case of *P. oceanica*, secreted phenolic compounds, together with anoxia inhibit microbial consumption of sucrose from root exudates ^25^.

#### Cellular salt tolerance

Salt tolerance in flowering plants is a complex trait, resulting from multiple cellular processes ^40^. In the extreme case of invasion of highly-saline, marine environments, one might assume wholesale changes in salt tolerance mechanisms themselves. Instead, it seems that canonical salt tolerance mechanisms were fine-tuned or “tweaked” towards higher efficiency on multiple levels. A major challenge associated with the marine environment is to prevent the accumulation of noxious levels of Na^+^ and Cl^-^, while allowing the efficient uptake of the essential ion K^+^. Angiosperms employ secondary Na^+^ transport mechanisms based on Na^+^/H^+^ antiporters fueled by a strong electrochemical H^+^ gradient. Surprisingly, no notable gene gains or losses were observed among the putative sodium transporting NHXs (*NHX1* and *SOS1/NHX7*), except for *C. nodosa*, which contains a few extra copies of *NHX1* and *SOS1* orthologs (Extended Data Table 7). Instead of an increased number of genes, we observed convergent amino acid substitutions in regulatory domains of *SOS1* orthologs in all four species (Suppl. Fig. S5.3.1), indicating altered regulation of *SOS1/NHX7* in these species, a notion that is also supported by the loss of *SOS3*, a key regulator of *SOS1* activity in *C. nodosa* (Extended Data Table 7). The electrochemical H^+^ gradients that fuel Na^+^ transport are established via H^+^ ATPases (*AHA*), V-ATPases and vacuolar H^+^-PPases (*AVP1*). Of these genes, only the *AVP1* genes were obviously expanded in all the seagrasses, containing almost twice the number of *AVP1* genes found on average in other angiosperms (Figure 3a and 3f). Interestingly, the expansion of *AVP1*-like genes can, at least partly, be linked to the ancient WGT followed by their specific retention, suggesting that these additional AVP copies were co-opted for adaptation to a marine lifestyle (Suppl. Fig. S5.4.2). Indeed, overexpression of such PPases has been shown to improve salt tolerance in several angiosperms (e.g., *Arabidopsis*, poplar, sugar cane) ^41-43^, by enhancing Na^+^ sequestration in the vacuole ^44^. Analysis of the K^+^-channel repertoire in seagrasses reveals an increased K^+^ uptake selectivity, as indicated by the loss of Shaker-type K^+^ channels with a TTGYGD-selectivity filter (Suppl. Fig. S5.3.2) ^45^, and a greatly reduced number of CYCLIC NUCLEOTIDE GATE CATION CHANNELs, that are relatively non-selective cation channels (Figure 3a, 3f and Extended Data Table 7). Moreover, the constant high K+ concentrations in seawater (9.7mM) renders high-affinity K+ transport systems superfluous, explaining the absence of *AtHAK5* in all seagrass genomes (Figure 3a and 3f). Also the Cl^-^ transporter repertoire is reduced in seagrasses (Figure 3a and 3f) and seagrasses lack orthologs for *NPF2*.*4* and *ALMT12/QUAC1, CLC-A, B* and *CLC-E*, likely reflecting their adaptation to a marine lifestyle (Figure 3a and 3f).

Elasticity of the cell wall is also a critical component of salt tolerance. This is mainly dictated by load bearing components of the cell wall, such as cellulose and pectins that cross-link the cellulose microfibrils. The bivalent cation Ca^2+^ stiffens the cell wall by establishing electrostatic bond between pectin strands. The excess of monovalent sodium in seawater may displace the divalent calcium and hinder dimerization of homogalacturonan chains that are present in canonical pectin ^46^. In addition to the canonical pectin polysaccharides, seagrasses deposit apiogalacturonan in their cell walls ^47^. The borate-bridges that cross-link apiogalacturonan chains are less sensitive to sodium displacement, providing an advantage to plants grown under high salt condition ^48^. One of the few known key enzymes in the synthesis of apiogalaturonan, is UDP-D-apiose/UDP-D-xylose synthase (*Api*), which converts UDP-D-glucuronate into UDP-D-apiose ^49^. Its expansion in seagrasses (in particular in *Zostera* and *Cymodocea*) is mirrored in the cell wall composition of seagrasses and therefore likely contributes to salt tolerance (Figure 3a). In addition, the apiogalactunonan could provide a way to sequester Boron in the cell wall and protect seagrasses against its toxic effects. No major changes could be observed for cellulose and hemi-cellulose biosynthesis (Extended Data Table 7). Interestingly, most of the evolutionary changes in seagrasses, linked to salt tolerance, are not mirrored in the genomes of mangrove species (*Avicinea marina* and *Rhizophora apiculate*) and is consistent with the independent evolution of salt tolerance in mangrove species ^50,51^.

#### Hypoxia

The solubility of oxygen in seawater is limited (typically around 10 mL O_2_ *L-1), while the sediments in which seagrasses grow are oxygen-free and reducing below a sediment depth of a few mm. This increases the O_2_ demand/draw-down by extensive root-rhizome tissues that are often >50% of total plant biomass. Consistent with the increased risk of hypoxia, all seagrasses have expanded their repertoire of Plant Cysteine Oxidases (PCO) and group VII Ethylene Responsive (ERF-VIIs) genes, for direct sensing and transcriptional adjustment to hypoxia (Figure 3a, 3d and Extended Data Table 8). As expected, most ERF-VIIs had higher expression in rhizomes and roots as compared to leaves (Suppl. Fig. S5.4.1). Also *P. acutifolius* contains an expanded hypoxia response machinery, reflecting its adaptation to submergence. Again, many, if not most, ERF-VII members reside within syntenic blocks retained from the WGT event in seagrasses, especially for *P. oceanica* and *T. testudinum* (Suppl. Fig. S5.4.2). Interestingly, this is also the case for multiple hypoxia-related genes. Some examples are: 1) the *PFK4* gene family, which contains the rate-limiting enzyme in the glycolysis pathway (including enolases), expanded in both seagrasses and *P. acutifolius*, and derived from the WGT event (Suppl. Fig. S5.4.2). 2) Lactate dehydrogenase, a rate limiting enzyme in lactate fermentation, is also expanded in seagrasses (Figure 3a and Extended Data Table 8) and has been shown to provide higher waterlogging tolerance in *Arabidopsis* upon overexpression ^52^; and 3) members encoding the energy-sensing sucrose nonfermenting kinase *SnRK1* ^53^ and *eIFiso4G1* (the dominant regulator in translational regulation by *SnRK1* under hypoxia ^54^ (Extended Data Table 8) are increased as a result of the WGT. In conclusion, it is interesting to speculate that the increase and specific retention of many hypoxia responsive genes, subsequent to the WGT (dated at ∼86 Mya), might have coincided with the Cenomanian-Turonian anoxic event (∼91± 8.6 Mya, ^55,56^), suggesting that this low oxygen period helped selecting for hypoxia tolerance in seagrasses. In *C. nodosa* and *P. acutifolius*, additional recent lineage specific WGDs and tandem duplications also contributed to further expansion of the hypoxia responsive genes as a potential adaptation to submergence.

#### Light perception and photosynthetic carbon acquisition

Seagrass growth and zonation are constrained by light availability, as ocean waters rapidly attenuate photosynthetic active radiation with depth and modify its spectral quality, enriching the blue wavelengths while reducing the red ^57^. Even in the clearest ocean waters, seagrasses mostly grow <40 m depth. Dissolved inorganic carbon (DIC) is mainly available as bicarbonate (HCO_3_^−^) in seawater (nearly 90% DIC at normal pH) that needs to be exploited via special acquisition systems, as it cannot diffuse passively across the cell plasma membrane ^58^. The availability of dissolved CO_2_ for photosynthesis is instead limited to ∼1% of the DIC pool, hence submerged plants and algae evolved CO_2_-concentrations mechanisms (CCMs) to overcome this low availability. A recent report identified an evolutionary adaptation of RuBisCO kinetics across submerged angiosperms from marine, brackish-water and freshwater environments that correlates with the development and effectiveness of CCMs ^59^.

The analysis of genes related to inorganic carbon (Ci) acquisition revealed a slight increase in extracellular *α-CA* (Carbonic Anhydrase α-type) copy number across the studied species (Suppl. Note S5.5.1). In *P. oceanica* and *P. acutifolius*, extra genes again have been specifically retained following the WGT event, although some copies have evolved through local tandem duplications as well. *α-CA* OG0013954 was found to be specific to seagrasses (except for *T. testudinum*) and *P. acutifolius* (Extended Data Table 9 and Suppl. Table S5.5), and most of the corresponding genes were highly expressed in leaves (Suppl. Fig. S5.5.1). This is consistent with their involvement in Ci acquisition and CCMs, as the presence of external Cas catalyzing the apoplastic dehydration of HCO3− to the RuBisCO substrate CO_2_, together with a higher activity of the extrusion proton pumps ^60^, is required for an adequate photosynthesis in most seagrass species ^61^. Furthermore, analysis of C4-pathway related genes revealed convergence, i.e., all genes are present in seagrasses (Extended Data Table 9).

The hypothesis that *C. nodosa* could be a C4 species ^62^ is here supported by the specific retention of 15 C4-related genes after WGT or WGD events, i.e., two encoding for C4 Phosphoenolpyruvate carboxylase (*PEPC*, retained after WGD) and, similar to what was observed in *P. acutifolius* (Extended Data Table 9). Notably, none of the studied seagrass species possesses the Ser residue characteristic of C4 *PEPC*, thus likely ruling out that a terrestrial-like C4-based (biochemical) CCM system is operating in seagrasses. This would suggest the presence of some kind of C3-C4 intermediate metabolism. Alternatively, homologs to C4 genes could have a role in the resistance of seagrasses to a variety of abiotic stresses, including salt stress ^63^.

Consistent with an augmented need for light capture, seagrasses showed an expansion of LHCB (light-harvesting complex B) as compared to freshwater plants that live in shallower environments (Extended Data Table 10 and Suppl. Note S5.5.2). Only *C. nodosa* showed the number of LHCB genes comparable to *P. acutifolius* and *Spirodela*. Other components of the photosynthetic machinery, including Photosystems I and II, were similar in number of genes to the ones of other species, either freshwater or terrestrial (Extended Data Table 10). Seagrasses conserved the full repertoire of orthologous genes for photosensory proteins and components of the light signaling systems (Extended Data Table 11 and Suppl. Note S5.5.3) that evolved in the green lineages during the different stages of plant terrestrialization ^64^.

UV-B tolerance and regulation of downstream signaling pathways vary among the seagrass species (Suppl. Note S5.5.3). Those living at lower latitudes (*T. testudinum* and *C. nodosa*), where UV-B radiation is expected to be intense and continuous throughout the year, kept the typical *UVR8* of land plants along with the main regulatory proteins (*RUP1,2*). *Z. marina*, instead, which occurs at higher latitudes and is exposed to lower level of UV-B radiation, lost the genes for both photoreceptors and their main negative regulatory proteins (Extended Data Table 11). In *P. oceanica*, a species restricted to the Mediterranean-climate region, the orthologous gene for *UVR8* lacks the sequence region C27 engaged in the regulation of UVR8 reversion state from the activated to the inactivated state (Yin et al. 2015). The species-specific adaptation in the UV-signaling and its negative feedback regulation (Suppl. Fig. S5.5.3), further reinforce the idea that ‘tweaking’ (and not massive change of key traits and their regulatory mechanisms) enabled the invasion of the marine environment.

Perception of surrounding light cues is also critical for the entrainment of the circadian clock system. This is essential for periodic regulation of physiology and the life cycle in plants, such as daily water and carbon availability, and hormone signaling pathways ^65^. All seagrass species, apart from *T. testudinum*, lost timing of CAB1 (*TOC1*) gene (Extended Data Table 12), one of the key clockwork components of the evening transcriptional-translational loop ^66,67^ belonging to the PSEUDO RESPONSE REGULATOR (PRR) family with a crucial function in the integration of light signals to the circadian control ^68^. Among its many entanglements, *TOC1* has a central role in adapting plant physiology to drought ^69,70^ and in regulating the day-night energy metabolism ^71^. Remarkably, *TOC1* is also lost in the freshwater *P. acutifolius* and *W. australiana*, the latter showing a reduced circadian time control of gene expression in comparison with *Arabidopsis* ^72^. The general reduction of clock genes in aquatic species suggests that the “absence of drought”, has led to a reduction of the regulatory daily-timing constraints for some metabolic and developmental plant processes. Interestingly, all seagrasses, possibly apart from *Z. marina*, retained some genes related to the phytochromes light-signaling pathway. These include PIFs and LAF1 (Extended Data Table 10) following WGT and WGD events, as well as genes related to the circadian clock and photoperiodism such as GI and ZTL (Extended Data Table 11).

#### NAC Transcription Factors

NAC transcription factors (TF) are one of the largest gene families. While a comparable number of sequences were found in seagrasses as compared with aquatics and land plants, specific orthogroups were different, in particular JUB1 genes, a central longevity regulator in addition to stress tolerance. Functional reorganization was observed in *C. nodosa* and *P. oceanica* (Suppl. Note S5.6 and Suppl. Fig. S5.6)

#### Nitrogen Metabolism

The key genes linked to nitrogen uptake/transport and assimilation have been retained in all seagrasses examined, while nitrate transporters (NRT) are strongly contracted (Extended Data Table 13, Suppl. Note S5.7 and Suppl. Table S5.7). This implies that seagrasses may have evolved alternative mechanisms for nitrogen uptake and utilization. Though our results are not particularly revealing, recent work on seagrass microbiomes has shown that nitrogen acquisition involves nitrogen-fixing bacteria in the roots ^73^ and that epiphytic micro-organisms on the leaves mineralize amino acids via their heterotrophic metabolism ^74^. Gaining a more mechanistic understanding of the plant role in these interactions, is now possible.

## CONCLUSION

Seagrass meadows are now recognized as invaluable ecosystems with multiple functions and services. They prevent erosion and hence preserve coastal seascapes, serve as biodiversity hotspots for micro- and megafauna, and have recently been proposed as a nature-based solution for climate mitigation owing to their carbon storage capacity in belowground biomass. Seagrasses also represent an extremely rare adaptation in the world of flowering plants, unlike adaptation to freshwater environments, which occurred hundreds or thousands of times. Only on three different occasions have seagrasses evolved from a freshwater ancestor to a (group of) species that lives continuously submerged in a highly saline environment, including subaqueous pollination (except the genus *Enhalus* ^75^).

Comparative genome analysis unveiled considerable convergence in seagrasses, but mainly for processes and pathways that have become redundant in a submerged marine environment. These include genes for stomata development, ethylene biosynthesis and signaling, disease resistance, and heat shock transcription factors (HSFs), which are involved in the rapid activation of stress-responsive genes and have been linked to the evolutionary adaptation of plants to the terrestrial environment. Jointly, these results illustrate that the invasion of the marine environment is associated with a significant loss of genes in multiple pathways that are no longer needed, a compelling example of “use it or lose it”.

Clear evidence of convergent positive (or gain of function) adaptation among the different lineages of seagrasses is harder to establish. Rather than unveiling major biological innovations or the rewiring of biological networks, adaptation to the marine environment seems to have mainly involved the fine-tuning of many different/supportive processes that likely all had to happen in parallel, possibly explaining why the transitioning to a marine lifestyle has been exceedingly rare. For instance, our analysis indicates that the adaptation of seagrasses to a marine (saline) environment was not accompanied by massive changes to individual salt tolerance traits, but rather involved more subtle changes in gene copy number and regulatory mechanisms. This gradual modulation of preexisting mechanisms is consistent with e.g., the presence of multiple halophytes within Alismatid families ^76^. The fine-tuning of many biological processes may also be at the basis of (some of the) plasticity displayed by seagrass populations and may have favoured their colonization of heterogeneous environments.

Many of the genes co-opted in different pathways in seagrasses seem to have been specifically retained following WGDs that occurred long ago, suggesting important interdependencies of large-scale (or major) genome evolution events and evolutionary adaptation. Prime examples identified here are hypoxia-responsive genes, genes involved in salt tolerance, flavonoid metabolism, carbon acquisition, and C4 photosynthesis. Therefore, the co-option of extra genes specifically retained following ancient whole genome duplications likely played a crucial role in facilitating survival in a marine environment.

We expect these new genomes to accelerate functional studies and contribute to transformative solutions in the management and conservation of seagrass ecosystems, as continuing worldwide losses of seagrass meadows are of major concern in times of climate change and loss of biodiversity.

## METHODS

### Sampling metadata, DNA and RNA preparation

Whole plants from each species were collected from the field, transported to the lab in a cool box, cleaned, frozen in LN2 and then stored at -80C. Collection and processing information are summarized in Suppl. Table S1.1. All samples were made with collection permits and followed the CBD-Nagoya Protocol. Care was taken to use tissue harvested from the basal area of young, clean leaves (10-cm pieces) to minimize epiphytic diatoms and bacteria If necessary. The seagrass tissues were then sent by overnight courier on dry ice to the Arizona Genomics Institute, Tucson, AZ, USA for extraction of nucleic acids (https://www.genome.arizona.edu). QC’d nucleic acid samples were then shipped on dry ice to the Joint Genome Institute (JGI) in Berkeley, CA, USA (https://jgi.doe.gov/) for further diagnostics and sequencing library preparation. For *P. acutifolius*, nucleic acids were extracted, QC’d and sequenced at the Max Planck-Genome-Centre Cologne, Germany (https://mpgc.mpipz.mpg.de/home/).

High Molecular Weight (HWM) DNA was extracted from young leaves of *T. testudinium, P. oceanica*, and *C. nodosa*, using the protocol of Doyle and Doyle (1987) with minor modifications. Young leaves, that had been flash frozen in LN2 and kept frozen at -80C, were ground to a fine powder in a frozen pestle and mortar with LN2 followed by very gentle extraction in CTAB buffer (that included proteinase K, PVP-40 and β-mercaptoethanol) for 20 mins at 37°C, followed by 20 mins at 50°C. Following centrifugation, the supernatant was gently extracted twice with 24:1 chloroform: iso-amyl alcohol. The upper phase was adjusted to 1/10^th^ volume with 3M Sodium acetate (pH=5.2), gently mixed, and DNA precipitated with iso-propanol. DNA was collected by centrifugation, washed with 70% EtOH, air dried for few minutes and dissolved thoroughly in 1x TE at room temperature. Size was validated by pulsed field electrophoresis. HMW DNA for *P. acutifolius* was extracted from 2 g of young leaves with the NucleoBond HMW DNA kit (Macherey Nagel). Quality was assessed with a FEMTOpulse device (Agilent) and the quantity was measured by a Quantus fluorometer (Promega).

RNA was extracted from seagrass leaves, rhizomes, roots, and flowers (Suppl. Table S1.1) with the NucleoSpin RNA Plant and Fungi Kit (Macherey-Nagel, USA), and checked for integrity by capillary electrophoresis using an Agilent (Santa Clara, CA, USA) 2100 Bioanalyzer with the Agilent RNA 6000 Nano Kit following manufacturer’s instructions. RNA was extracted from leaves and roots of *P. acutifolius* with the RNAeasys Plant Kit (Qiagen), including an on-column DNase I treatment. Quality was assessed with an Agilent Bioanalyser and the quantity was calculated by an RNA-specific kit from Quantus (Promega).

### Genome Sequencing

The genomes of *T. testudinium, P. oceanica*, and *C. nodosa* were determined following a whole genome shotgun sequencing strategy and standard sequencing protocols. Sequencing reads were produced using the Illumina NovaSeq platform and the PacBio SEQUEL II platform at the Department of Energy (DOE) Joint Genome Institute (JGI) in Walnut Creek, California, and the Hudson Alpha Institute in Huntsville, Alabama. One 400bp insert 2×150 Illumina fragment library and one HiC library was sequenced for each organism. Technical sequencing statistics are summarized in Suppl. Table S2.1.1. Prior to assembly, Illumina fragment reads were screened for Phix contamination and reads composed of >95% simple sequences were removed. Furthermore, Illumina reads <50bp, after trimming for adapter and checking for quality (q<20), were also removed. For the Illumina sequencing, the final combined read set consisted of 4,284,278,120 high-quality reads with 161x coverage for *T. testudinium*, 6,543,657,580 high-quality reads with 327x coverage for *P. oceanica*, and 693,903,610 high-quality reads with 208x coverage for C. nodosa. For the PacBio sequencing, a total of 18 PB chemistry 3.1 chips (30-hour movie time) were sequenced with a HiFi read yield of 231.8 Gb with 51.53x coverage, 238.3 Gb with 79.44x coverage and 39.6 Gb with 79.24x coverage for *T. testudinium, P. oceanica* and *C. nodosa*, respectively.

For *P. acutifolius*, all libraries (PacBio, RNA and Tell-seq) and PacBio HiFi sequencing were performed at the Max Planck-Genome-Centre Cologne, Germany (https://mpgc.mpipz.mpg.de/home/). Short-read libraries and sequencing (RNA-seq and Tell-seq) were performed at Novogene Ltd (UK), using a NovaSeq 6000 S4 flowcell Illumina system. An Illumina-compatible was prepared with the NEBNext® Ultra™ II RNA Library Prep Kit for Illumina. PacBio-HiFi libraries were prepared according to the manual “Procedure & Checklist - Preparing HiFi SMRTbell® Libraries using SMRTbell Express Template Prep Kit 2.0” with an initial DNA fragmentation by g-Tubes (Covaris) and final library size selection on BluePippin (Sage Science). Size distribution was again controlled by FEMTOpulse (Agilent). Size-selected libraries were sequenced on a Sequel II with Binding Kit 2.0 and Sequel II Sequencing Kit 2.0 for 30 h (Pacific Biosciences). The same genomic DNA was used for TELL-seq but without fragmentation. Library preparation was done as outlined in the manual “TELL-Seq™ WGS Library Prep User Guide” (ver. November 2020). Illumina “sequencing-by-synthesis” was performed on a HiSeq 2500, 2 × 250 bp with additional index sequencing cycles to read out the unique fragment barcodes. Sequences were analyzed as recommended by Universal Sequencing Technology (UST, Canton, U.S.A). The final combined read set consisted of 54,401,190 Illumina high-quality reads with 13.4 coverage and 1,900,000 PacBio HiFi reads with 43.5 coverage (Suppl. Table S2.1.1)

### Genome assembly and construction of pseudomolecule chromosomes

For *T. testudinium, P. oceanica* and *C. nodosa*, the following assembly strategy was used: the PacBio HiFi data was assembled using HiFiAsm and subsequently polished using RACON (https://github.com/lbcb-sci/racon). Due to the high heterozygosity of our sequenced seagrasses, both haplotypes were nearly complete resulting in a genome assembly composed of a highly contiguous primary set of chromosomes and a more fragmented alternative set of chromosomes (Suppl. Fig. S2.1.1). For *T. testudinium*, the initial primary assembly consisted of 1,987 contigs with a contig N50 of 483.4 Mb, and a total assembled size of 4,866.1 Mb. For *P. oceanica*, the initial primary assembly consisted of 3,470 contigs, with a contig N50 of 355.8 Mb, and a total assembled size of 3,192.0 Mb (Suppl Table S3). For C. nodosa, we produced an initial primary assembly of 1,362 contigs, with a contig N50 of 18.5 Mb, and a total assembled size of 466.0 Mb (Suppl. Table S2.1.2). Misjoins in the assemblies were identified using HiC data as part of the JUICER/JuiceBox pipeline^77^ for each of the three seagrass genomes. After resolving the misjoins, the broken contigs were then oriented, ordered, and joined together with HiC data using the JUICER/JuiceBox pipeline. In *T. testudinum*, there were 5 misjoins identified in the polished primary assembly, and a total of 15 joins were applied to the primary assembly to form the final assembly consisting of 9 chromosomes. In both the *P. oceanica* and *C. nodosa* polished primary genomes, there were no misjoins identified. A total of 6 joins were applied to the primary assemblies of *P. oceanica* and *C. nodosa* to form the final assembly consisting of 10 chromosomes and 18 chromosomes, respectively. Each chromosome join is padded with 10,000 Ns. Significant telomeric sequence was identified using the (TTTAGGG)_n_ repeat, and care was taken to make sure that contigs terminating in telomere were properly oriented in the production assembly. The remaining scaffolds were screened against bacterial proteins, organelle sequences, GenBank nr and removed if found to be a contaminant. Heterozygous SNP/indel phasing errors were corrected using the CCS data (51.53x for *T. testudinum*, 79.44x for *P. oceanica* and 79.24x for *C. nodosa*). Finally, homozygous SNPs and INDELs were corrected in the releases using Illumina reads (2×150, 400bp insert). A total of 2,613 homozygous SNPs and 82,421 homozygous INDELs were corrected in *T. testudinum*. A total of 1,643 homozygous SNPs and 100,570 homozygous INDELs were corrected in *P. oceanica* and total of 1,426 homozygous SNPs and 12,492 homozygous INDELs were corrected in the *C. nodosa*. Due to the high heterozygosity of the three genomes, both haplotypes of each chromosome were well represented in the assemblies. The primary set of chromosomes were constructed from the primary assembly, while an alternative set of chromosomes were constructed from the alternate assembly. Chromosomes for the alternate haplotype were then oriented, ordered, and joined together using synteny from the primary chromosomes (Suppl. Table S2.1.3).

For *Potamogeton acutifolius*, we used HiFiAsm ^78^ to assemble a draft genome assembly of a total length of 611 Mb with N50 = 3.09 Mb and scaffolded it further with Tell-seq data (linked reads; <u>bioRxiv 2019, 852947</u>) using the ARCS software ^79^ and reaching final N50 = 4.45 Mb (6,705 scaffolds in total, the length of the largest scaffold = 31.2 Mb). We assessed the completeness of gene space assembly using BUSCO - a set of conserved single-copy genes for Embryophyta, which resulted in 96.9% of complete genes, while 1.1% of genes from the set were fragmented and only 2% were missing: C:96.9% [S:93.7%, D:3.2%], F:1.1%, M:2.0%, n:1614.

### Genome annotation

#### Structural and functional annotation of genes

Our annotation pipeline integrated three independent approaches, the first one based on transcriptome data, the second one being an ab initio prediction and the third one based on protein homology. Both of RNA-seq and Iso-seq data from different tissues (Suppl. Table S3.2.1 – Suppl. Table S3.2.4) were used to aid the structural annotation and RNA-seq datasets were first mapped using Hisat2 (v2.1.0, arguments --dta) ^80^ and subsequently assembled into transcript sequences by Stringtie2 ^81^, whereas Iso-seq sequences were aligned to the seagrasses genome using GMAP ^82^. All transcripts from RNA-seq and Iso-seq were combined using Cuffcompare (v2.2.1) and subsequently merged with Stringtie2 (arguments --merge -m 150) to remove fragments and redundant structures ^81^. Transdecoder v5.0.2 (github.com/TransDecoder) was then used to predict protein sequences with diamond v2.0.14 results (--evalue 1e-5 --max-target-seqs 1 -f 6). BARKER v2.1.2 ^83^ was used for ab initio gene prediction using model training based on RNA-seq data. Homology-based annotation was based on the protein sequences from related species (*Z. marina* v1.0, *Spirodela polyrhiza, Oryza sativa* and *Arabidopsis thaliana*) as query sequences to search the reference genome using TBLASTN with e-value ≤1e^−5^, then regions mapped by these query sequences were subjected to Exonerate to generate putative transcripts. Additionally, an independent, homology-based gene annotation was performed using GeMoMa ^84^ using the same species with TBLASTN.

All structural gene annotations were joined with EvidenceModeller ^85^ v1.1.1, and BUSCO v4.0.4 (Benchmarking Universal Single-Copy Orthologs) ^86^ was used to assess the quality of the annotation results. Finally, we used GenomeView ^87^ to do the gene curations manually based on the RNA-seq and Iso-seq data. Putative gene functions were identified using InterProScan ^88^ with different databases, including PFAM, Gene3D, PANTHER, CDD, SUPERFAMILY, ProSite and GO. Meanwhile, functional annotation of these predicted genes was obtained by aligning the protein sequences of these genes against the sequences in public protein databases and the UniProt database using BLASTP with the e-value ≤1e – 5.

#### Annotation of non-protein coding RNA families (rRNA, snoRNAs, tRNAs and miRNAs)

The latest versions of the genome.fasta and genome.gff files for *Z. marina, C. nodosa, P. oceanica, T. testudinum* and *P. acutifolius* were downloaded from JGI ^89^ Infernal v1.1.4 (Dec 2020) ^90^ was used to perform sequence similarity searches of each genome sequence versus the RFAM database (RNA families database, Dec2021) ^91^. The output from Infernal was filtered keeping only the hits with an E-value threshold E<0.01. A second filtering step was performed to remove redundant information, i.e., overlapping matches with similar hits. A third filtering step was performed by retaining all the hits matching with a coverage of at least 95% and removing all partial/fragmented matches with incomplete hits from the reference collection. rRNA, tRNA, snoRNA and miRNA regions were selected and annotated in the annotation.jff files for each species. An updated functional annotation including the identified loci in the genomes was performed by scanning the Uniprot database ^92^ with BLASTp ^3^. Introns and the corresponding sequence regions were extracted by GenomeTools ^93^ and Bedtools ^94^ programs. The functional annotation of the long introns (>= 20kb) was performed by similarity searches in the NCBI nucleotide ^95^ database with the BLASTn tool ^3^.

#### Annotation of repeats and transposable elements (TEs)

Two complementary approaches were used to identify repetitive DNA sequences. First, a de novo repeat identification was carried out with RepeatModeler v2.0.1 (https://www.repeatmasker.org/RepeatModeler/) based on the default TE Rfam database, followed by RepeatMasker v4.1 (https://www.repeatmasker.org/) to discover and classify repeats based on the custom repeat libraries from RepeatModeler v2.0.1. Second, LTR_Finder ^96^ (v1.0.7), LTR_harvest ^97^ from genometools (v1.5.9) and LTR_retriever ^98^ (v2.9.0) were used to identify and trace the LTR elements, which are subsequently characterized at clade/lineage level by searching coding domains within the sequences, using the tool Domain based ANnotation of Transposable Elements (DANTE) (https://github.com/kavonrtep/dante). Transposable elements not classified by RepeatModeler were analyzed using DeepTE ^99^. We merged the libraries from RepeatModeler, LTR_retriever and DeepTE by USEARCH ^100^ with 80% identity as the minimum threshold for combining similar sequences into the final non-redundant de novo repeat library. Finally, we used RepeatMasker v4-1-0 (-e rmblast -gff -xsmall -s -norna -no_is -lib) to identify and classify repeats in the genome assemblies of seagrasses and Potamogeton.

#### Dating bursts of repeats in seagrass genomes

The identification of high-quality intact LTR-RTs and the calculation of insertion age for intact LTR-RTs were carried out using LTR_retriever (v2.9.0), using the formula T=K/2r. The nucleotide substitution rate “r” was set to 1.3e-8 substitutions per site per year.

### Identifying Whole Genome Duplications

#### KS age distributions, gene tree-species tree reconciliation, and absolute dating of WGDs

Ks age distribution analysis was performed using the wgd package ^101^. Anchor pairs (i.e., paralogous genes lying in collinear or syntenic regions of the genome) were obtained using i-ADHoRe ^102^. Ks distribution analysis was also performed using the KSRATES software ^103^, which locates ancient polyploidization events with respect to speciation events within a phylogeny, comparing paralog and ortholog K_S_ distributions, while correcting for substitution rate differences across the involved lineages (see Suppl. Note S4.2.1).

OrthoFinder ^104^ was used to build orthologous gene families. For each orthogroup, a multiply sequence alignment (MSA) based on amino acid sequences was obtained using PRANK ^105^ and then used as input for Markov Chain Monte Carlo (MCMC) analysis in mrbayes ^106^. A time-calibrated species tree was inferred by MCMCtree from the PAML package ^107^, using reference speciation times of 42–52 million years ago (MYA) for the divergence between *Oryzae sativa* and *Brachypodium distachyon*, 118-129 MYA for that between *Spirodela polyrhiza* and *Z. marina*, and 130-140 for that between *Spirodela* and other terrestrial monocots ^108^. A gene duplication-loss (DL)+WGD model, under critical and relaxed branch-specific rates, was implemented for the inference of the significance and corresponding retention rates of the assumed WGD events under Bayesian inference ^21^. (see Suppl. Note S4.2.2)

Absolute dating of WGD events was done as described previously for *Zostera marina* ^13^. Paralogous gene pairs located in duplicated segments (so called anchors) and duplicated pairs lying under the WGD peak (so-called peak-based duplicates) were collected for phylogenetic dating. Anchors, which are assumed to correspond to the most recent WGD, were detected using i-ADHoRe 3.0 ^102^. For each WGD paralogous pair, an orthogroup was created that included the two paralogues plus several orthologues from other plant species, as identified by InParanoid (v. 4.1) ^109^, using a broad taxonomic sampling. Gene duplicates were then dated using the BEAST v. 1.7 package ^110^ under an uncorrelated relaxed clock model with the LG+G (four rate categories) evolutionary model. A starting tree with branch lengths satisfying all fossil-prior-constraints was created according to the consensus APGIII phylogeny. Fossil calibrations were implemented using log-normal calibration priors (see Suppl. Note S4.2.3).

### Phylogenetic tree construction and estimation of divergence times

Protein sets were collected for 23 species (see Suppl. Note S4.3). These species were selected as representatives for monocots and eudicots, and representing different habitats from terrestrial, freshwater-floating, freshwater-submerged, to marine-submerged. Orthofinder v2.3 ^111^ was used to delineate gene families with mcl inflation factor 3.0. All-versus-all Diamond blast with an E-value cutoff of 1e−05 was performed and orthologous genes were clustered using OrthoFinder. Single-copy orthologous genes were extracted from the clustering results. MAFFT ^112^) with default parameters was used to perform multiple sequence alignment of protein sequences for each set of single-copy orthologous genes, and to transform the protein sequence alignments into codon alignments after removing the poorly aligned or divergent regions using trimAl ^113^. The resulting codon alignments from all single copy orthologs were then concatenated into one supergene for species phylogenetic analysis. A maximum-likelihood phylogenetic tree of single-copy protein alignments and codon alignments was constructed using IQ-TREE ^114^ with the GTR+G model and 1,000 bootstrap replicates. Divergence times between species were estimated using MCMCtree from the PAML package under the GTR+G model (see Suppl. Note S4.3).

### Gene family comparisons

Gene families analyzed in the paper were searched in the output from Orthofinder and a big table was compiled to show the detailed information for each orthogroup, which is defined as the group of genes from multiple species descended from a single gene in the last common ancestor. For the superfamilies, we used the phylogenetic tree to further classify them into subfamilies. We adopted a custom criterion to assess the expansion and contraction of gene families. If the average gene number in seagrasses increased or reduced by >40% compared to non-seagrass species, we called it expansion or contraction. Syntenic analysis of genes are performed using MCScanX ^115^ and i-ADHoRe ^102^. Lastly, circos plots were drawn using Circos ^116^.

## ACKNOWLEDGEMENTS

Y.VdP., J.O., T.R. and G.P. acknowledge funding from the US-Dept. of Energy, Joint Genome Institute, Berkeley, California, USA, under the Community Sequencing Program 2018, Project Number 504341 (Marine Angiosperm Genomes Initiative-MAGI). The CSP award also included support sequencing and plant bioinformatics from HudsonAlpha Institute for Biotechnology, Huntsville, AL; and DNA/RNA extraction and processing from the Arizona Genomics Institute, Tuscon, AZ. Y.VdP. acknowledges funding from the European Research Council (ERC) under the European Union’s Horizon 2020 research and innovation program (No. 833522) and from Ghent University (Methusalem funding, BOF.MET.2021.0005.01). P.N. acknowledges funding by the Deutsche Forschungsgemeinschaft (DFG, German Research Foundation) – project number 497665889, 1606/3-1 for research on *Potamogeton*. M.K. acknowledge funding through the Helmholtz School for Marine Data Science (MarDATA), Grant No. HIDSS-0005. The work of G.P., E.D., J.P., and M.R. was partially supported by the project Marine Hazard, PON03PE_00203_1 (MUR, Italian Ministry of University and Research). M.D.D., L.L.W., M.P.T. and Y.Y.S. acknowledge funding from Universiti Malaysia Terengganu (SRG Vot55317). The work (proposal: 10.46936/10.25585/60001196) conducted by the U.S. Department of Energy Joint Genome Institute (https://ror.org/04xm1d337), a DOE Office of Science User Facility, is supported by the Office of Science of the U.S. Department of Energy operated under Contract No. DE-AC02-05CH11231.

## Author contributions table

X.M., V.S. and J.Ch. contributed equally to the work and are joint first authors. Y.VdP., J.O., T.R. and G.P. conceived the project, provided the overall evolutionary context, and wrote the proposal. Y.VdP., J.O., T.R. and G.P. and S.V. wrote and edited the main manuscript, and organized and further edited the individual contributions for the Supplementary Notes (as listed for the Supplementary Information below). All co-authors then provided specific feedback in forming the final version.

J.O., J.Ca., G.P., L.M.G., T.R., A.B., A.M., and P.N. contributed to sample tissue collection, preparation, and shipping for DNA extraction. S.Raj. performed the HMW DNA extractions and QC, as well as RNA extractions and QC for annotation assistance. J.S. and J.G. coordinated genome sequencing management steps for the seagrasses. A.M. and P.N. coordinated genome sequencing management steps for *Potamogeton*. K.B. was responsible for overall JGI technical coordination, liaison with principal investigators and project manager.

J.J., C.P., J.S., Y.VdP., S.R., A.S., J.vV. and T.B. performed analysis activities surrounding genome assembly (PacBio, HiC), supporting transcriptomics for annotation. J.J., Y.VdP., S.R., X.M. were responsible for deposition and maintenance of the species on the ORCAE site, and deposition of the new genomes to NCBI and Phytozome. J.Ch., X.M.,S.M., J.Ch. were responsible for manuscript graphics.

Analysis of architectural features of genome evolution and annotation of specific gene families, including the written contributions to the main paper and Supplementary Information sections as follows: M.L.C., L.A. for the Orthogroups Master Extended Data; M.L.C., A.S., X.M., JCh. for overview of gene families; H.C., X.M., J.Ch., Y.VdP. for Whole Genome Duplications/Triplications and dating; M.L.C., A.S. for Transposable Elements and repeat elements; M.K., T.R. for Organellar genomes; M.L.C., L.A. for Non-protein coding RNA families; S.V., X.M. for Stomata; S.V., X.M. for Volatile metabolites and signaling, ethylene; S.V. for Plant body development, lignification, vascular tissue; T.B. for Plant defense, R-genes; L.A. for Heat shock factors; S.V. for Flavenoids and phenolics; S.V. for Cellular salt tolerance; S.V., B.V. for Cell wall plasticity; S.V. for Hypoxia; G.P. for Light perception, photosynthesis, light harvesting, transcription factors; G.P., M.R. Carbon acquisition, CCMs; G.P. for UVB tolerance; G.P., E.D. for Clock genes; J.P. for NAC genes; D.M., L.W., M.P.T., Y.Y.S. for Nitrogen metabolism.

