## Supplementary_Information for "Seagrass genomes reveal a hexaploid ancestry facilitating adaptation to the marine environment"

**Supplementary Fig. S4.1** Insertion time distributions of LTR/Gypsy and LTR/Copia in *T. testudinum*, *P. oceanica*, *C. nodosa* and *Z. marina*. Note differences in the scale of the y-axis. See main text for details.	24

**Supplementary Fig. S4.2.1** *K*_S_ distributions for anchor pair duplicates (duplicates laying in duplicated, colinear blocks) and the whole paranome of four seagrasses, as well as for *P. acutifolius* and *S. polyrhiza*, generated by the wgd software (see Methods)*.*	26

**Supplementary Fig. S4.2.2** Comparison of *Z. marina*, *P. oceanica*, and *T. testudinum* with *Aristolochia fimbriata*.	27

**Supplementary Fig. S4.2.3** Comparison of *P. oceanica* and *Aristolochia fimbriata* with *C. nodosa*.	28

**Supplementary Fig. S4.2.4** Comparison of *Aristolochia fimbriata*, *Z. marina, P. oceanica* and *C. nodosa* with *P. acutifolius.*	29

**Supplementary Fig. S4.2.5** *K*_S_ Distributions for paralogs and the whole paranome of four seagrasses and *P. acutifolius* generated by KSRATES software.	30

**Supplementary Fig. S4.2.6** Bayesian inference of duplication, loss, and retention rates in WHALE analysis.	32

**Supplementary Fig. S4.2.7a** Estimation of the 'absolute age’ of the *T. testudinum* WGT event by phylogenomic dating of *T. testudinum* paralogues.	33

**Supplementary Fig. S4.2.7b** Estimation of the 'absolute age’ of the *P. oceanica* WGT event by phylogenomic dating of *P. oceanica* paralogues.	33

**Supplementary Fig. S4.2.7c** Estimation of the 'absolute age’ of the most recent *C. nodosa* WGD event, by phylogenomic dating of *C. nodosa* paralogues.	33

**Supplementary Fig. S5.4.2** Syntenic relationship of genes mentioned in the main text for *P. oceanica*, *T. testudinum*, *Z. marina*, *C. nodosa*, and *P. acutifolius*.	46

**Supplementary Fig. S5.6** Evolutionary analysis by Maximum Likelihood method of *JUB1* in seagrasses.	56

### 1. Sampling species, metadata, and DNA and RNA preparation

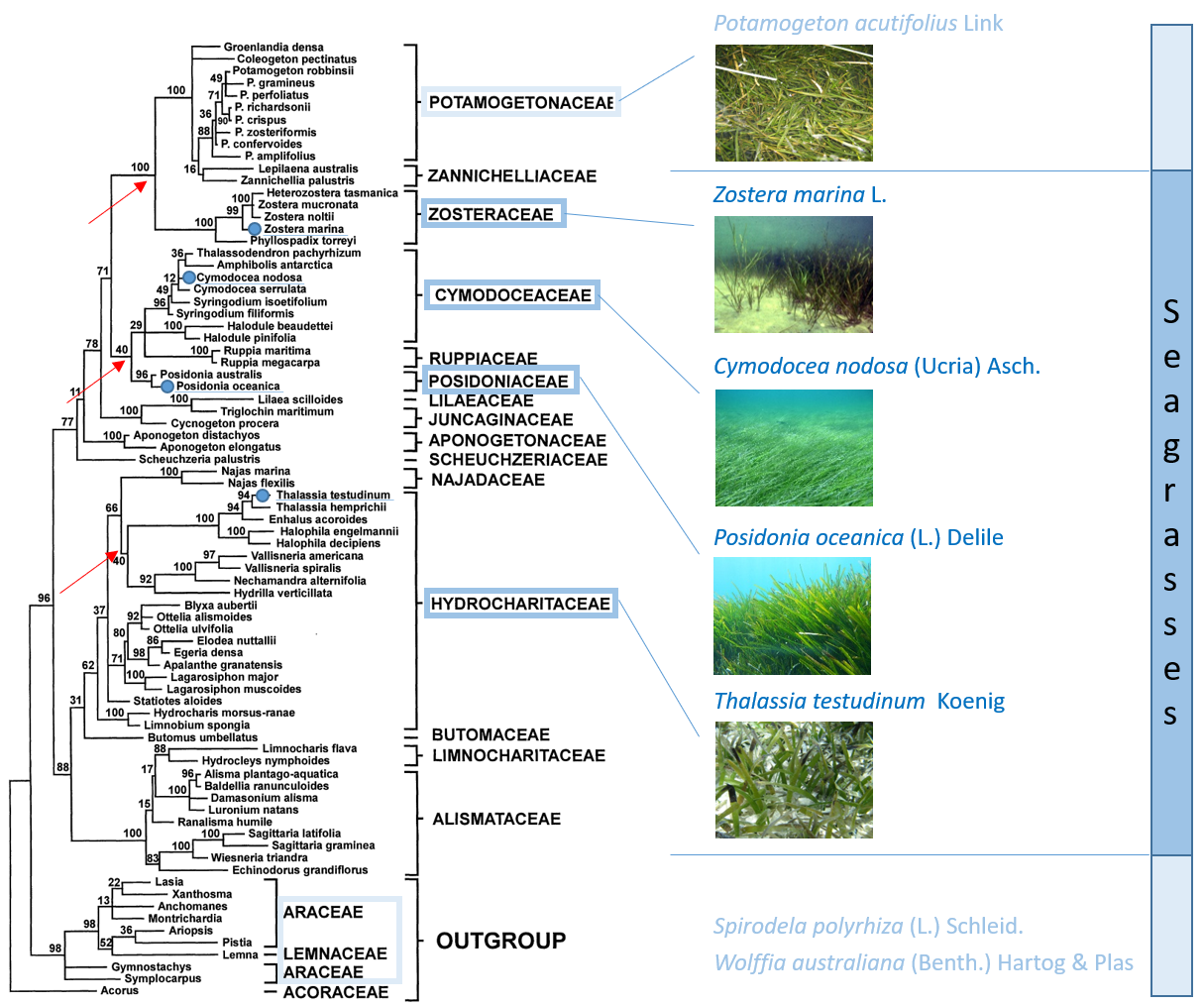

##### **Supplementary Fig. S1.1** Phylogenetic tree of Alismatales, including seagrasses.

The tree is based on a Maximum Parsimony analysis of *rbcL* genes (adapted from (Les et al. 1997)). The four seagrass species discussed in the present study are marked by blue dots. The one freshwater species sequenced in the present study, *Potamogeton acutifolius*, is missing from the tree, but close relatives are included (sister group of Zosteraceae). Solid red arrows indicate nodes denoting the aquatic-marine splits.

##### **Supplementary Table S1.1** Plant material metadata.

See Materials and Methods for details on DNA and RNA preparation. Additional information for *Zostera marina* can be found in (Olsen et al. 2016) and (Ma et al. 2021).

| **Species** | **Code** | **Location** | **Date** | **Collectors** | **Depth** | **Lat** | **Long** | **Tissue** | **HMW DNA extraction method**  **/RNA extraction method** |
| --- | --- | --- | --- | --- | --- | --- | --- | --- | --- |
| *Thalassia testunidum*  Diploid  2N=18 | Tt-101 | Coconut Grove Dog Park, Crocodile Point, Miami, FL USA | 12 June 2019 | JL Olsen &  JE Campbell | -1 m | 40.78735 | 14.08974 | Leaf  Leaf  Rhizome  No Root | CTAB via  Arizona Genomics Institute.  /NucleoSpin RNA Plant and Fungi Kit (Machenery Nagel, USA) |
| *Posidonia oceanica*  Diploid  2N=20 | Po-2 | Gulf of Pozzuoli, Napoli, Italy | 15 May 2019 | G Procaccini | -7 m | 40.78735 | 14.08974 | Leaf  Leaf  Rhizome Root | As above  As above |
| *Cymodocea nodosa*  Diploid  2N=18  Known polyploidy in some populations | Cn-1 | Miseno Cape, Napoli, Italy | 15 May 2019 | G Procaccini & L Marin Guirao | -10 m | 40.78396 | 14.07458 | Leaf  Leaf Rhizome  Root Flowers | As above  As above |
| *Potamogeton acutifolius*  Diploid  2N=26  Known polyploidy in other species  Freshwater | Pa | Baláta-lake, Hungary, Somogy county, near Kaszó | 18 Sept 2020 | A. Mesterházy | -1.2 m | 14.31388 | 17.205 | Leaf  Leaf  Turion  Root | Max Planck-Genome  with a NucleoBond HMW DNA kit (Macherey Nagel).  RNAeasy Plant Kit (Qiagen) |

### 2. Genome sequencing and assemblies

#### 2.1 Nuclear genomes

##### **Supplementary Table S2.1.1** Genomic libraries included in each seagrass genome assembly and their respective assembled sequence coverage levels in the final release.

|  | Library | Sequencing Platform | Average Read/Insert Size | Read Number | Assembled Sequence Coverage (x) |
| --- | --- | --- | --- | --- | --- |
| *Thalassia testudinium* | JDRM | Illumina | 400 | 2,454,318,746 | 92.04 |
|  | IYTB | Illumina/HiC | N/A | 1,829,959,374 | 68.62 |
|  | Total Illumina |  |  | 4,284,278,120 | 161 |
|  |  | CCS PacBio | 18,712* | 11,884,855 | 51.53 |
| *Posidonia oceanica* | JDRD | Illumina | 400 | 2,700,014,314 | 135 |
|  | JDMC | Illumina/HiC | N/A | 3,843,643,266 | 192.1 |
|  | Total Illumina |  |  | 6,543,657,580 | 327 |
|  |  | CCS PacBio | 19,006* | 12,018,008 | 79.44 |
| *Cymodocea nodosa* | JLSQ | Illumina | 400 | 397,732,894 | 119.3 |
|  | JHNL | Illumina/HiC | N/A | 296,170,716 | 88.9 |
|  | Total Illumina |  |  | 693,903,610 | 208 |
|  |  | CCS PacBio | 23,472* | 1,698,568 | 79.24 |
| *Potamogeton acutifolius* | N/A | Illumina/Tell-Seq | N/A | 54,401,190 | 13.4 |
|  | N/A | PacBio HiFi | 14,000* | 1,900,000 | 43.5 |

*Average read length of PacBio reads

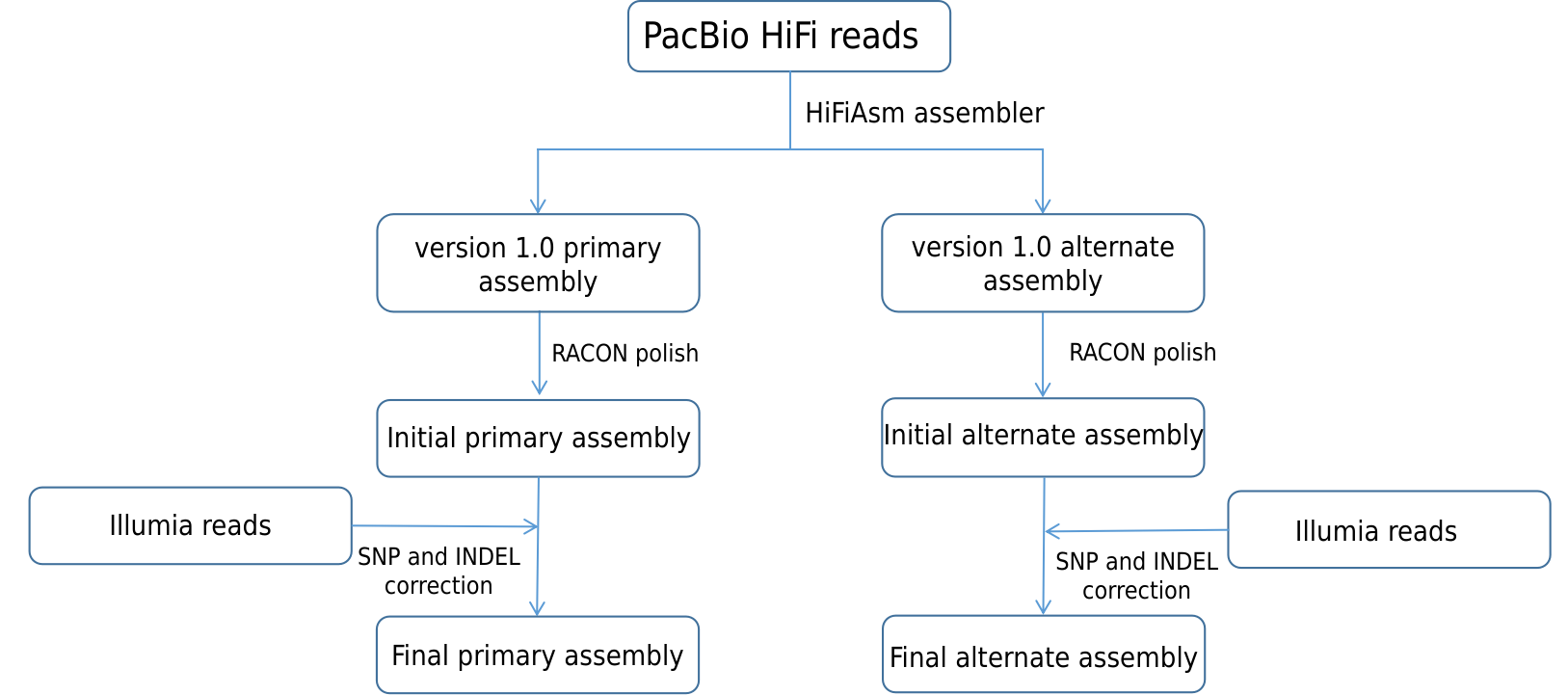

##### **Supplementary Fig. S2.1.1** Genome assembly pipeline used for *T. testudinum*, *P. oceanica* and *C. nodosa*

##### **Supplementary Table S2.1.2** Summary statistics of the initial output of the primary RACON polished HiFiAsm assembly.

The table shows number of contigs and total numbers of assembled base pairs for each set of scaffolds greater than the size listed in the left-hand column.

|  | **Minimum Scaffold Length** | **Number of Scaffolds** | **Number of Contigs** | **Scaffold Size** | **Basepairs** | **% Non-gap Basepairs** |
| --- | --- | --- | --- | --- | --- | --- |
| *Thalassia testudinium* | 5 Mb | 14 | 14 | 4,757,383,018 | 4,757,383,018 | 100.00% |
|  | 2.5 Mb | 14 | 14 | 4,757,383,018 | 4,757,383,018 | 100.00% |
|  | 1 Mb | 17 | 17 | 4,761,201,586 | 4,761,201,586 | 100.00% |
|  | 500 Kb | 21 | 21 | 4,764,120,924 | 4,764,120,924 | 100.00% |
|  | 250 Kb | 29 | 29 | 4,766,780,278 | 4,766,780,278 | 100.00% |
|  | 100 Kb | 152 | 152 | 4,784,190,954 | 4,784,190,954 | 100.00% |
|  | 50 Kb | 609 | 609 | 4,813,980,144 | 4,813,980,144 | 100.00% |
|  | 25 Kb | 1,965 | 1,965 | 4,865,616,055 | 4,865,616,055 | 100.00% |
|  | 10 Kb | 1,987 | 1,987 | 4,866,121,113 | 4,866,121,113 | 100.00% |
|  | 5 Kb | 1,987 | 1,987 | 4,866,121,113 | 4,866,121,113 | 100.00% |
|  | 2.5 Kb | 1,987 | 1,987 | 4,866,121,113 | 4,866,121,113 | 100.00% |
|  | 1 Kb | 1,987 | 1,987 | 4,866,121,113 | 4,866,121,113 | 100.00% |
|  | 0 bp | 1,987 | 1,987 | 4,866,121,113 | 4,866,121,113 | 100.00% |
| *Posidonia oceanica* | 5 Mb | 15 | 15 | 3,003,306,251 | 3,003,306,251 | 100.00% |
|  | 2.5 Mb | 15 | 15 | 3,003,306,251 | 3,003,306,251 | 100.00% |
|  | 1 Mb | 21 | 21 | 3,012,259,886 | 3,012,259,886 | 100.00% |
|  | 500 Kb | 29 | 29 | 3,017,253,515 | 3,017,253,515 | 100.00% |
|  | 250 Kb | 52 | 52 | 3,024,690,979 | 3,024,690,979 | 100.00% |
|  | 100 Kb | 201 | 201 | 3,045,962,177 | 3,045,962,177 | 100.00% |
|  | 50 Kb | 1,061 | 1,061 | 3,100,982,328 | 3,100,982,328 | 100.00% |
|  | 25 Kb | 3,383 | 3,383 | 3,190,011,804 | 3,190,011,804 | 100.00% |
|  | 10 Kb | 3,468 | 3,468 | 3,191,950,648 | 3,191,950,648 | 100.00% |
|  | 5 Kb | 3,468 | 3,468 | 3,191,950,648 | 3,191,950,648 | 100.00% |
|  | 2.5 Kb | 3,468 | 3,468 | 3,191,950,648 | 3,191,950,648 | 100.00% |
|  | 1 Kb | 3,470 | 3,470 | 3,191,952,950 | 3,191,952,950 | 100.00% |
|  | 0 bp | 3,470 | 3,470 | 3,191,952,950 | 3,191,952,950 | 100.00% |
| *Cymodocea nodosa* | 5 Mb | 21 | 21 | 370,838,187 | 370,838,187 | 100.00% |
|  | 2.5 Mb | 22 | 22 | 375,803,086 | 375,803,086 | 100.00% |
|  | 1 Mb | 26 | 26 | 381,781,051 | 381,781,051 | 100.00% |
|  | 500 Kb | 27 | 27 | 382,524,700 | 382,524,700 | 100.00% |
|  | 250 Kb | 46 | 46 | 388,996,552 | 388,996,552 | 100.00% |
|  | 100 Kb | 148 | 148 | 403,137,124 | 403,137,124 | 100.00% |
|  | 50 Kb | 678 | 678 | 437,596,903 | 437,596,903 | 100.00% |
|  | 25 Kb | 1,362 | 1,362 | 465,959,990 | 465,959,990 | 100.00% |
|  | 10 Kb | 1,362 | 1,362 | 465,959,990 | 465,959,990 | 100.00% |
|  | 5 Kb | 1,362 | 1,362 | 465,959,990 | 465,959,990 | 100.00% |
|  | 2.5 Kb | 1,362 | 1,362 | 465,959,990 | 465,959,990 | 100.00% |
|  | 1 Kb | 1,362 | 1,362 | 465,959,990 | 465,959,990 | 100.00% |
|  | 0 bp | 1,362 | 1,362 | 465,959,990 | 465,959,990 | 100.00% |

##### **Supplementary Note S2.1** Final Primary assemblies for main and alternate haplotypes.

The final primary assembly (see Methods) of *T. testudinum* contains 4,261.9 Mb of sequence, consisting of 184 contigs with a contig N50 of 371.6 Mb and a total of 98.92% of assembled bases in 9 chromosomes. The final primary assembly of *P. oceanica* contains 2,963.0 Mb of sequence, consisting of 19 contigs with a contig N50 of 355.8 Mb and a total of 99.98% of assembled bases in 10 chromosomes. The final primary assembly of *C. nodosa* contains 379.5 Mb of sequence, consisting of 22 contigs with a contig N50 of 21.9 Mb and a total of 100% of assembled bases in 18 chromosomes (Supplementary Table S2.1.3).

Correspondingly, the final alternative release of *T. testudinum* contains 4,177.2 Mb of sequence, consisting of 3,364 contigs with a contig N50 of 6.1 Mb and a total of 85.18% of assembled bases in 9 chromosomes. The final alternative release of *P. oceanica* contains 2,496.3 Mb of sequence, consisting of 826 contigs with a contig N50 of 8.7 Mb and a total of 99.67% of assembled bases in 10 chromosomes. The final alternative assembly for *C. nodosa* contains 374.7 Mb of sequence, consisting of 28 contigs with a contig N50 of 20.8 Mb and a total of 99.91% of assembled bases in 18 chromosomes (Supplementary Table S2.1.3). The final assembly of *P. acutifolius* is in the Table 1.

##### **Supplementary Table S2.1.3** Final summary primary and alternate assembly statistics for each chromosome-scale assembly. (see (Ma et al. 2021) for further details on *Zostera marina* v3.1.

| Final Primary Assembly Release Haplotype 1 | genome release stats: | *Thalassia testudinium* | *Posidonia oceanica* | *Cymodocea nodosa* | *Zostera marina v3.1* |
| --- | --- | --- | --- | --- | --- |
|  | Number of chromosomes (1N) | 9 | 10 | 18 | 6 |
|  | Number of main genome scaffold total: | 169 | 13 | 18 | 310 |
|  | Number of main genome contig total: | 184 | 19 | 22 | 432 |
|  | Length of main genome scaffold sequence total: (MB) | 4262.1 | 2963.1 | 379.6 | 260.5 |
|  | Length of main genome contig sequence total: | 4261.9 | 2963 | 379.5 | 259.3 |
|  | Main genome scaffold L/N50: (MB) | 4/523.9 | 4/355.8 | 8/22.6 | 4/34.6 |
|  | Main genome contig L/N50: (MB) | 5/371.6 | 4/355.8 | 8/21.9 | 12/7.0 |
|  | Number of scaffolds > 50 KB: | 49 | 13 | 18 | 217 |
|  | % main genome in scaffolds > 50 KB: | 99.9 | 100 | 100 | 98.9 |
| Final Alternative Assembly Release Haplotype 2 | Number of chromosomes (1N) | 9 | 10 | 18 | N/A |
|  | Number of alternate genome scaffold total: | 2264 | 29 | 22 | N/A |
|  | Number of alternate genome contig total: | 3364 | 826 | 28 | N/A |
|  | Length of alternate genome scaffold sequence total: (MB) | 4188.2 | 2504.3 | 374.8 | N/A |
|  | Length of alternate genome contig sequence total: (MB) | 4177.2 | 2496.3 | 374.7 | N/A |
|  | Genome scaffold L/N50: (MB) | 5 / 458.8 Mb | 3/352.0 | 8/22.1 | N/A |
|  | Genome contig L/N50:   (MB) | 162 / 6.1 Mb | 85/8.7 | 8/20.8 | N/A |
|  | Number of scaffolds > 50 KB | 805 | 29 | 22 | N/A |
|  | % main genome in scaffolds > 50 KB | 98.7 | 100 | 100 | N/A |

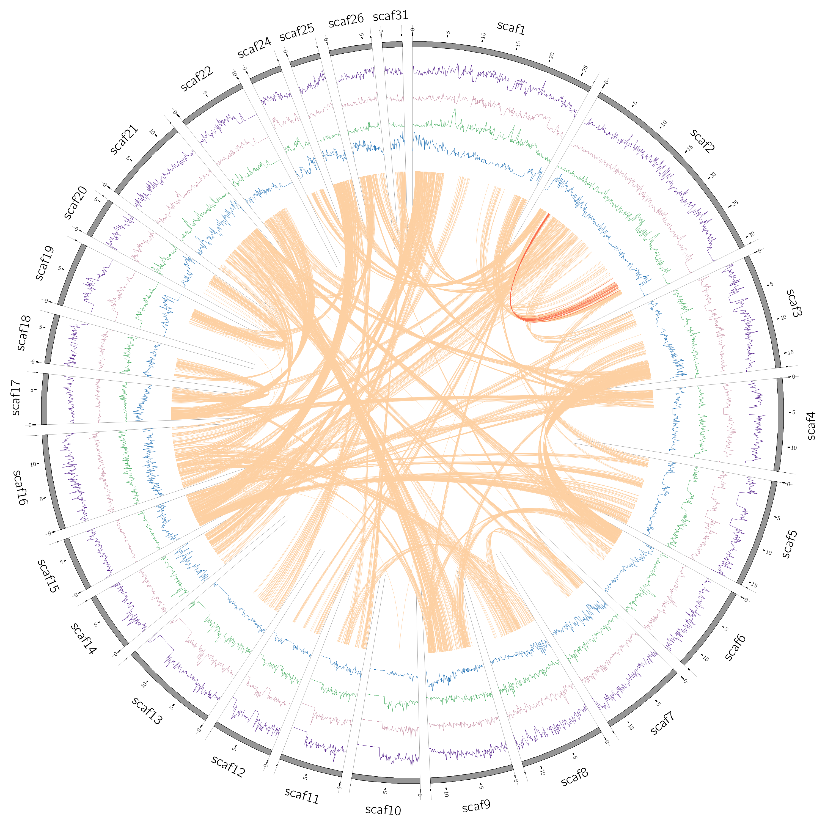

##### **Supplementary Fig. S2.1.2** Distribution of the genomic features for the 26 largest scaffolds of *P. acutifolius*.

Tracks from the inner to outer side correspond to gene density (blue); LTR/Gypsy density (green); LTR/Copia (orange); DNA transposable elements (pink) and chromosomes (with length in Mb). Curved lines through the center denote synteny between different scaffolds.

#### 2.2 Chloroplast genomes

##### **Supplementary Note S2.2** Chloroplast genome assemblies and annotations

Complete chloroplast genomes were assembled *de novo* from Illumina short read data with NOVOPlasty using the *rbcL* gene as a seed (Dierckxsens et al. 2017). *T. testudinum* cpDNA was additionally, manually curated according to the chloroplast derived PacBio contigs from the main genome assembly. The *Z. marina* genome was obtained from (Yu et al. 2022). All chloroplast genomes were polished with pilon (Walker et al. 2014) in addition to manual curation. The gene content of the chloroplast genomes was annotated using the software GeSeq (Tillich et al. 2017) and with Chloë (Zhong 2020). As the preferred annotator for CDS and rRNA, ARAGORN (Laslett and Canback 2004) was used for tRNA annotation. Genes that were much shorter than expected are marked as fragmented. Figures were drawn with Chloroplot (Zheng et al. 2020).

We identified 79, 78, 75, and 74 protein-coding genes (only counting genes in the inverted repeats once) in the *C. nodosa*, *Z. marina*, *T. testudinum*, and *P. oceanica* chloroplast genomes, respectively, of which *Z. marina* lost one gene (*rps19*), *T. testudinum* lost four genes (*accD, infA, ndhB,* and *ndhF*) and *P. oceanica* lost five genes (*ndhG, ndhH, ndhI, ndhJ,* and *ndhK*) (Suppl. Fig. S2.2.1 – Suppl. Fig. S2.2.4). We note that the chloroplast NADH dehydrogenase complex, encoded by *ndh* genes, has been lost in *P. oceanica* and *T. testudinum*. The loss has been proposed to have happened independently in various Alismatales lineages, correlating with the submerged fresh or marine habitat (Ross et al. 2016). Notably, the complex is lost in the two species that demonstrate the highest levels of genome perturbation.

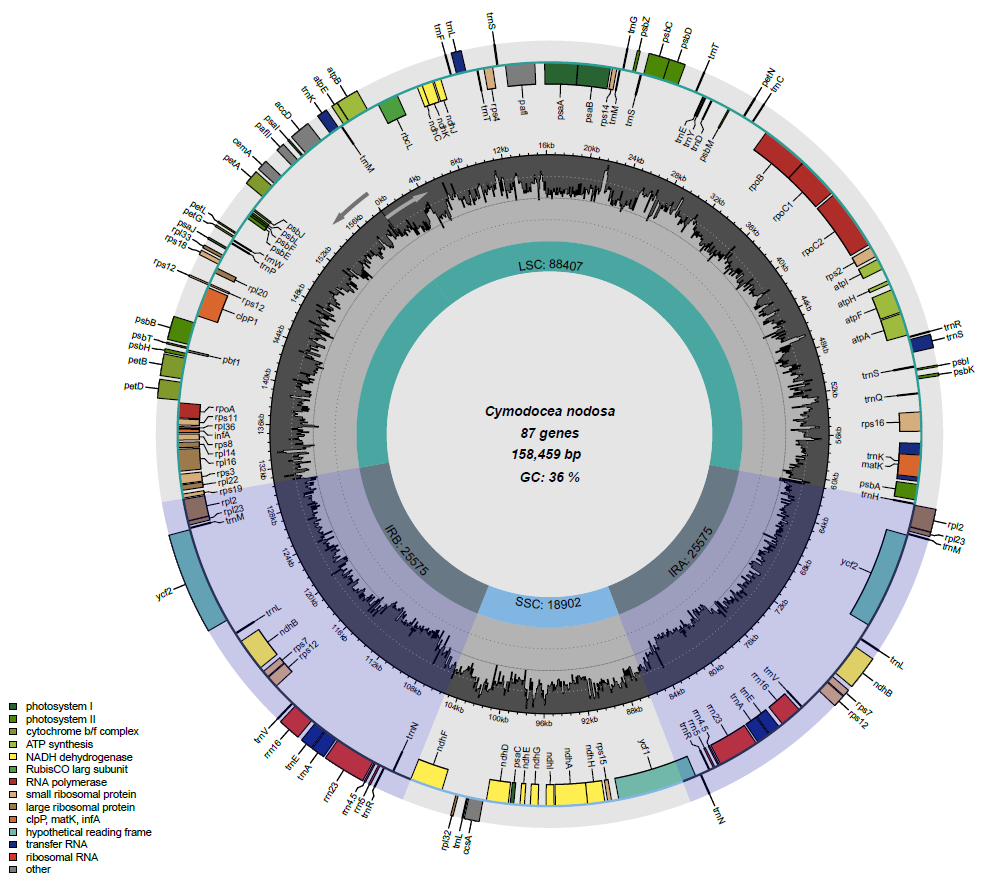

##### **Supplementary Fig. S2.2.1** The complete *C. nodosa* chloroplast genome.

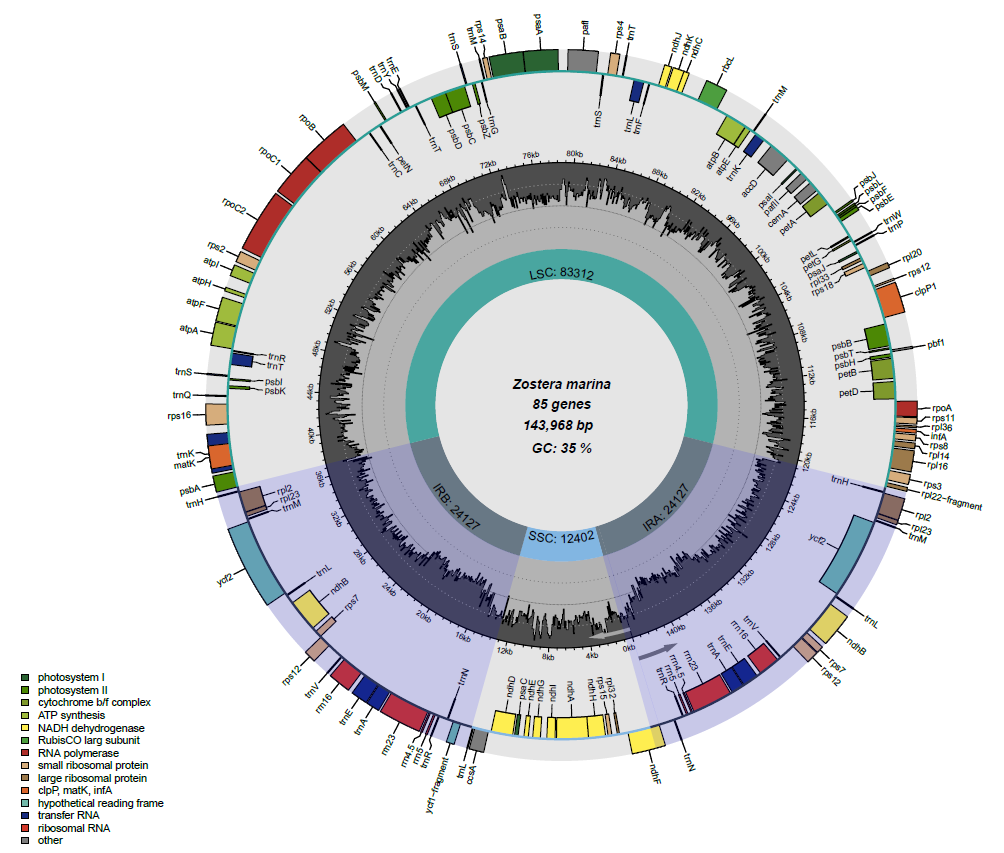

##### **Supplementary Fig. S2.2.2** The complete *Z. marina* chloroplast genome.

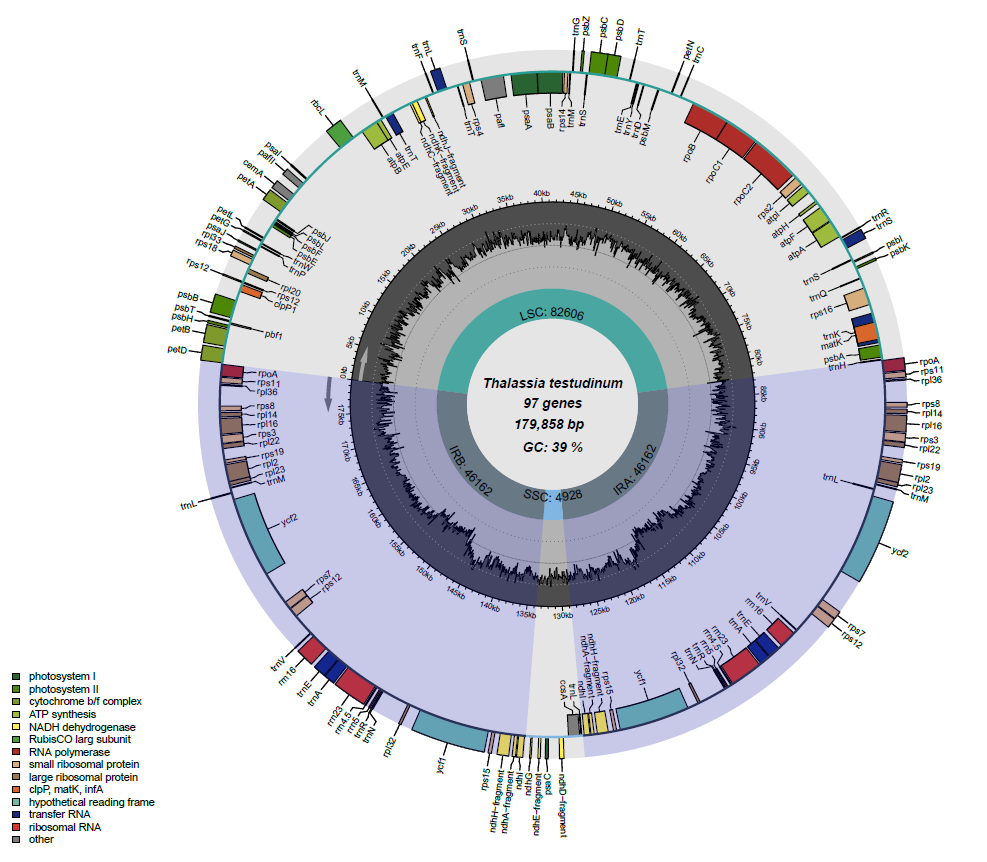

##### **Supplementary Fig. S2.2.3** The complete *T. testudinum* chloroplast genome.

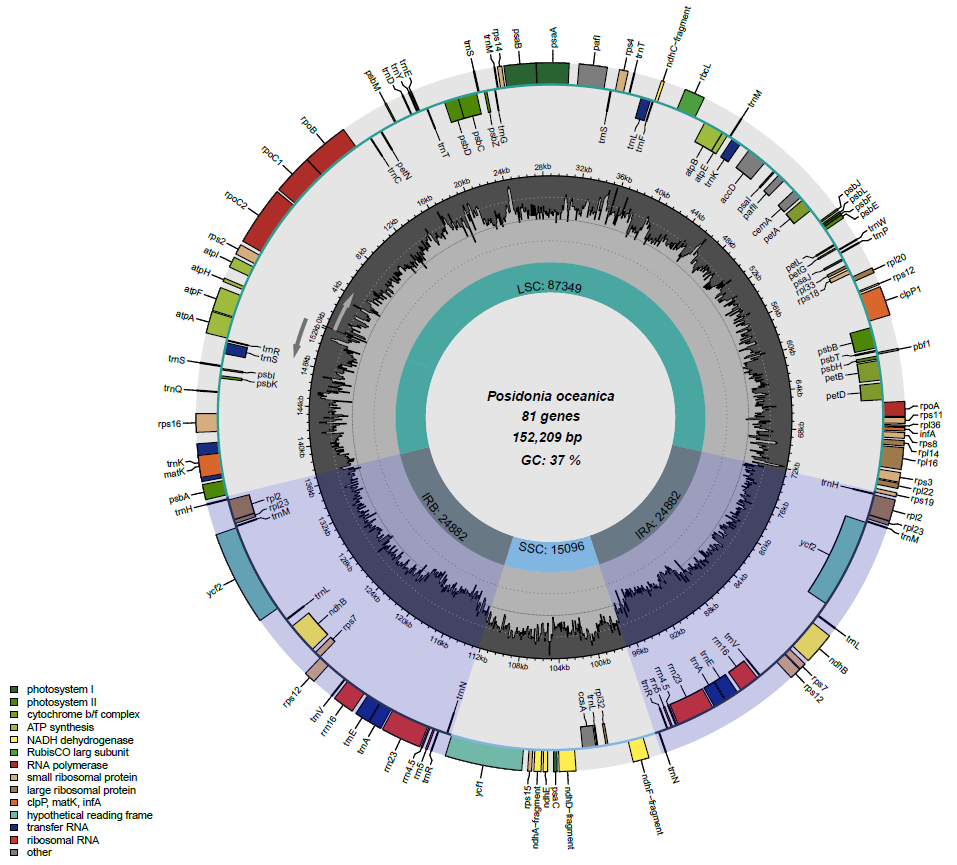

##### **Supplementary Fig. S2.2.4** The complete *P. oceanica* chloroplast genome.

#### 2.3 Mitochondrial genomes

##### **Supplementary Note S2.3** Mitochondrial genome assemblies and annotations

Mitochondrial genomes were manually assembled de novo from PacBio contigs containing at least five mitochondrial genes. All mitochondrial genomes were polished with pilon. Genes were identified by BLAST and TBLASTN (Altschul et al. 1990) using the mitochondrial gene collection in (Petersen et al. 2017).

Unlike animal mtDNA, plant mtDNA genomes vary enormously in size and do not exist as a single stable circle but rather as a dynamic combination of linear, branched and small circular loops called isoforms (Morley and Nielsen 2017). The additional DNA is dominated by repeats and noncoding regions. The various isoforms are due to recombination (Kozik et al. 2019). Therefore, the classic circular representation is inappropriate. Accordingly, we found varying degrees of genome size, completeness, and fragmentation in the seagrass mitochondrial genomes.

The *Z. marina* mitochondrial genome is complete (Khachaturyan et al., submitted) as is *C. nodosa*. *T. testudinum* is incomplete and only a single mt-contig was recovered for *P. oceanica*.

The complete assembly of the *C. nodosa* mitochondrial genome resulted in one circular chromosome of 293,083 bp with two recombinationally active direct repeats of 2,683 bp and 2,476 bp. Given the possible recombination events, the genome has six chromosomes (Suppl. Fig. S2.3) All PacBio assembly contigs containing fragments of five or more mitochondrial genes agree with at least one of the six chromosomes. In total, 36 of 44 mitochondrial protein and rRNA coding genes (or their fragments), were detected by BLAST in the assembled genome, thus providing additional evidence of completeness.

The incomplete assembly of the *T. testudinum* mitochondrial genome (not shown) comprises two circular chromosomes, one of 192,371 bp and the other of 136,585 bp, with no shared DNA between them. At least one additional scaffold of 90,412 bp is likely to have originated from one of the circles although the exact configuration is unclear; the scaffold has an open loop shape and overlaps with the 192,371 bp circle. The two circular chromosomes were further used in the analysis. In total 30 of 44 mitochondrial protein and rRNA coding genes (or their fragments) are encoded in the two chromosomes. However, a few additional genes detected in other small scaffolds and in nuclear chromosomes strongly suggests that the assembly status of *T. testudinum* mitochondrial genome remains incomplete.

Only a single mitochondrial contig (132,748 bp) was found for *P. oceanica* and was used for further analysis (not shown). This contig contains 20 of 44 mitochondrial protein and rRNA coding genes (or their fragments) confirming the fragmented status of the assembly. Additional mitochondrial genes were detected on nuclear chromosomes.

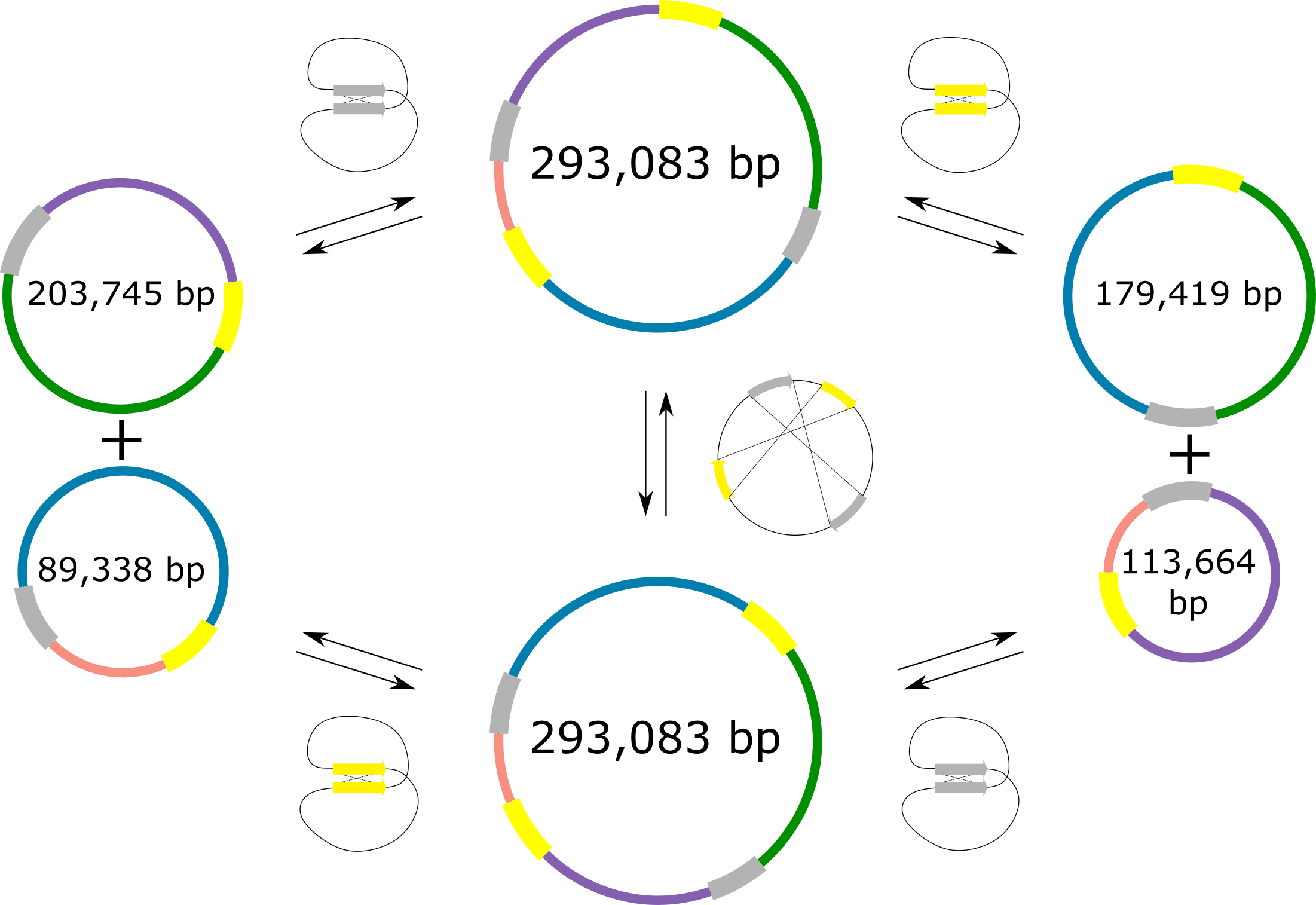

##### **Supplementary Fig. S2.3** *C. nodosa* mitochondrial chromosomes according to genome recombination activity.

See text for explanation and discussion. The two yellow and two grey segments represent two pairs of 2,683 bp and 2,476 bp respectively. Other colored segments represent parts of the mitogenome that remain unbreakable during recombination. The two smaller circles (a.k.a subgenomes) are formed when one repeat pair recombines, while the mirrored conformation is the result of recombination via both repeat pairs simultaneously.

#### 2.4 Nuclear-mitochondria and nuclear-chloroplast transfer

##### **Supplementary Note S2.4** Nuclear-mitochondria (NUMTs) and nuclear-chloroplast (NUPTs) integrants.

Nuclear-mitochondria (NUMTs) and nuclear-chloroplast (NUPTs) integrants were identified by BLAST using the parameters suggested in (Smith et al. 2011). Shared-overlapped regions in nuclear chromosomes were joined into “joined NUMTs” and “joined NUPTs”.

The integrated organellar DNAs within the nuclear genome have been shown to play important roles. Several processes can enhance DNA transfer including biotic and abiotic stress, increased organelle copy number or large gene-free regions composed of multiple repeats (Zhao et al. 2019; Ma et al. 2020; Zhang et al. 2020).

We assessed the intensity of intracellular DNA transfer of shared DNA segments between the nucleus and mitochondria (NUMTs), and the nucleus and chloroplasts (NUPTs). *T. testudinum* revealed large uninterrupted NUMTs that point towards relatively recent mitochondrial-DNA transfer, whereas *Z. marina* has very few (Suppl. Table S2.4). The recent LTR/Gypsy burst in *T. testudinum* (Suppl. Fig. S4.1) is probably the main cause for *T. testudinum*’ s extreme genome size and intron expansion. It is further correlated with increased intracellular DNA transfer from organellar to nuclear genome. Transposable elements have been proposed to contribute to the post-insertion dynamics of the transferred DNA rather than the insertion rate (Michalovova et al. 2013). Thus, multiple insertions of organellar DNA may be another consequence of a general genome instability caused by TE expansion.

##### **Supplementary Table S2.4** NUMTs and NUPTs.

Intracellular DNA transfer and the relative age of shared DNA segments between the nucleus and mitochondria (NUMTs), and the nucleus and chloroplasts (NUPTs). Recent insertions are longer (e.g., *T. testudinum*). Over time, segments become shorter (e.g., *Z. marina*).

| **Species** | Mitogenome status | Total length | Max joined NUMT length | Mean joined NUMT length | Joined NUMT >10000bp | Percent shared positions with nucleus |
| --- | --- | --- | --- | --- | --- | --- |
| ***Cymodocea nodosa*** | complete | 292,353 | 71,177 | 796 | 3 | 59 |
| ***Posidonia oceanica*** | fragment | 132,748 | 49,857 | 880 | 12 | 99 |
| ***Thalassia testudinum*** | incomplete | 328,953 | 95,806 | 1,275 | 108 | 100 |
| ***Zostera marina v3.1*** | complete | 187,048 | 9,623 | 391 | 0 | 26 |
| **Species** | Chloroplast genome status | Total length | Max joined NUPT length | Mean joined NUPT length | Joined NUPT >3000bp | Percent shared positions with nucleus |
| ***Cymodocea nodosa*** | complete | 158,459 | 11,621 | 387 | 18 | 94 |
| ***Posidonia oceanica*** | complete | 152,209 | 30,959 | 289 | 29 | 99 |
| ***Thalassia testudinum*** | complete | 180,136 | 129,692 | 527 | 181 | 100 |
| ***Zostera marina v3.1*** | complete | 143,968 | 6,039 | 396 | 1 | 44 |

### 3. Genome annotation

#### 3.1 Non-protein coding RNA annotations

##### **Supplementary Note S3.1** rRNA, tRNA and snoRNAs

The prediction of non-protein coding RNA families (i.e., rRNAs, tRNA and snoRNAs) in *Z. marina*, *C. nodosa*, *P. oceanica*, *T. testudinum* and *P. acutifolius* highlighted an overall expansion in the number of loci in *T. testudinum* when compared to other species (Suppl. Table S3.1). The 5S rRNA (119 nucleotides in length), the minor component of the large subunit of the ribosome, was detected in 10,923 distinct loci in the *T. testudinum* genome, and in 3,566 loci in *P. acutifolius*, 2,605 in *C. nodosa*, 730 in *P. oceanica*, and 364 in *Z. marina*. Clusters of the repetitive units of the large and small ribosomal subunits, organized by the 18S followed by the 5.8S and the 25S rRNAs, of 154, 3,401 and 1,851 nucleotides in length, respectively, were found on a single chromosome in *C. nodosa* (chr 18) and *P. oceanica* (chr 6), on three different chromosomes in *T. testudinum* (chr4, chr5 and chr8) and in different scaffolds in *Z. marina* and *P. acutifolius* (Suppl. Table 3.1 and Suppl. Fig. S3.1). All the repetitive units show an almost conserved organization and the length of the two internal transcribed sequences (ITSs) within a species is conserved too, even in the repeated loci along the three different chromosomes in *T. testudinum*.

Of note, *C. nodosa* does not show duplications of the represented RNA families, while an evident expansion, also accompanied by the snoRNAs class, is revealed for *T. testudinum* (Suppl. Table S3.1). This expansion appears also evident in *P. acutifolius* scaffolds, although the scaffold nature of this genome did not allow to confirm the real trend. Also transfer RNA (tRNA) sequences, with a length of 71 nt, were more abundant in *T. testudinum* (1,200 hits) and *P. acutifolius* (4,954), compared to *C. nodosa* (228), *P. oceanica* (367), and *P. oceanica* (478) species. Interestingly, snoRNAs, the RNA class that guides chemical modifications of other rRNAs, show a strong overrepresentation in *T. testudinum* (12,846 hits), compared to the other species (224 in *P. acutifolius*, 99 in *C. nodosa*, 238 in *P. oceanica*, 155 in *Z. marina*), although the number of loci detected in the *P. acutifolius* are lower than *P. oceanica* (Suppl. Table S3.1). This will require deeper investigations when the *P. acutifolius* chromosomes will be defined.

##### **Supplementary Table S3.1** Number of loci of major families of non-protein coding RNAs detected in seagrasses.

| Species | Genomic element | 5S | 5.8S | LSU_EUK(28S) | SSU_EUK(18S) | tRNA | snoRNA |
| --- | --- | --- | --- | --- | --- | --- | --- |
| *Posidonia oceanica* | Chr1 | 3 | - | - | - | 68 | 57 |
|  | Chr2 | 3 | - | - | - | 51 | 33 |
|  | Chr3 | 2 | - | - | - | 27 | 24 |
|  | Chr4 | - | - | - | - | 32 | 16 |
|  | Chr5 | 18 | - | - | 1 | 53 | 21 |
|  | Chr6 | - | 160 | 155 | 153 | 43 | 19 |
|  | Chr7 | 2 | 1 | 1 | - | 35 | 7 |
|  | Chr8 | 1 | - | - | - | 12 | 15 |
|  | Chr9 | 2 | - | - | - | 19 | 19 |
|  | Chr10 | 699 | 1 | 1 | - | 20 | 25 |
|  | Scaffolds | - | - | - | - | 7 | 2 |
|  | TOTAL | 730 | 162 | 157 | 154 | 367 | 238 |
| *Cymodocea nodosa* | Chr1 | 2,195 | - | - | 1 | 8 | 10 |
|  | Chr2 | 17 | - | - | - | 20 | 6 |
|  | Chr3 | 21 | - | - | - | 19 | 10 |
|  | Chr4 | 20 | - | - | - | 23 | 4 |
|  | Chr5 | 15 | - | - | - | 22 | 11 |
|  | Chr6 | 18 | - | - | - | 8 | 9 |
|  | Chr7 | 17 | - | - | - | 14 | 6 |
|  | Chr8 | 19 | 1 | - | - | 21 | 3 |
|  | Chr9 | 17 | - | - | - | 6 | 3 |
|  | Chr10 | 13 | - | - | - | 13 | 1 |
|  | Chr11 | 16 | - | - | - | 16 | 4 |
|  | Chr12 | 17 | - | - | - | 1 | 4 |
|  | Chr13 | 146 | - | - | - | 3 | 6 |
|  | Chr14 | 23 | - | - | - | 13 | 3 |
|  | Chr15 | 16 | - | - | - | 7 | 3 |
|  | Chr16 | 15 | - | - | - | 6 | 7 |
|  | Chr17 | 13 | - | - | - | 15 | 6 |
|  | Chr18 | 7 | 51 | 49 | 49 | 13 | 3 |
|  | TOTAL | 2,605 | 52 | 49 | 50 | 228 | 99 |
| *Zostera marina* | Chr1 | 6 | - | - | 1 | 55 | 14 |
|  | Chr2 | 5 | - | - | 1 | 79 | 7 |
|  | Chr3 | 2 | - | - | - | 59 | 33 |
|  | Chr4 | 5 | 1 | 1 | - | 48 | 39 |
|  | Chr5 | 4 | 1 | - | - | 47 | 30 |
|  | Chr6 | 1 | - | - | - | 55 | 20 |
|  | Scaffolds | 341 | 120 | 90 | 124 | 135 | 12 |
|  | TOTAL | 364 | 122 | 91 | 126 | 478 | 155 |
| *Thalassia testudinum* | Chr1 | 27 | - | - | 1 | 120 | 1,619 |
|  | Chr2 | 41 | - | - | - | 137 | 1,712 |
|  | Chr3 | 28 | - | - | - | 94 | 1,729 |
|  | Chr4 | 33 | 74 | 73 | 73 | 126 | 1,553 |
|  | Chr5 | 16 | 67 | 61 | 64 | 125 | 1,387 |
|  | Chr6 | 3,435 | - | - | 3 | 113 | 1,459 |
|  | Chr7 | 41 | 8 | 5 | 6 | 84 | 1,306 |
|  | Chr8 | 7,294 | 111 | 106 | 107 | 75 | 1,189 |
|  | Chr9 | 8 | - | - | - | 40 | 885 |
|  | Scaffolds | - | - | - | - | 286 | 7 |
|  | TOTAL | 10,923 | 260 | 245 | 254 | 1,200 | 12,846 |
| *Potamogeton acutifoliis* | Scaffolds | 3,566 | 1,548 | 1,665 | 1,693 | 4,954 | 224 |

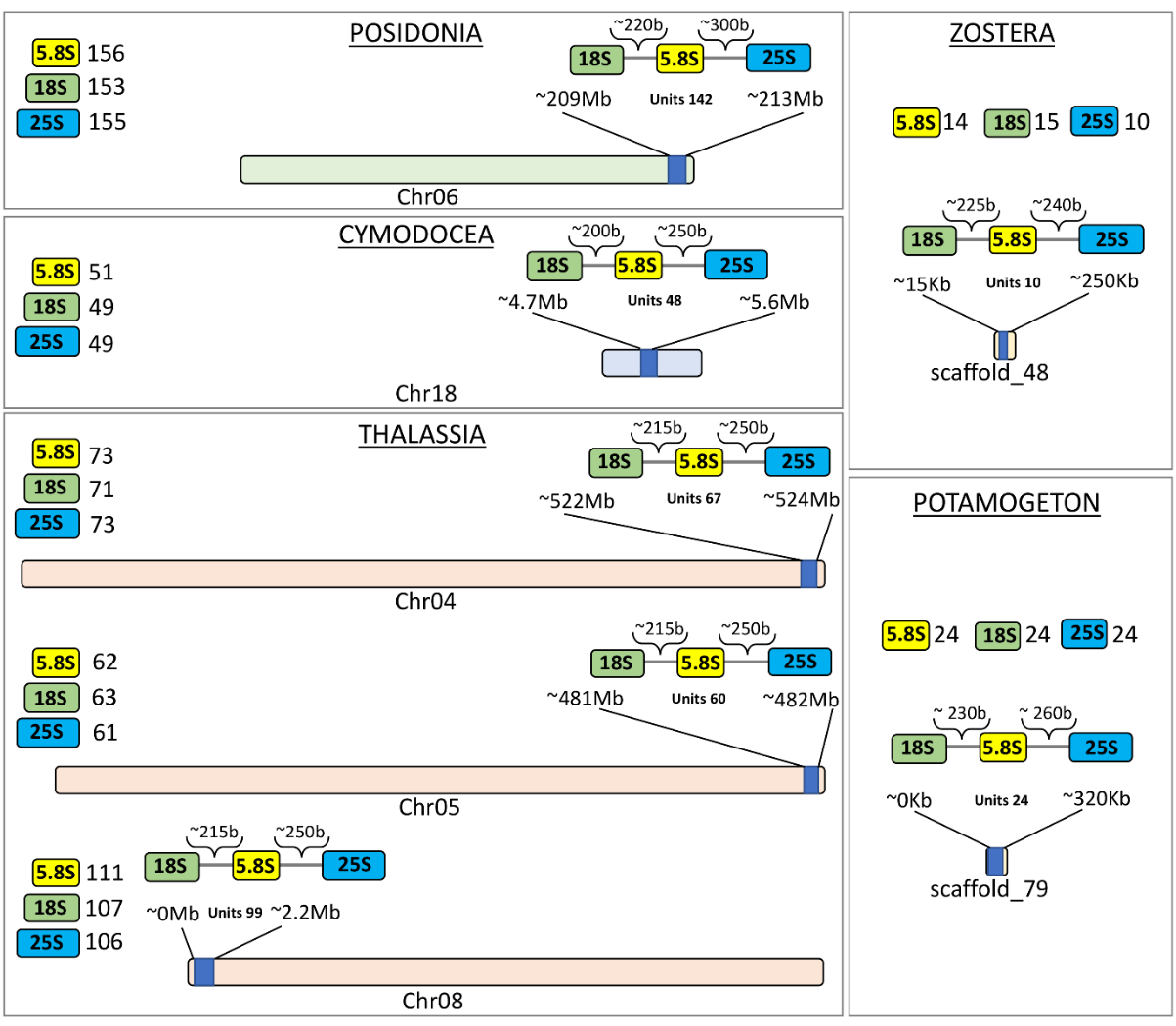

##### **Supplementary Fig. S3.1** Organization of 18S, 5.8S and 28S rRNA repeat units and in the clusters on seagrasses chromosomes.

Considering *Zostera marina* and *Potamogeton acutifolius*, the organization of the longest clusters and the scaffold in which they have been found. The length of the ITSs and the number of repeats in a cluster (units number) are reported.

#### 3.2 Transcriptome libraries, sequencing, and assembly

See Methods and Suppl. Table S1.1 for RNA preparation

##### **Supplementary Note S3.2** Transcriptome libraries

Strand-specific RNASeq library(s) were created and quantified by qPCR. RNA sequencing was performed using an Illumina instrument (Suppl. Tables S3.2.1 – Suppl. Table S3.2.4). Raw fastq file reads were filtered and trimmed using the JGI QC pipeline resulting in the filtered fastq file. Using BBDuk (https://sourceforge.net/projects/bbmap/), raw reads were evaluated for artifact sequence by kmer matching (kmer=25), allowing 1 mismatch and detected artifact was trimmed from the 3' end of the reads. RNA spike-in reads, PhiX reads and reads containing any Ns were removed. Quality trimming was performed using the phred trimming method set at Q6. Finally, following trimming, reads under the length threshold were removed (minimum length 25 bases or 1/3 of the original read length - whichever is longer).

Filtered fastq files were used as input for de novo assembly of RNA contigs. Reads were assembled into consensus sequences using Trinity (v2.11.0) (Grabherr et al. 2011). Trinity partitions the sequence data into many individuals de Bruijn graphs, each representing the transcriptional complexity at a given gene or locus, and then processes each graph independently to extract full-length splicing isoforms and to tease apart transcripts derived from paralogous genes. Trinity combines three independent software modules: Inchworm, Chrysalis, and Butterfly, applied sequentially to process large volumes of RNA-seq reads. Trinity was run with the --normalize_reads (In-silico normalization routine) and --jaccard_clip (Minimizing fusion transcripts derived from gene dense genomes) options.

##### **Supplementary Table S3.2.1** Transcriptome sequencing data for *Thalassia testudinum*

|  | **Sequencing** | **Tissue** | **library ID** | **Replicate** | **Number of raw Reads** | **Number of clean Reads** | **Number of mapped reads** | **% of aligned reads** |
| --- | --- | --- | --- | --- | --- | --- | --- | --- |
| Summary of transcriptome sequencing in *Thalassia* | RNA-seq  (NovaSeq S4) | Rhizome | GZGXW | 1 | 121,598,766 | 118,237,066 | 114,398,701 | 96.75 |
|  |  |  | GZGXX | 2 | 73,636,900 | 70,598,156 | 67,892,538 | 96.17 |
|  |  |  | GZGXY | 3 | 132,873,546 | 129,273,100 | 124,854,697 | 96.58 |
|  |  | Leaf | GZGXT | 1 | 93,743,754 | 90,902,870 | 87,180,663 | 95.91 |
|  |  |  | GZGXU | 2 | 62,515,064 | 61,189,816 | 58,999,410 | 96.42 |
|  | **Sequencing** | **Tissue** | **library ID** | **Replicate** | **Number of full-length ccs reads** | **Number of high-quality isoforms** | **Number of mapping reads** | **% of aligned reads** |
|  | Iso-seq  (SEQUELII) | Leaf | GYNYG | 1 | 3,645,896 | 217,994 | 347,411 | 98.4 |
|  |  |  | GYNYH | 2 | 3,267,856 | 135,187 |  |  |
|  |  | Rhizome | GYNYO | 1 | 3,002,957 | 177,124 | 351,956 | 99.1 |
|  |  |  | GYNYN | 2 | 2,801,968 | 178,044 |  |  |

##### **Supplementary Table S3.2.2** Transcriptome sequencing data for *Posidonia oceanica*

|  | **Sequencing** | **Tissue** | **library ID** | **Replicate** | **Number of raw Reads** | **Number of clean Reads** | **Number of mapping reads** | **% of aligned reads** |
| --- | --- | --- | --- | --- | --- | --- | --- | --- |
| Summary of transcriptome sequencing in *Posidonia* | RNA-seq  (NovaSeq S4) | Leaf | GZGXH | 1 | 118,219,220 | 110,207,116 | 106,925,423 | 97.02 |
|  |  |  | GYOZB | 2 | 103,334,962 | 89,188,206 | 85,643,795 | 96.03 |
|  |  |  | GZGXN | 3 | 112,093,828 | 106,876,636 | 103,508,597 | 96.85 |
|  |  | Rhizome | GZGXO | 1 | 90,749,690 | 87,505,084 | 84,244,810 | 96.27 |
|  |  |  | GZGXP | 2 | 97,902,404 | 95,308,522 | 92,137,451 | 96.67 |
|  | **Sequencing** | **Tissue** | **library ID** | **Replicate** | **Number of full-length ccs reads** | **Number of high-quality isoforms** | **Number of mapping reads** | **% of aligned reads** |
|  | Iso-seq  (SEQUELII) | Leaf | GZHTC | 1 | 1,852,186 | 93,731 | 265,793 | 99.7 |
|  |  |  | GZHTB | 2 | 2,542,984 | 172,871 |  |  |
|  |  | Rhizome | GZHTH | 1 | 2,525,643 | 183,914 | 335,745 | 99.6 |
|  |  |  | GZHTG | 2 | 1,941,033 | 152,859 |  |  |

##### **Supplementary Table S3.2.3** Transcriptome sequencing data for *Cymodocea nodosa*

|  | **Sequencing** | **Tissue** | **library ID** | **Replicate** | **Number of raw Reads** | **Number of clean Reads** | **Number of mapping reads** | **% of aligned reads** |
| --- | --- | --- | --- | --- | --- | --- | --- | --- |
| Summary of transcriptome sequencing in *Cymodocea* | RNA-seq  (NovaSeq S4) | Leaf | GZGWX | 1 | 229,014,654 | 221,616,172 | 215,735,219 | 97.35 |
|  |  |  | GZGWY | 2 | 85,687,350 | 81,851,552 | 78,999,864 | 96.52 |
|  |  | Rhizome | GZGWZ | 1 | 152,179,412 | 142,836,610 | 138,185,544 | 96.74 |
|  |  | Root | GZGXA | 1 | 77,449,636 | 75,252,242 | 72,717,889 | 96.63 |
|  |  |  | GZGXB | 2 | 103,516,046 | 102,007,494 | 99,073,508 | 97.12 |
|  |  | Flower | GZGXC | 1 | 167,193,082 | 157,226,850 | 153,290,632 | 97.50 |
|  |  |  | GZGXG | 2 | 170,015,134 | 160,442,432 | 156,397,982 | 97.48 |
|  | **Sequencing** | **Tissue** | **library ID** | **Replicate** | **Number of full-length ccs reads** | **Number of high-quality isoforms** | **Number of mapping reads** | **% of aligned reads** |
|  | Iso-seq  (SEQUELII) | Mixed pool of Rhizome, root and flower | GWWWG | 1 | 1,103,846 | 130,315 | 568,012 | 98.5 |
|  |  |  | GWWWC | 2 | 2,318,964 | 171,811 |  |  |
|  |  | Leaf | GZHTN | 1 | 2,102,451 | 157,082 |  |  |
|  |  |  | GZHTO | 2 | 1,982,756 | 117,547 |  |  |

##### **Supplementary Table S3.2.4** Transcriptome sequencing data for *Potamogeton acutifolius.*

|  | **Sequencing** | **Tissue** | **library ID** | **Replicate** | **Number of raw reads** | **Number of clean reads** | **Number of mapping reads** | **% of aligned reads** |
| --- | --- | --- | --- | --- | --- | --- | --- | --- |
| Summary of transcriptome sequencing in *Potamogeton* | RNA-seq (NovaSeq 6000 S4) | Leaf | PALF | 1 | 85,063,302 | 85,063,302 | 35,512,097 | 41.7 |
|  |  | Root | PARO | 1 | 68,366,930 | 68,366,930 | 35,807,278 | 52.4 |
|  |  | Turion | PATU | 1 | 64,107,268 | 64,107,268 | 56,978,015 | 88.9 |

### 4. Genome Evolution

#### 4.1 Transposable elements

##### **Supplementary Table S4**.**1** Statistics on transposable elements (TE).

| **TE Statistics** | ***T. testudinum*** | ***P. oceanica*** | ***C. nodosa*** | ***Z. marina*** | ***P. acutifolius*** |
| --- | --- | --- | --- | --- | --- |
| Overall TE content (%) | 87.36 | 85.18 | 64.65 | 65.57 | 40.76 |
| Overall LTR (%) | 72.27 | 65.89 | 45.72 | 41.72 | 12.41 |
| LTR/Gypsy (%) | 63.18 | 57.8 | 17.43 | 32.11 | 3.39 |
| LTR/Copia (%) | 8.28 | 4.09 | 28.29 | 9.49 | 6.11 |
| LINE (%) | 5.06 | 2.63 | 2.38 | 4.05 | 0.59 |
| SINE (%) | 0.03 | 0.23 | 0.00 | 0.09 | 0.23 |
| DNA transposon (%) | 7.78 | 8.39 | 2.93 | 5.71 | 7.14 |
| Unclassified (%) | 2.21 | 2.76 | 13.61 | 14.00 | 18.95 |

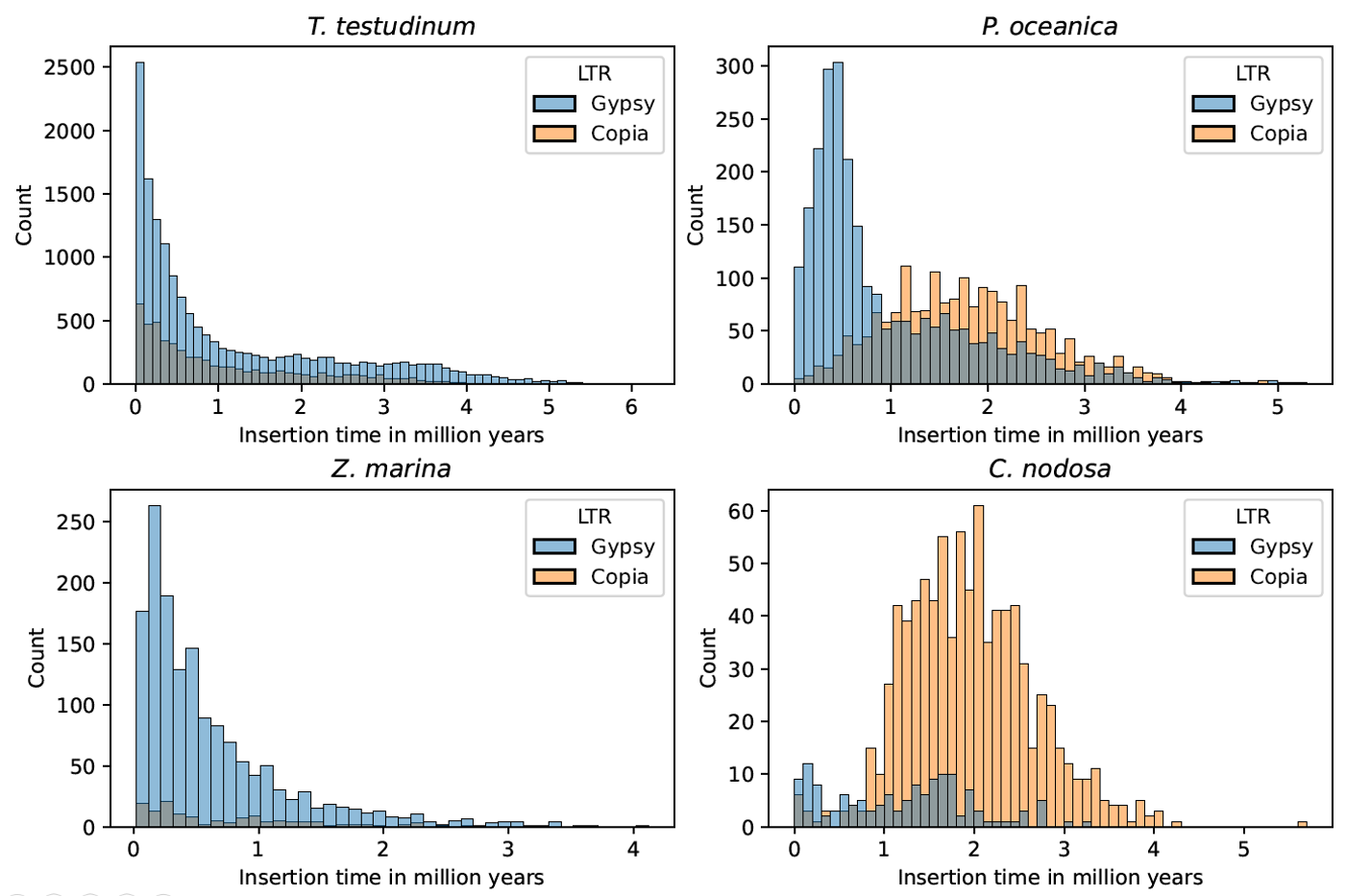

##### **Supplementary Fig. S4.1** Insertion time distributions of LTR/Gypsy and LTR/Copia in *T. testudinum*, *P. oceanica*, *C. nodosa* and *Z. marina*. Note differences in the scale of the y-axis. See main text for details.

#### 4.2 Identifying Whole Genome Duplications

##### **Supplementary Note S4.2.1** *K_S_* age distributions

*K*_S_ age distribution analysis was performed using the wgd package (Zwaenepoel and Van de Peer 2019b). The paranome (the entire collection of duplicated genes) was obtained with ‘wgd mcl’ using all-against-all BlastP and MCL clustering. Anchor pairs (i.e., paralogous genes lying in collinear or syntenic regions of the genome) were obtained using i-ADHoRe (Simillion et al. 2008), employing the default settings in ‘wgd syn’. *K*_S_ distribution analysis was also performed using the KSRATES software (Sensalari et al. 2021), which locates ancient polyploidization events with respect to speciation events within a phylogeny. It compares paralog and ortholog *K*_S_ distributions, while correcting for substitution rate differences across the involved lineages. First, an all-versus-all amino acid level similarity search for the set of protein-coding sequences was conducted using BLAST (v2.6.0+) with an E-value cut-off of 1e−10. The resulting sequence similarity graph was clustered by the Markov clustering algorithm mcl (v10-201) (Dongen 2008) with default inflation factor to identify the paralogous gene families. Second, a codon-level, multiple sequence alignment (MSA) was obtained by inferring an amino acid MSA using the MUSCLE (v3.8.31) (Edgar 2004) under default parameters, which was then back-translated to a codon-level nucleotide MSA. The gene families which had >200 members or an MSA length less <100 amino acids were filtered out. A maximum likelihood estimate of each gene pair in the family (for the pairwise synonymous distance (*K*_S_)) was obtained using the CODEML program or the PAML (Yang 2007b) package (v4.9j). Third, an estimate of the phylogenetic tree topology of each paralogous gene family was obtained using Fasttree (v2.1.7) (Price et al. 2010) and rooted using midpoint rooting.

The most recent common ancestor (MRCA) node depth, of each gene pair, was associated with the pairwise *K*_S_ estimate as a weight, in which for each duplication node in the phylogenetic tree, all n pairwise *K*_S_ estimates of the descendant clades were added to the *K*_S_ distribution with a weight of 1/n to reduce redundancy. The *K*_S_ ortholog age distributions were based on the one-to-one orthologs (reciprocal best hits) between two species using BLAST (v2.6.0+) with an E-value cut-off of 1e−10 following the same *K*_S_ estimation process as for the paranome age distributions. The substitution rate correction across different species was achieved as follows: 1) The original divergence times between the focal species and the other species (represented as *K*_S_ distance) were obtained by the mode of the ortholog *K*_S_ distributions using bootstrapped kernel density estimation; 2) A substitution-rate-adjusted *K*_S_ estimate was calculated by transforming the original *K*_S_ distance into branch-specific *K*_S_ distance using the reference of outgroup species and then rescaling the *K*_S_ distance with the diverged species into the *K*_S_ timescale of the focal species for each divergence event. The corrected divergence times in *K*_S_ timescale were then comparable with the paranome *K*_S_ distribution of focal species after above processes. The maximum number of outgroup species/trios selected to correct each divergent species pair was set as 6 and the consensus peak for multiple outgroups was set as best outgroup in KSRATES. Other parameters were set as default for rate correction using KSRATES. The species tree adopted consists of *Thalassia testudinum*, *Cymodocea nodosa*, *Posidonia oceanica*, *Potamogeton acutifolius*, *Zostera marina*, *Spirodela polyrhiza*, *Wolffia Australiana*, with *Brachypodium distachyon* and *Oryza sativa* as outgroup species, covering all the four families of seagrasses (Posidoniaceae, Zosteraceae, Hydrocharitaceae and Cymodoceaceae).

The syntenic analysis was performed by MCscan (Python version) with default parameters (Tang et al. 2008). Collinear blocks containing fewer than five orthologous gene pairs were filtered out. The collinearity information of the gene set and other genomic features in the seagrass’s genomes were visualized by Tbtools and Circos(Krzywinski et al. 2009; Chen et al. 2020).

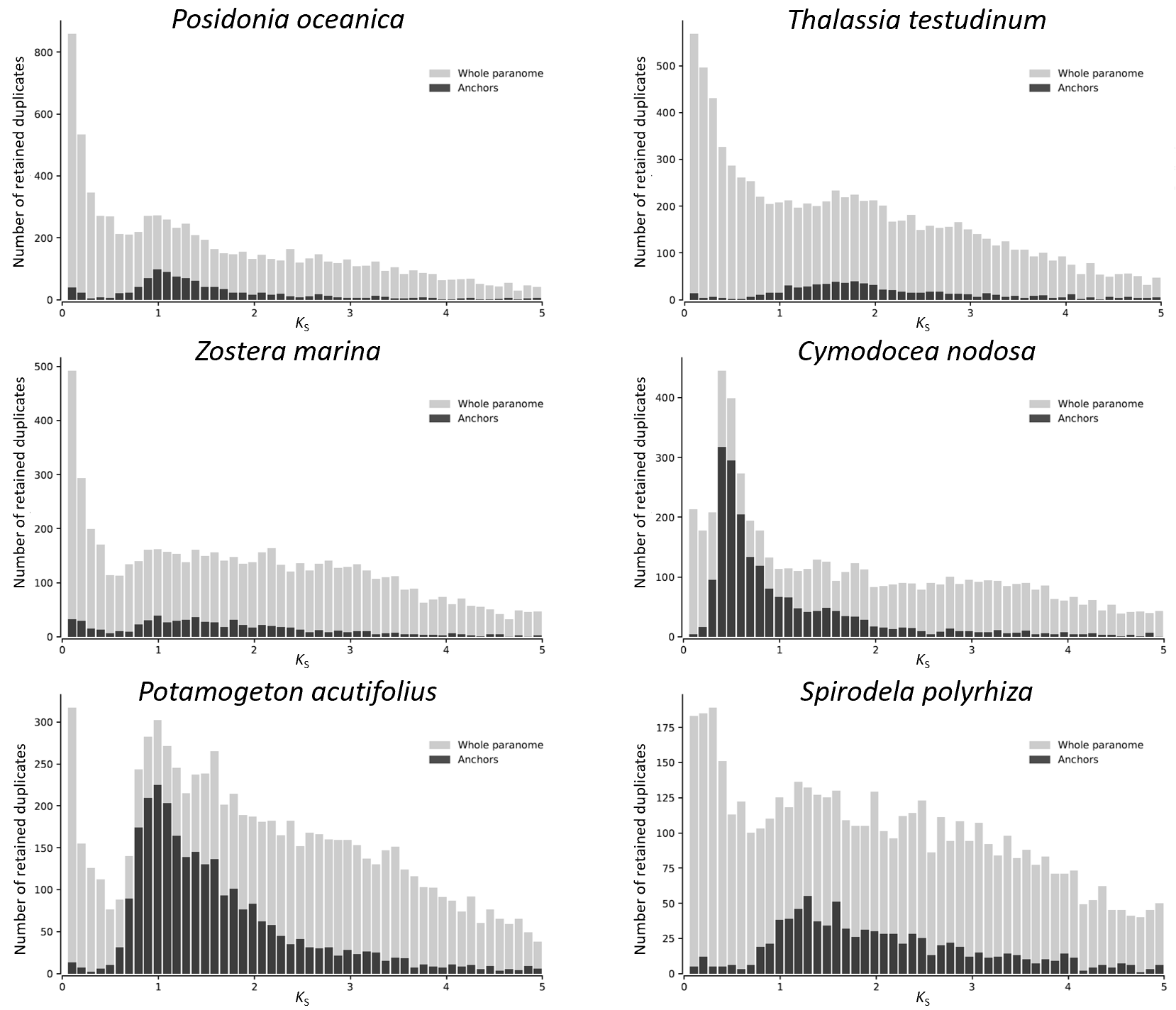

##### **Supplementary Fig. S4.2.1** *K*_S_ distributions for anchor pair duplicates (duplicates laying in duplicated, colinear blocks) and the whole paranome of four seagrasses, as well as for *P. acutifolius* and *S. polyrhiza*, generated by the wgd software (see Methods)*.*

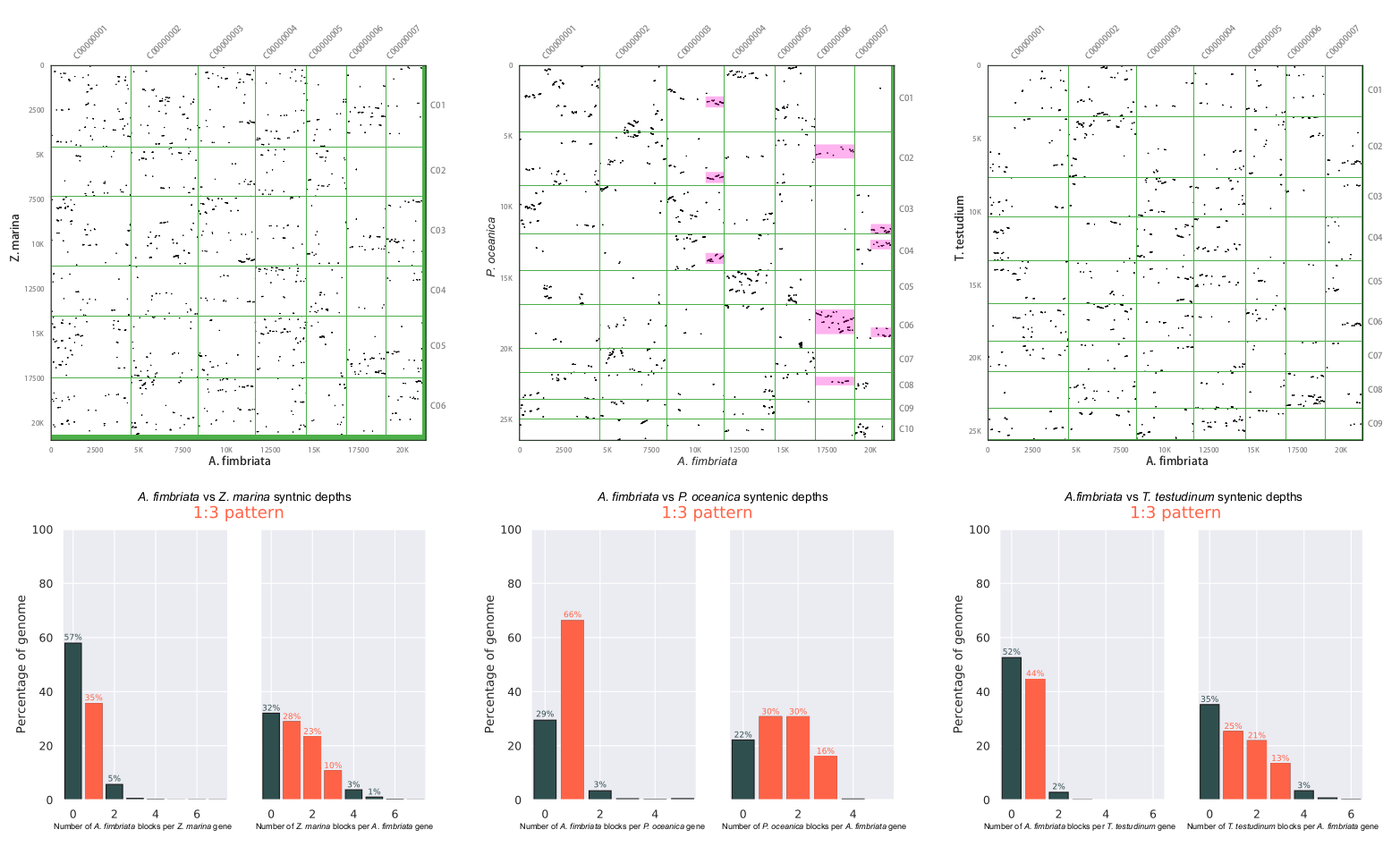

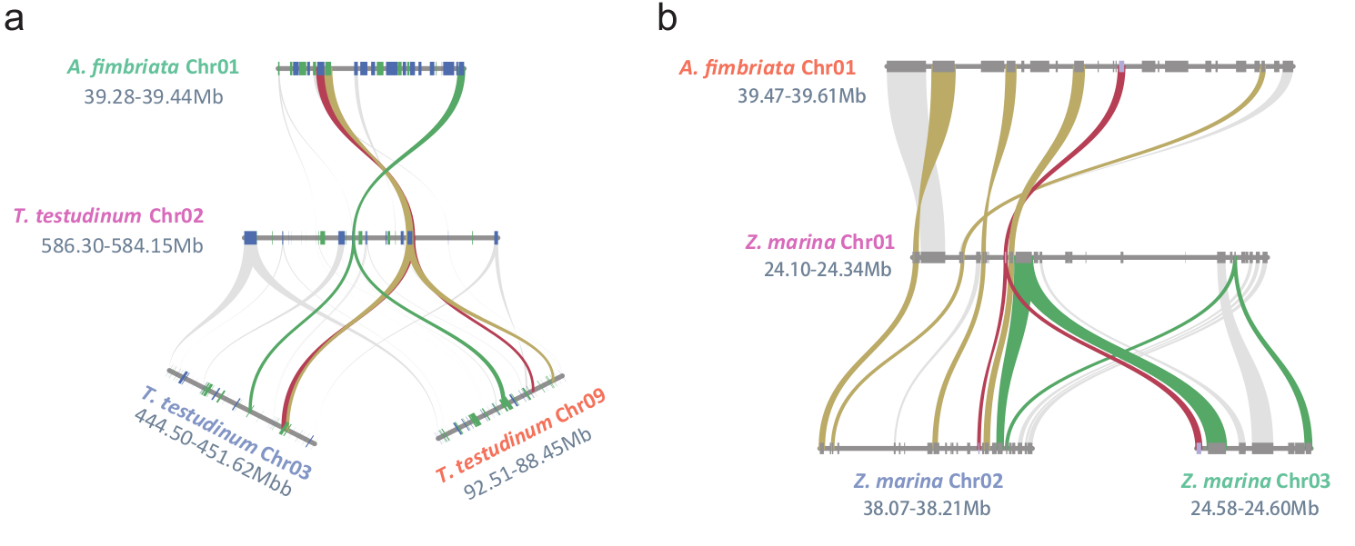

##### **Supplementary Fig. S4.2.2** Comparison of *Z. marina*, *P. oceanica*, and *T. testudinum* with *Aristolochia fimbriata*.

From the dot plots, only the comparison between *A. fimbriata* and *P. oceanica* shows a clear 1:3 relationship. However, closer inspection does also show several triplicated genomic blocks for many *A. fimbriata* genes (middle panel). The lower panel shows an example of such cases for *A. testudinum* and *Z. marina*, respectively.

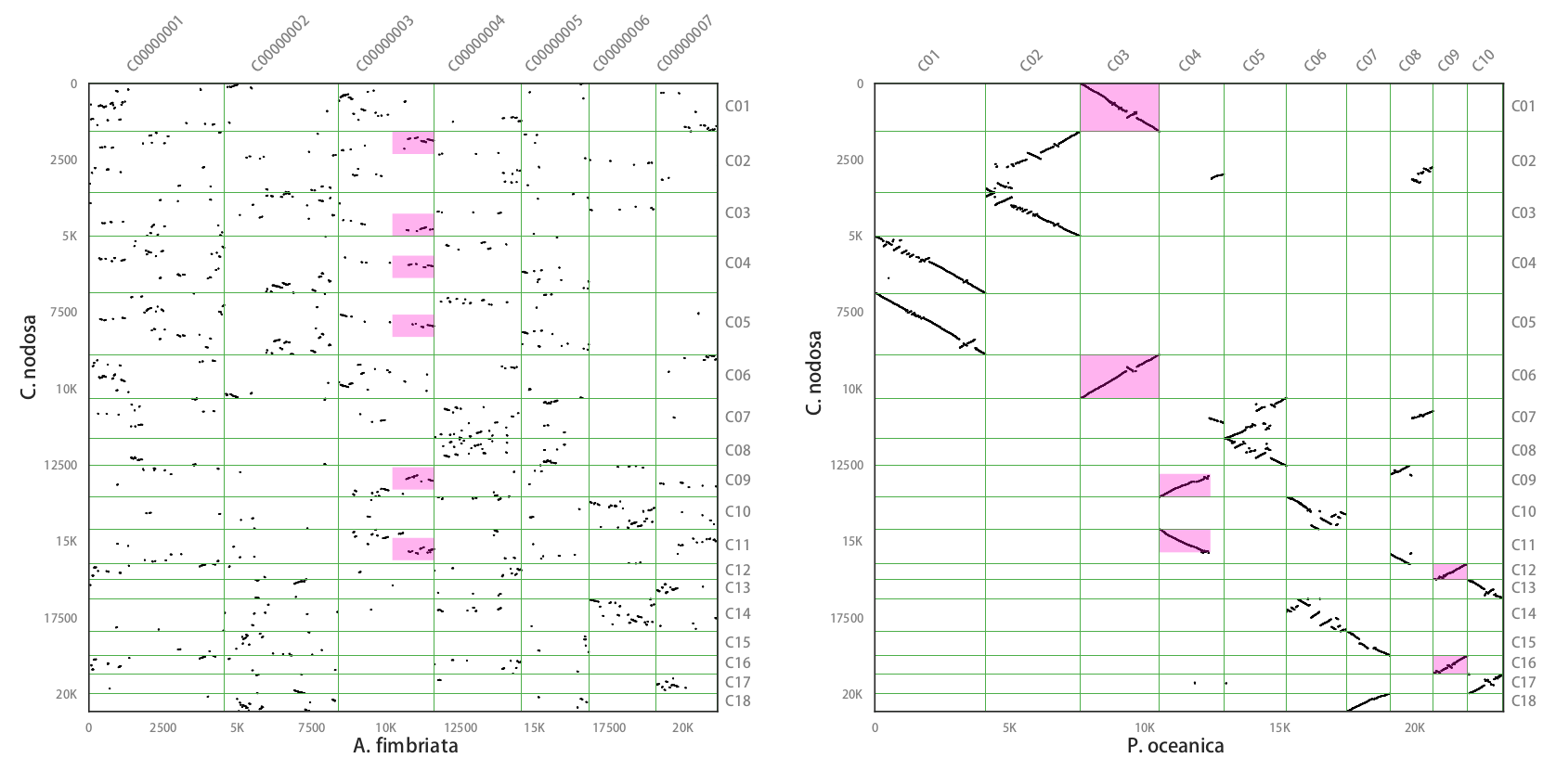

##### **Supplementary Fig. S4.2.3** Comparison of *P. oceanica* and *Aristolochia fimbriata* with *C. nodosa*.

From the dot plots, the comparison between *A. fimbriata* and *C. nodosa* shows a 1:6 synteny relationship, while comparison of *P. oceanica* and *C. nodosa* shows a 1:2 synteny relationship, in support of an extra WGD in *C. nodosa*, after its divergence with *P. oceanica*.

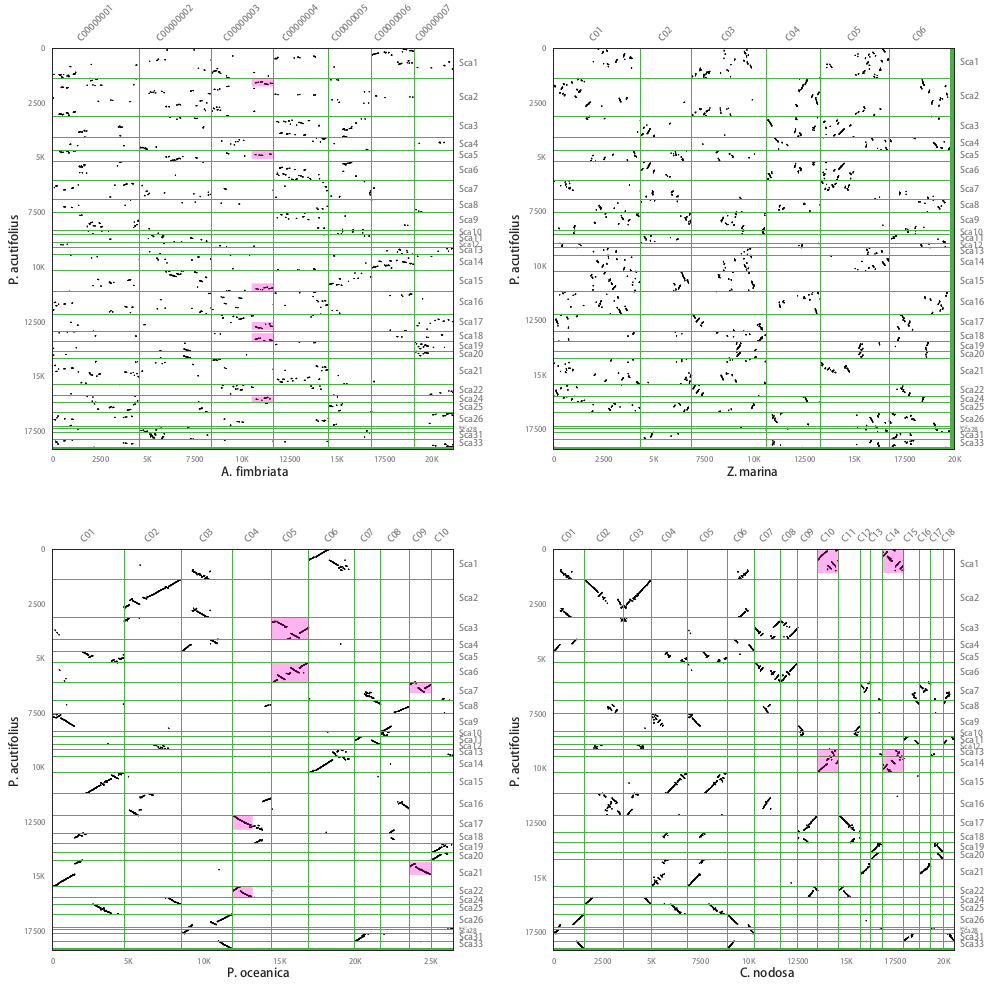

##### **Supplementary Fig. S4.2.4** Comparison of *Aristolochia fimbriata*, *Z. marina, P. oceanica* and *C. nodosa* with *P. acutifolius.*

From the dot plots, the comparison between *A. fimbriata* and *P. acutifolius* shows a 1:6 synteny relationship, while comparison of *Z. marina* and *P. acutifolius* shows a non-obvious 1:2 synteny relationship, in support of an extra WGD in *P. acutifolius*, after its divergence with *Z. marina*. *P. acutifolius* shows a 2:1 relationship with *P. oceanica*, and a 2:2 relationship with *C. nodosa*, strongly confirming their respective extra WGDs.

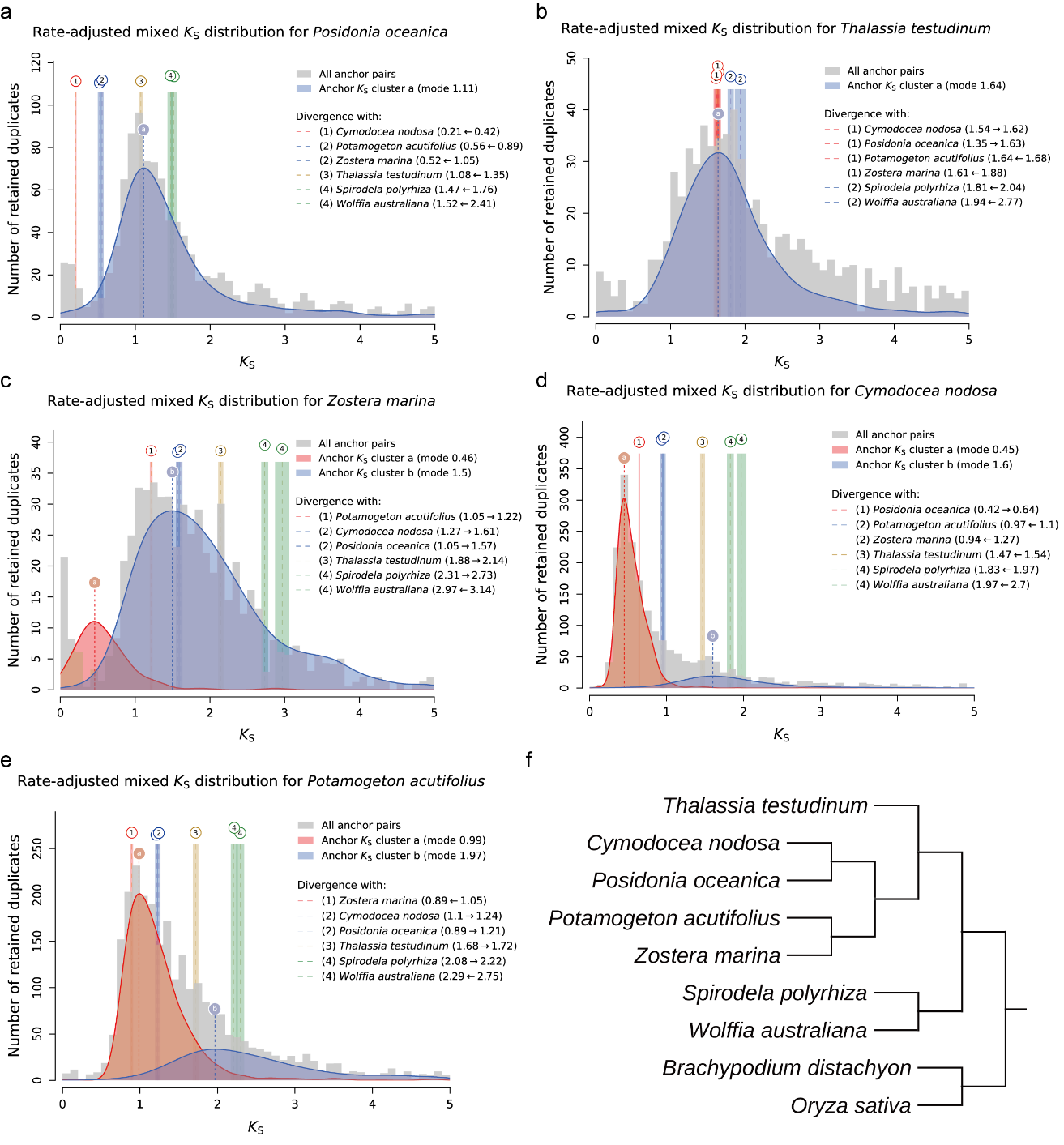

##### **Supplementary Fig. S4.2.5** *K*_S_ Distributions for paralogs and the whole paranome of four seagrasses and *P. acutifolius* generated by KSRATES software.

a – e, *K*_S_ distributions in *P. oceanica*, *T. testudinum*, *Z. marina*, *C. nodosa* and *P. acutifolius*. F, topology used in KSRATES analysis.

##### **Supplementary Note S4.2.2** Gene tree-species tree reconciliation

OrthoFinder (Emms and Kelly 2019) (v2.3.3) was used to build orthologous gene families with the inflation factor set to 3.0 and the remaining parameters set as default. Gene families that did not have at least one gene from both clades at the root or have a family size exceeding 2 times the median of the square root of the family size, based on a Poisson outlier criterion, were filtered out (Zwaenepoel and Van de Peer 2019a). An amino acid, multiply sequence alignment (MSA) was obtained using PRANK (Loytynoja and Goldman 2005) for each gene family and the resulting MSA was then used as input for the Markov Chain Monte Carlo (MCMC) analysis in mrbayes (Huelsenbeck and Ronquist 2001) (v3.2.6) to sample from the posterior probability distribution. The rate matrix for amino acid data (Aamodelpr) was set to a fixed (LG) and the rate was set as gamma-distributed approximating four rate categories. The sampling frequency was set to 10 and the number of generations was set to 110,000 to reach a total of 11,000 posterior samples. The ALEobserve (Szöllõsi et al. 2013) was then used to construct the conditional clade distribution (CCD) containing marginal clade frequencies with a ‘burn-in’ of 1000, based on the 11,000 posterior samples for each gene family. The topology of the species tree was set as shown in Fig. 3). The time-calibrated species tree was inferred by MCMCtree from the PAML package (Yang 2007a), using reference divergence times of 42-52 million years ago (MYA) for the most common ancestor of *Oryzae sativa* and *Brachypodium distachyon*, 118-129 MYA for that between *Spirodela polyrhiza* and *Zostera marina* and 130-140 for that between *Spirodela* and other terrestrial monocots (An et al. 2019).

The duplication-loss (DL)+WGD model, under critical and relaxed branch-specific rates, was implemented for the inference of the significance and corresponding retention rates of the assumed WGD events under Bayesian inference (BI) (Zwaenepoel and Van de Peer 2019a). In the critical-branch-specific DL+WGD model, the prior η, which denotes the parameter of the geometric prior distribution on the number of genes at the root, was set to follow a truncated-univariate-Beta distribution with shape parameters as (3,1) in the interval [0.01, 0.99]; the prior r, which denotes the mean of the branch rates distribution, was set to follow a flat distribution; the prior σ, which denotes the deviation of the branch rates distribution, was set to follow an exponential distribution with scale 0.1, λ (which denotes the duplication rate of each branch) was set to follow a multivariate normal distribution. For each branch, the loss rate μ was set to be equal to λ, while in the relaxed branch specific model, λ and μ were independent with the rates variation parameter τ set to follow an exponential distribution with scale 1 (Zwaenepoel and Van de Peer 2019a). To estimate duplication and loss rate λ and μ per branch incorporating both small-scale gene duplications and WGDs, another model without WGD nodes where all branch lengths were set as 1, the prior η was set to follow a Beta distribution with shape as (3,1) and the duplication and loss rate were set respectively to follow a normal distribution with mean as 0 and standard deviation as 5 were implemented. The Bayes Factor was calculated using the “bfact.jl” script within the public github repository of WHALE to measure the strength of evidence in favor of the assumed WGD models using the Savage-Dickey density ratio.

In total, 9 WGD(T) models were set on the branches leading to the MRCA of Potamogetonaceae, Zosteraceae, Posidoniaceae, Cymodoceaceae and Hydrocharitaceae (labelled as WGD1 or WGT1), the MRCA of Potamogetonaceae, Zosteraceae, Posidoniaceae and Cymodoceaceae (WGD2 or WGT2), the *C. nodosa* lineage (WGD3), the *P. acutifolius* lineage (WGD4), the *T. testudinum* lineage (WGD5 or WGT5), the MRCA of *S. polyrhiza* and *W. australiana* (WGD6), the *S. polyrhiza* lineage (WGD7), the *W. australiana* lineage (WGD8), the MRCA of *B. distachyon* and *O. sativa* (WGD9), respectively (Suppl. Fig. S4.2.6). Posterior mean of duplication (left) and loss (right) rates estimated under DL+WGD modelling colored on the time-calibrated species tree. In the left panel, green squares indicate the significantly supported WGDs under the relaxed branch-specific model while empty squares indicate the WGDs that are not significantly supported under the relaxed branch-specific model. In the right panel, light green squares indicate the significantly supported WGDs under the critical branch-specific model while empty squares indicate the WGDs that are not significantly supported under the critical branch-specific model.

##### **Supplementary Fig. S4.2.6** Bayesian inference of duplication, loss, and retention rates in WHALE analysis.

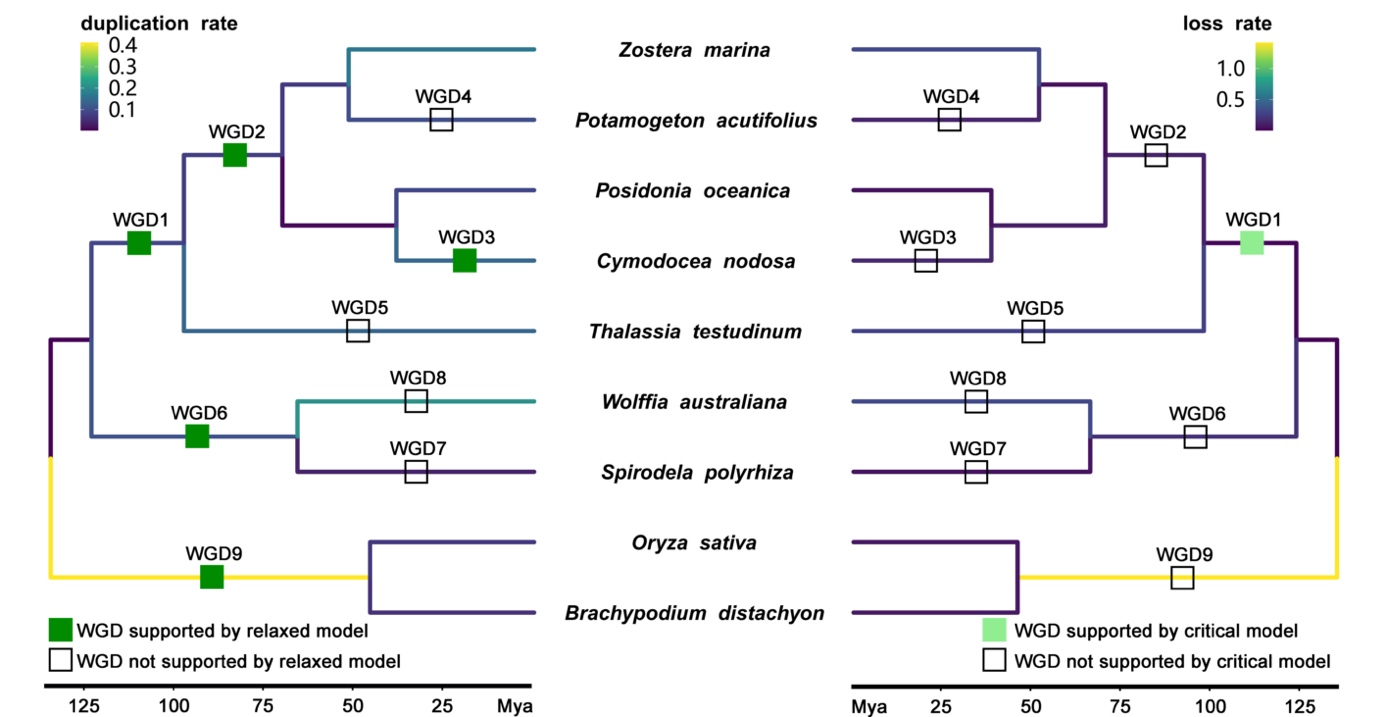

##### **Supplementary Note S4.2.3** Absolute dating of WGDs

Absolute dating of WGD events was done as described previously for *Zostera marina* (Olsen et al. 2016). Paralogous gene pairs located in duplicated segments (so called anchors) and duplicated pairs lying under the WGD peak (so-called peak-based duplicates) were collected for phylogenetic dating. Anchors, which are assumed to correspond to the most recent WGD, were detected using i-ADHoRe 3.0 (Simillion et al. 2008). For each WGD paralogous pair, an orthogroup was created that included the two paralogues plus several orthologues from other plant species, as identified by InParanoid (v4.1), using a broad taxonomic sampling, i.e., one representative from the order Cucurbitales, two from the Rosales, two from the Fabales, two from the Malpighiales, two from the Brassicales, one from the Malvales, one from the Solanales, two from the Poales, one orthologue from *Musa acuminata* (Zingiberales), and one orthologue from *Spirodela polyrhiza* (Alismatales). WGT/WGD paralogues were then dated using the BEAST v1.7 package under an uncorrelated relaxed clock model with the LG+G (four rate categories) evolutionary model. A starting tree with branch lengths satisfying all fossil-prior-constraints was created according to the consensus APGIII phylogeny. Fossil calibrations were implemented using log-normal calibration priors on the following nodes: the node uniting the Malvidae based on the fossil Dressiantha bicarpellate (Gandolfo et al. 1998) with prior offset = 82.8, mean = 3.8528, and s.d. = 0.5), the node uniting the Fabidae based on the fossil *Paleoclusia chevalieri* (Crepet and Nixon 1998) with prior offset = 82.8, mean = 3.9314, and s.d. = 0.5, the node uniting the Alismatales (including *Zostera marina* and *Spirodela polyrhiza*) with the other monocots based on the oldest fossil monocot pollen, Liliacidites (Doyle et al. 2008; Iles et al. 2015) from the Trent’s Reach locality, with prior offset = 125, mean = 2.0418, and s.d. = 0.5 (Janssen and Bremer 2004; Nauheimer et al. 2012) and the root with prior offset = 124, mean = 4.0786, and s.d. = 0.5 (Smith et al. 2010). A run without data was performed to ensure proper placement of the marginal calibration prior distributions. The Markov chain Monte Carlo (MCMC) for each orthogroup was run for 106 generations, sampling every 1,000 generations, resulting in a sample size of 104. The resulting trace files of all orthogroups were evaluated manually using Tracer v1.570 with a burn-in of 1,000 samples to ensure proper convergence (minimum ESS for all statistics at least 200).

To resolve the absolute dates of other involved WGDs in our analysis (Figure 2), we also redated the WGDs of *Elaeis guineensis*, *Asparagus officinalis*, *Rhizophora apiculata*, *Avicennia marina* and *Utricularia gibba* using the same pipeline as above. Moreover, the fossil of Sabalites carolinensis (Berry 1914) were also chosen as calibrations as previous study (Vanneste et al. 2014). The final WGD dates were shown in Suppl. Table S4.2 and Suppl. Fig. S4.2.8.

##### **Supplementary Fig. S4.2.7a** Estimation of the 'absolute age’ of the *T. testudinum* WGT event by phylogenomic dating of *T. testudinum* paralogues.

The solid black line represents the KDE (Kernel Density Estimation) of the dated paralogues, and the vertical dashed black line represents its peak at 76.12 Mya, which was used as the consensus WGD age estimate. The grey lines represent density estimates from 2,500 bootstrap replicates and the vertical black dotted lines represent the corresponding 90% confidence interval for the WGD age estimate, 70 - 82.271 Mya. The histogram shows the raw distribution of dated paralogues.

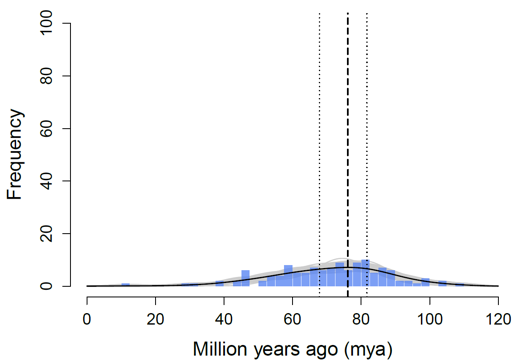

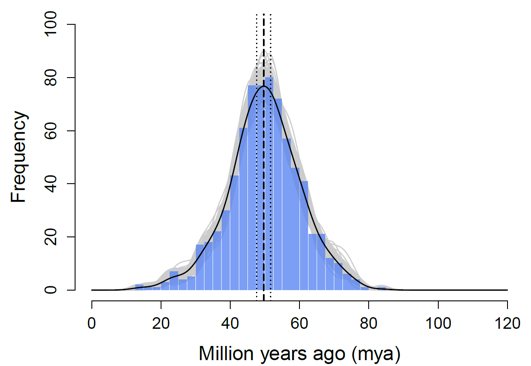

##### **Supplementary Fig. S4.2.7c** Estimation of the 'absolute age’ of the most recent *C. nodosa* WGD event, by phylogenomic dating of *C. nodosa* paralogues.

Interpretation is as in Figure S4.2.7a.

##### **Supplementary Fig. S4.2.7b** Estimation of the 'absolute age’ of the *P. oceanica* WGT event by phylogenomic dating of *P. oceanica* paralogues.

Interpretation is as in Figure S4.2.7a.

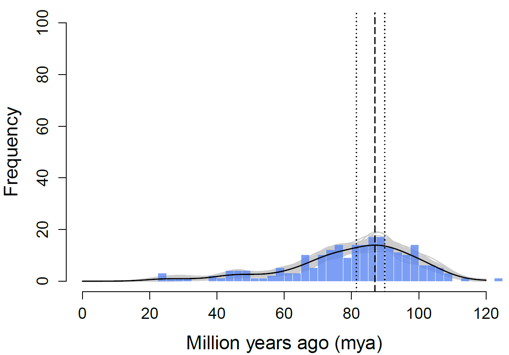

##### **Supplementary Table S4.2** The absolute of WGD events taken from literature

| **Species name** | **Date** | **Family** | **Order** | **Clade** | **Phylogenetic position of WGD** | **Data Source** |
| --- | --- | --- | --- | --- | --- | --- |
| *Spirodela polyrhiza* | 83-94, 95-100 | Araceae | Alismatales | Monocots | Araceae | (Wang et al. 2014; An et al. 2019) |
| *Oryza sativa* | 63.08-69.89,100-120,110-135 | Poaceae | Poales | Commelinids | Poaceae, Poales, non-Alismatales monocots | (Tang et al. 2010; Vanneste et al. 2014; Ming et al. 2015) |
| *Elaeis guineensis* | 48.25-55.97, 96.71-110.48 | Arecaceae | Arecales | Commelinids | Partial Arecaceae, Arecales | **Our dating, see Figure S4.2.8** |
| *Asparagus officinalis* | 59.24-72.43 | Asparagaceae | Asparagales | Monocots | Asparagus | **Our dating, see Figure S4.2.8** |
| *Arabidopsis thaliana* | 49.27-50.99 | Brassicaceae | Brassicales | Rosids | Partial Brassicaceae | (Guo et al. 2018) |
| *Populus trichocarpa* | 60-65, 125-140 | Salicaceae | Malpighiales | Rosids | Specific to Populus and Salix, Core eudicots | (Tuskan et al. 2006) |
| *Rhizophora apiculata* | 50.36-56.95 | Rhizophoraceae | Malpighiales | Rosids | Rhizophoraceae | **Our dating, see Figure S4.2.8** |
| *Vitis vinifera* | 125-140 | Vitaceae | Vitales | Rosids | Core eudicots | (International Peach Genome et al. 2013) |
| *Avicennia marina* | 60.46-78.78 | Acanthaceae | Lamiales | Asterids | *Avicennia marina* | **Our dating, see Figure S4.9** |
| *Utricularia gibba* | 43.37-53.99 | Lentibulariaceae | Lamiales | Asterids | Utricularia | **Our dating, see Figure S4.2.8** |
| *Solanum lycopersicum* | 62.64-64.84 | Solanaceae | Solanales | Asterids | Solanaceae | (Vanneste et al. 2014) |

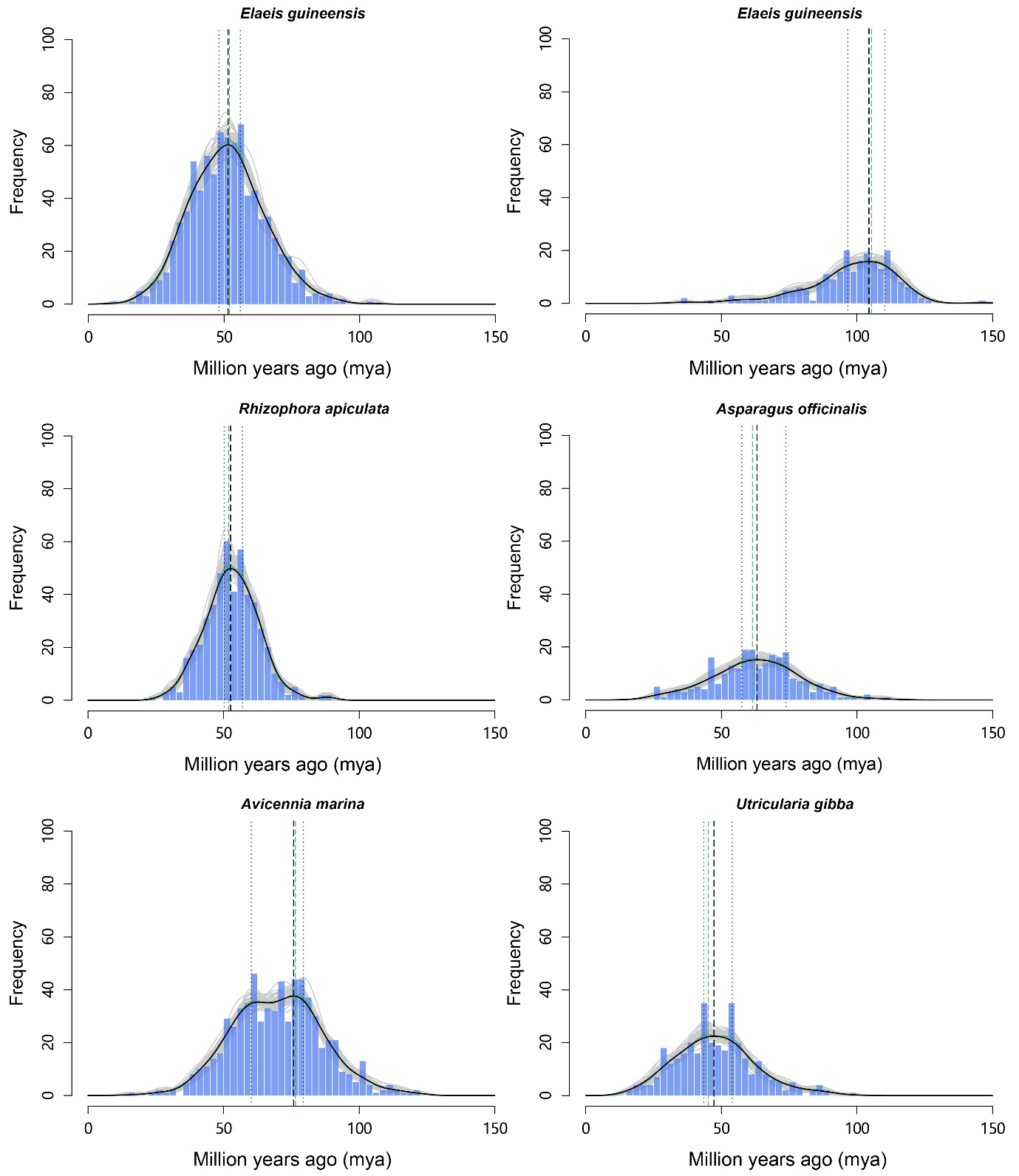

##### **Supplementary Fig. S4.2.8** Estimation of the 'absolute age’ of seven independent WGD events experienced by *E. guineensis*, *A. officinalis*, *R. apiculata*, *A. marina* and *U. gibba* respectively by phylogenomic dating of corresponding paralogues.

The solid black line represents the KDE (Kernel Density Estimation) of the dated paralogues, and the vertical dashed black line represents the peak which was used as the consensus WGD age estimate. The grey lines represent density estimates from 2,500 bootstrap replicates and the vertical black dotted lines represent the corresponding 90% confidence interval for the WGD age estimate. The histogram shows the raw distribution of dated paralogues.

#### 4.3 Phylogenetic tree construction and estimation of divergence time

##### **Supplementary Note S4.3** Species selection and construction of time-calibrated phylogeny.

Protein sets were collected for 23 species, including *Oryza sativa* (PLAZA 5.0), *Brachypodium distachyon* (PLAZA 5.0), *Ananas comosus* (PLAZA 5.0), *Elaeis guineensis* (PLAZA 5.0), *Asparagus officinalis* (PLAZA 5.0), *Beta vulgaris* (PLAZA 5.0), *Utricularia gibba* (PLAZA 5.0), *Solanum lycopersicum* (PLAZA 5.0), *Coffea canephora* (PLAZA 5.0), *Vitis vinifera* (PLAZA 5.0), *Populus trichocarpa* (PLAZA 5.0), *Arabidopsis thaliana* (PLAZA 5.0), *Theobroma cacao* (PLAZA 5.0), *Avicennia marina* (PLAZA 5.0), *Spirodela polyrhiza* (PLAZA 5.0), *Amborella trichopoda* (PLAZA 5.0), *Wolffia australiana* (https://duckweeds.plantprofile.net/), *Rhizophora apiculata* (from the author), our four seagrasses and *Potamogeton acutifolius* (from ORCAE, https://bioinformatics.psb.ugent.be/orcae/). These species were selected as representatives for monocots and eudicots, and representing different habitats from terrestrial, freshwater-floating, freshwater-submerged, to marine-submerged. Orthofinder v2.3 (Emms and Kelly 2015) was used to delineate gene families with mcl inflation factor 3.0. All-versus-all Diamond blast with an E-value cutoff of 1e−05 was performed and orthologous genes were clustered using OrthoFinder. Single-copy orthologous genes were extracted from the clustering results. MAFFT (Rozewicki et al. 2019) with default parameters was used to perform multiple sequence alignment of protein sequences for each set of single-copy orthologous genes, and to transform the protein sequence alignments into codon alignments after removing the poorly aligned or divergent regions using trimAl (Capella-Gutiérrez et al. 2009). The resulting codon alignments from all single-copy orthologs were then concatenated into one supergene for species phylogenetic analysis. A maximum-likelihood phylogenetic tree of single-copy protein alignments and codon alignments was constructed using IQ-TREE (Minh et al. 2020) with the GTR+G model and 1,000 bootstrap replicates. Divergence times between the 23 plant species were estimated using MCMCtree from the PAML package under the GTR+G with reference divergence times of 124-170 MYA for the common ancestor of monocots and eudicots, 118-129 MYA for the divergence between Spirodela and Zostera and 130-140 MYA between Spirodela and other terrestrial monocots (An et al. 2019). We used MCMCTree to obtain 10,000 trees from the posterior sampling every 150 iterations after a burn-in of 500,000 iterations. We compared two independent runs with each other to verify convergence and with a run of the MCMC algorithm under the prior alone to compare the posterior distribution for the node ages to the effective prior implied by the fossil calibrations.

### 5. Adaptation to the marine environment

#### 5.1 Pathogen resistance gene

##### **Supplementary Note S5.1** Pathogen resistance gene

Effector-triggered immunity is one of the two main arms of the plant immune system and allows angiosperms to specifically detect pathogen effectors or their impact on host proteins. The detection is guided by nucleotide-binding leucine rich repeat receptors (*NLR*s), which is one of the largest gene families in plants, and under diversifying selection (Jacob et al. 2013). Of the two main domains of *NLR* resistance genes, the nucleotide binding site (NBS) domain is responsible for downstream signaling, and the leucine rich repeat (LRR) domain binds the target.

*NLR* genes are often difficult to identify in genomes. Therefore, we used two software packages, NLR-Annotator (Steuernagel et al. 2020) and NLGenomeSweeper (Toda et al. 2020), followed by manual curation in a genome browser.

The number of *NLR* genes is strongly reduced (N=44) in *Z. marina* (Olsen et al. 2016) and similar reductions have since been found in many freshwater species (N= 100 range) (Liu et al. 2021), which is far less than in terrestrial species (N=100-300-500 [2300 in wheat]). Thus, we expected to see a similar extreme reduction in our new seagrass species but this was not the case; *C. nodosa* (N=87), *P. oceanica* (N=95) and *T. testudinum* (N=54).

Confirming our general hypothesis of convergent evolution at the genomic level, these seagrass species have low counts of *NLR* gene copies (Suppl. Table S5.1). Further, *NLR*s with a TIR domain are completely absent, which is typical of many monocots, and a few genes are missing the LRR domain. We also found that 30-40% of gene copies are non-functional in *C. nodosa*, *P. oceanica*, *T. testudinum* and *P. acutifolius*, either because of stop mutations or partial copies. By contrast, only 8% are non-functional in *Z. marina* (Suppl. Table S5.1). It is difficult to speculate why *Z. marina* has fewer degraded/partial copies (i.e., 92% are complete).

The *NLR* gene copies occur in clusters towards the terminal ends of the chromosomes consistent with findings in other plants (Jacob et al. 2013). These clusters are made up of tandem copies as evidenced by their relationship shown in the NBS-domain-based phylogenetic tree and chromosomal location (Suppl. Fig. S5.1.1 and Suppl. Fig. S5.1.2). Here, they cluster into several clades, each including all of the species, thus indicating that the ancestor also contained these gene lineages. Similarly, when incorporating other more distantly related species in the NBS tree (not shown), the seagrass-genes-branches are distributed throughout, so old *NLR* gene lineages are still maintained, despite the reduction in total number compared to other plants. Single lineages are expanded into clusters of dozens of copies at the species level, especially in *C. nodosa* and *P. acutifolius* (Suppl. Fig. S5.1.1). From an evolutionary perspective, clustering is considered as a reservoir of genetic variation (Jacob et al. 2013).

##### **Supplementary Table S5.1** *NLR* gene counts by domain architecture and completeness in seagrasses and *P. acutifolius.*

Completeness was determined by the NLR-Annotator software. Complete/partial refers to the set of motifs needed for a functional gene; pseudogene refers to loci that do not have complete open reading frames. Gene class abbreviations in parentheses are a more general designation used in some papers.

Note: Due to their partly fragmented nature not all *NLR* genes are present in the genome annotation with gene IDs. The genomic coordinates of all identified NLR genes can however be found in Extended Data Table 5.

| ***NLR* gene class** | ***Cymodocea nodosa*** | ***Posidonia oceanica*** | ***Thalassia testudinum*** | ***Zostera marina v3.1*** | ***Potamogeton acutifolius*** |
| --- | --- | --- | --- | --- | --- |
| CC-NBS (CN) | 2 | 0 | 0 | 0 | 2 |
| CC-NBS-LRR (CNL) | 59 | 58 | 27 | 21 | 71 |
| NBS-LRR (NL) | 26 | 37 | 27 | 23 | 40 |
| NBS (N) | 0 | 0 | 0 | 0 | 2 |
| TIR-NBS (TN) | 0 | 0 | 0 | 0 | 0 |
| TIR-NBS-LRR (TNL) | 0 | 0 | 0 | 0 | 0 |
| **Total** | **87** | **95** | **54** | **44** | **115** |
| **NLR gene completeness** | | | | | |
| Complete | 69 | 62 | 42 | 26 | 87 |
| Complete pseudogene | 20 | 28 | 15 | 18 | 27 |
| Partial | 13 | 14 | 12 | 0 | 25 |
| Partial pseudogene | 44 | 41 | 13 | 4 | 17 |
| **% Complete** | **61** | **62** | **70** | **92** | **73** |

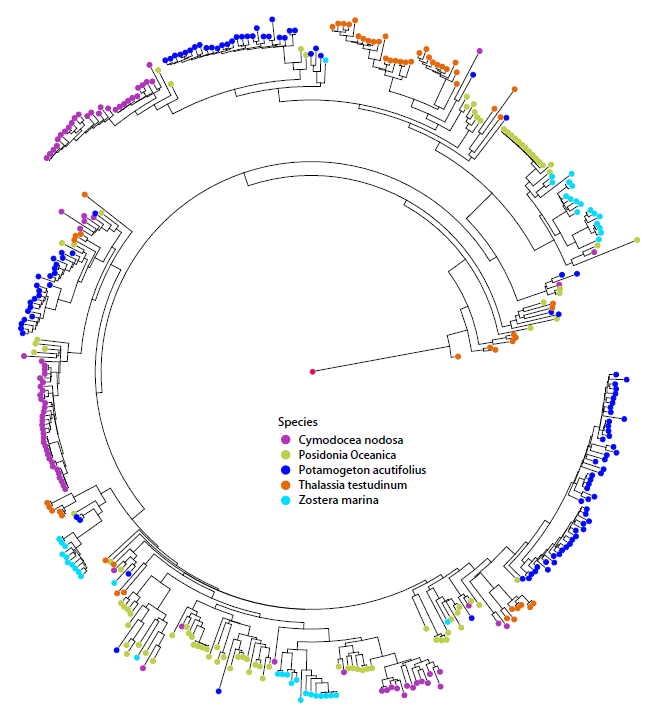

##### **Supplementary Fig. S5.1.1** Phylogenetic tree of seagrass *NLR* genes based on NBS domain.

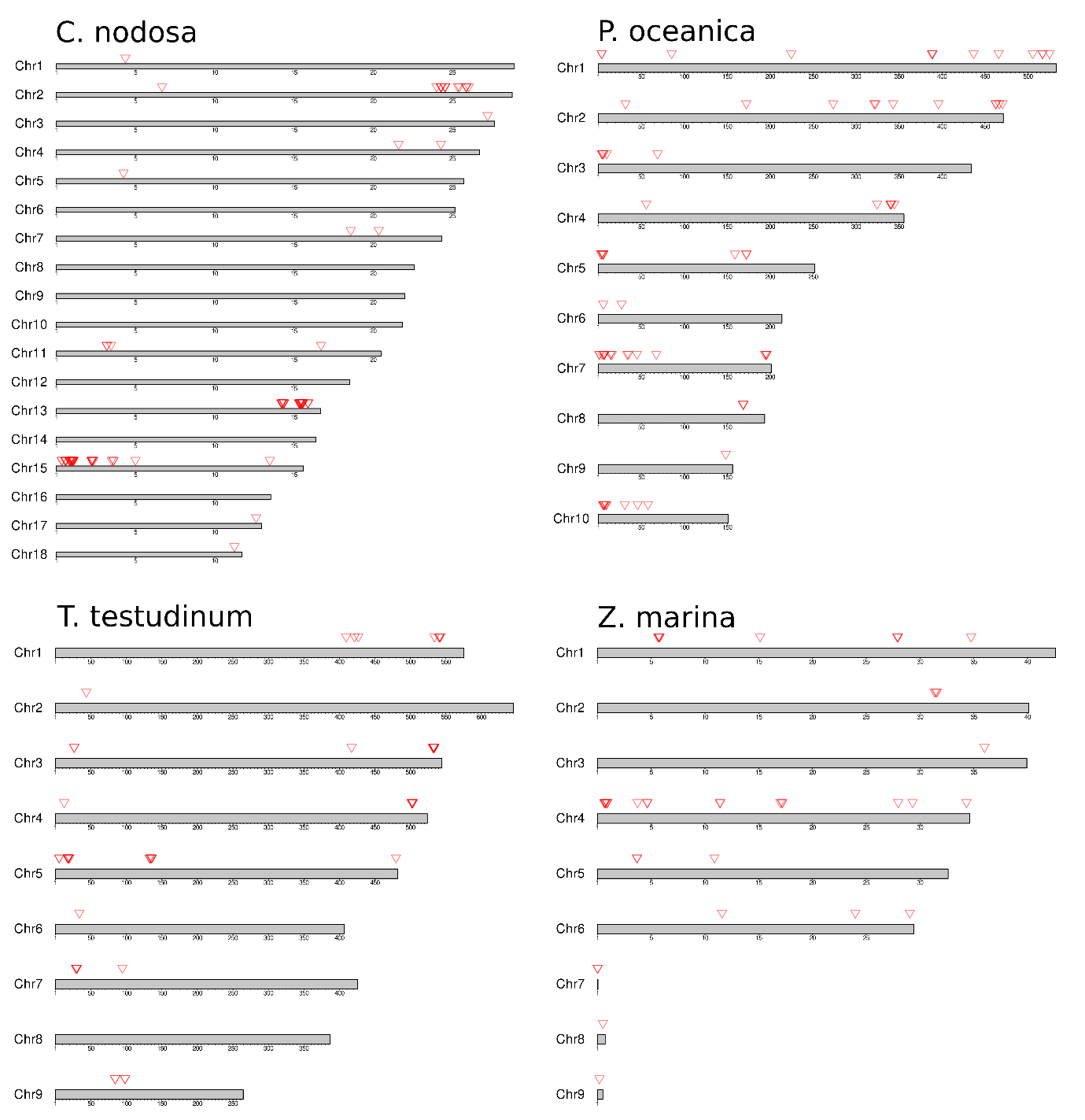

##### **Supplementary Fig. S5.1.2** Distribution of seagrass *NLR* genes across chromosomes**.**

*NLR* genes are indicated by red arrows and predominantly occur in the distal regions.

#### 5.2 Heat Shock factor (*HSF*) gene family evolution

##### **Supplementary Note S5.2** *HSF* gene family

Heat shock transcription factors (*HSF*s) are a family of DNA-binding proteins that activate a cascading network of genes that act together to enhance plant tolerance to abiotic stress conditions, including heat, cold, drought, hypoxia, salinity, toxicity and excessive irradiance (Scharf et al. 2012). Based on the topology of their domains, *HSF*s are classified into three major classes (*HSFA*, *HSFB* and *HSFC*), which are further subdivided into 16 subfamilies: *HSFA1-HSFA9*, *HSFB1-HSFB5* and *HSFC1-HSFC2*. Individual *HSF*s have unique functions as part of different signal transduction pathways operating in response to environmental stress and during plant development (von Koskull-Döring et al. 2007). To examine the different composition of *HSF* gene families in seagrasses, *HSF* families of *Cymodocea nodosa*, *Posidonia oceanica*, *Thalassia testudinum* and *Zostera marina,* were compared with those of the freshwater plant *Pomatogeton acutifolius*, ten terrestrial eudicots (*Amborella trichopoda, Arabidopsis thaliana, Beta vulgaris, Solanum lycopersicum, Populus trichocarpa, Vitis vinifera, Coffea canephora, Theobroma cacao* and the mangroves *Avicennia marina* and *Rhizophora apiculata*), five terrestrial monocots (*Oryza sativa, Brachypodium distachyon, Elaeis guineensis, Ananas comosus* and *Asparagus officinalis*) and three other freshwater plant species (one eudicot: *Utricularia gibba*; two monocots: *Spirodela polyrhiza* and *Wolffia australiana*). *HSF* sequences were searched in the Plant Tanscriptional Factor Data Base (PlantTFDB, <http://planttfdb.gao-lab.org/>) and in Phytozome 13 (<https://phytozome-next.jgi.doe.gov/>). Protein sequences were subsequently downloaded from PLAZA (<https://bioinformatics.psb.ugent.be/plaza/>), the PlantTFDB or, when needed, from the species genome web page (*Wolffia australiana*: <https://duckweeds.plantprofile.net/>; *Utricularia gibba*: <http://genomevolution.org/CoGe/>). Sequences were then uploaded to the HEATSTER platform to check their identity as *HSF*s and to use a single criterion for their classification within the 16 *HSF* sub-classes. Classification and annotation was performed in HEATSTER via two successive steps of repeated searches in a motif database (motifs: DBD and OD with HR-A and HR-B region). In those cases where a specified sequence did not contain all *HSF*-associated motifs, classification was based on the recognition of the most conserved domain (DBD) with an E-value < 1e-20.

In our analysis, the average number of *HSF*s in land plants (Suppl. Table S5.2) was similar to the values recently reported in a comprehensive study involving 29 eudicots and 10 monocots (Wang et al. 2018). Interestingly, we found that aquatic plants have a lower number of *HSF*s than terrestrial plants (54% on average), with no clear differences between marine and freshwater species. The four studied seagrass species have on average 11.8 (± 1.0) sequences recognized as *HSF*s, with all three types of *HSF*s (A, B and C) showing a strong contraction. The average number of class A and B members is reduced by 67.4% and 43% respectively in seagrasses compared to terrestrial monocots (Suppl. Table S5.2). In addition, some subclasses are completely absent in seagrasses (Extended Data Table 5). In the *HSFA* class, subclass A3, A7, A8 and A9 are lacking in seagrasses but are common in terrestrial monocots (except A9 found only in eudicots). Along with their role in the response to heat stress, these subfamilies have specialized functions in the response of plants and seeds to different abiotic stresses, mainly dehydration, drought stress and oxidative stress (Sakuma et al. 2006; von Koskull-Döring et al. 2007; Scharf et al. 2012; Personat et al. 2014). The emergence of new functionalities has been associated with the weak purifying selection of these subfamilies in terrestrial plants (Wang et al. 2018). Contrarily, subfamilies A1, A2, A5 and A6, which are subjected to more severe selection pressure and directly involved in the heat stress response (Heerklotz et al. 2001; Mishra et al. 2002; Scharf et al. 2012; Xue et al. 2014), are represented in seagrasses. These results reveal that marine plants have lost several subclasses of *HSFA* previously acquired to cope with the stress conditions associated with a terrestrial lifestyle.

Regarding the *HSFB* class, subclasses B3 and B5 are not present in seagrasses, but neither are they present in terrestrial monocots as they presumably arose after the split of monocots and eudicots (Scharf et al. 2012; Guo et al. 2016). Interestingly, seagrasses retained B1 and B2 subclasses (Extended Data Table 5), both of which are involved in promoting the activity of *HSFA1*, which is the master regulator of the heat stress response in *Arabidopsis* (Ikeda et al. 2011; Scharf et al. 2012). Moreover, *HSFB1* form a triad with *HSFA1* and *HSFA2* in tomato, acting as synergistic coactivator of heat stress responsive genes during the exposure and recovery to high temperatures (Mishra et al. 2002; Scharf et al. 2012). It is therefore evident that marine plants have conserved all major *HSF* subclasses that functionally cooperate in the heat stress response and thermotolerance of plants (A1, A2, A5, A6, B1 and B2).

*HSFC* family members (i.e., C1 and C2) are completely absent in seagrasses. Although the function of class C *HSF*s is the least known, they appear to be integrated into signaling pathways not directly related to the heat stress response (Scharf et al. 2012). Interestingly, *HSFC2* act as transcriptional activator of heat shock protein (*HSP*) genes in wheat during heat, drought and salt stress (Xue et al. 2014), and is up-regulated in rice by oxidative and heat stress (Mittal et al. 2012) . Similarly, *HSFC1* showed altered expression levels in Arabidopsis under several stress conditions, including cold stress, freeze stress and dehydration stress (Lee et al. 2005; Xin et al. 2007; Ding et al. 2013; Zhuang et al. 2018). These results indicate that the functionally specialized *HSFC* family, which emerged during the evolution of plants towards a terrestrial lifestyle, has been lost when plants returned to the sea.

In summary, after undergoing a strong expansion during the colonization of the terrestrial environment, the *HSF*s gene family has experienced a strong contraction in plants that re-colonized the marine environment (seagrasses). Despite having a small number of members, seagrasses have retained those *HSF* subfamilies under strong purifying selection in land plants, probably to maintain important biological functions. Among them is the main group of *HSF*s directly related to heat stress response and thermotolerance in plants. In contrast, subfamilies of *HSF*s with specialized functionalities for plant adaptation to terrestrial habitats and not directly related to heat stress response but to other types of abiotic stresses (e.g., drought) have been lost in seagrasses. The greater homogeneity and stability of environmental conditions at sea relative to those on land is the most likely cause of these changes. Finally, only tropical seagrasses retained some of the key heat stress-related *HSF*s from WGD and WGT events (*C. nodosa*: *HSFA1* and *HSFB4*; *T. testudinum*: *HSFB2*), which could be related to their warmer native environment and higher heat stress tolerance compared to temperate seagrasses (*P. oceanica* and *Z. marina*).

##### **Supplementary Table S5.2** Average (± SD) number of total *HSF*s and number of *HSF*s from the three main classes (*HSFA*, *HSFB* and *HSFC*) in the analyzed plant genomes.

The basal angiosperm *Amborella trichopoda* (11 sequences) and the freshwater eudicot plant *Utricularia gibba* (21 sequences) were not included in the table.

| **Plant group** | **No. species** | **Total *HSF*s** | ***HSFA*** | ***HSFB*** | ***HSFC*** |
| --- | --- | --- | --- | --- | --- |
| TERRESTRIAL | 14 | 28.0 ± 10.0 | 16.5 ± 6.6 | 8.3 ± 3.7 | 2.1 ± 1.8 |
| Eudicots | 9 | 26.1 ± 10.3 | 15.1 ± 6.5 | 8.3 ± 4.3 | 1.2 ± 0.9 |
| Monocots | 5 | 31.4 ± 9.7 | 19.2 ± 6.5 | 8.4 ± 2.5 | 3.8 ± 1.8 |
| AQUATIC (monocots) | 7 | 12.9 ± 3.3 | 7.4 ± 2.6 | 5.4 ± 1.3 | 0.0 ± 0.0 |
| Freshwater | 3 | 14.3 ± 5.1 | 9.0 ± 3.6 | 5.3 ± 1.5 | 0.0 ± 0.0 |
| Seagrasses | 4 | 11.8 ± 1.0 | 6.3 ± 0.5 | 5.5 ± 1.3 | 0.0 ± 0.0 |

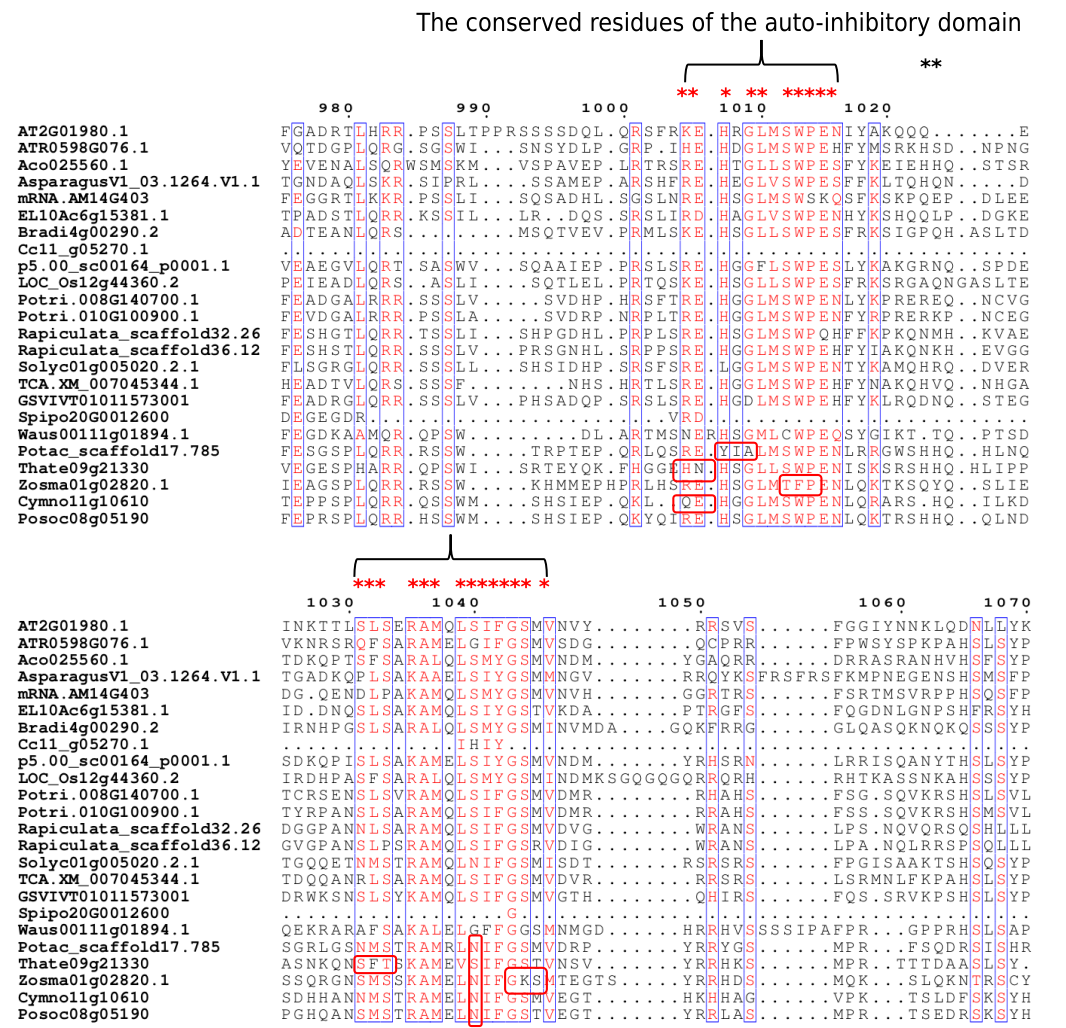

#### 5.3 Cellular salt tolerance

##### **Supplementary Fig. S5.3.1** Sequence alignment showing amino acid substitutions in regulatory domains of *SOS1* orthologs of seagrasses, indicating a diverged but convergent regulation of *SOS1*/*NHX7* in these species.

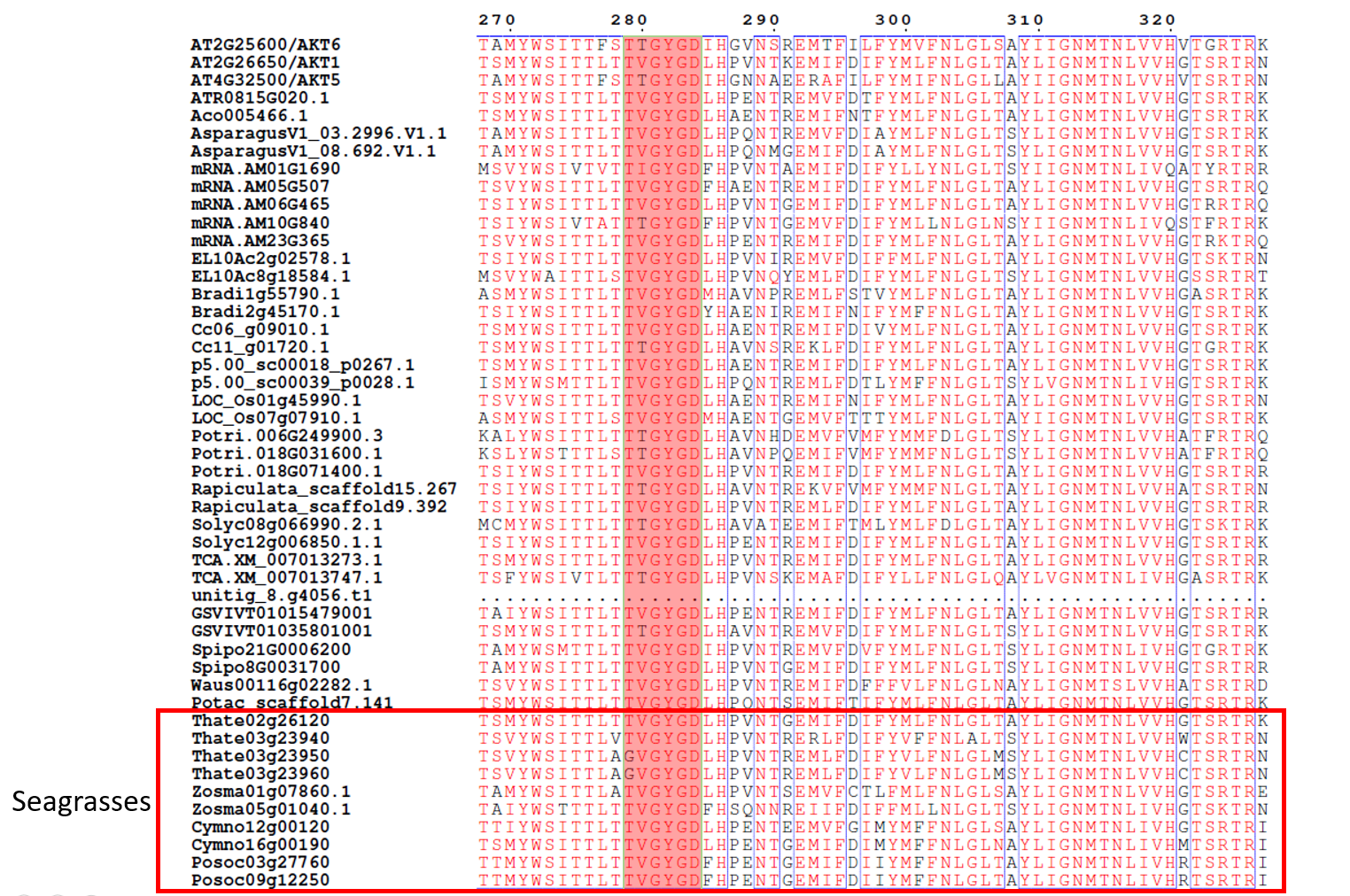

##### **Supplementary Fig. S5.3.2** Sequence alignment of *AKT5/6/1* showing the loss of Shaker-type K^+^ channels with a TTGYGD-selectivity filter in all seagrasses.

#### 5.4 Hypoxia

b)

a)

d)

c)

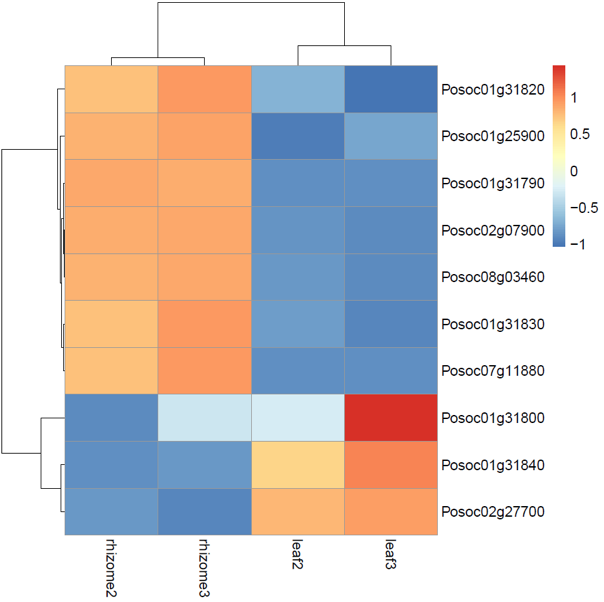

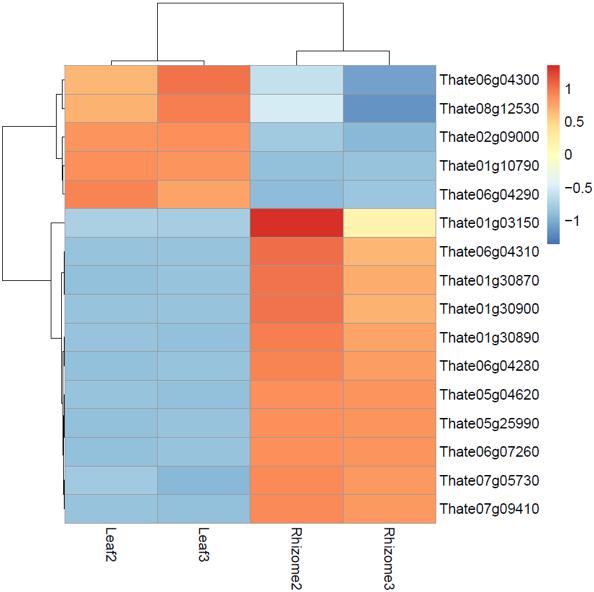

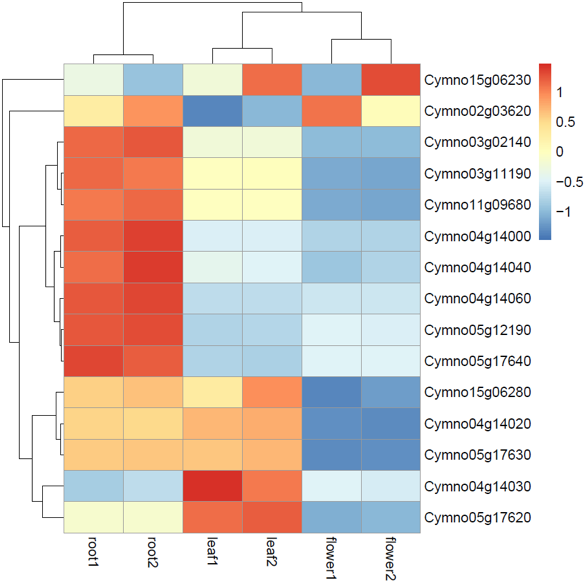

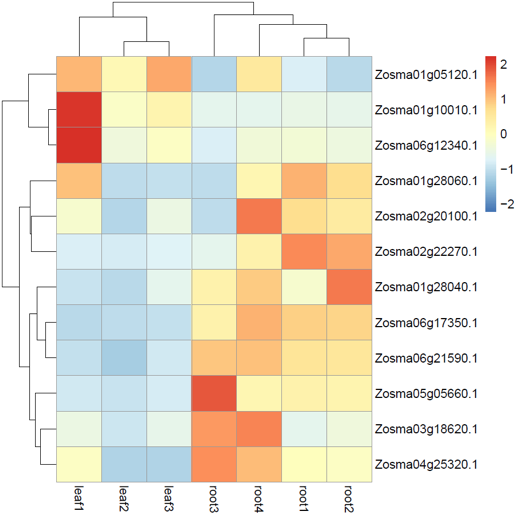

##### **Supplementary Fig. S5.4.1** Differential expression of *ERF-VII*s in the four seagrass species.

a. Leaf vs. rhizome in *P. oceanica*, b. Leaf vs. rhizome in *T. testudinum*, c. leaf vs. rhizome vs. flower in *C. nodosa*; d. leaf vs. root in *Z. marina*. Most *ERF-VII*s had higher expression in rhizomes and roots as compared to leave in four seagrasses.

##### **Supplementary Fig. S5.4.2** Syntenic relationship of genes mentioned in the main text for *P. oceanica*, *T. testudinum*, *Z. marina*, *C. nodosa*, and *P. acutifolius*.

Different colors represent different families derived from the whole-genome triplication event, as well as from the whole-genome duplication events for *C. nodosa* and *P. acutifolius*, including *ERF-VII, PCO, PFK4, LDH, SnRK1, eIFiso4G1, AVP1* genes*.* *ERF-VII* (orange lines) retained triplicated copies in *P. oceanica*, *T. testudinium* and *P. acutifolius*. *PFK4* (blue lines) retained triplicated copies in *P. oceanica*, *C. nodosa* and *P. acutifolius.* *PCO* (green lines) retained triplicated copies in *T. testudinium.*

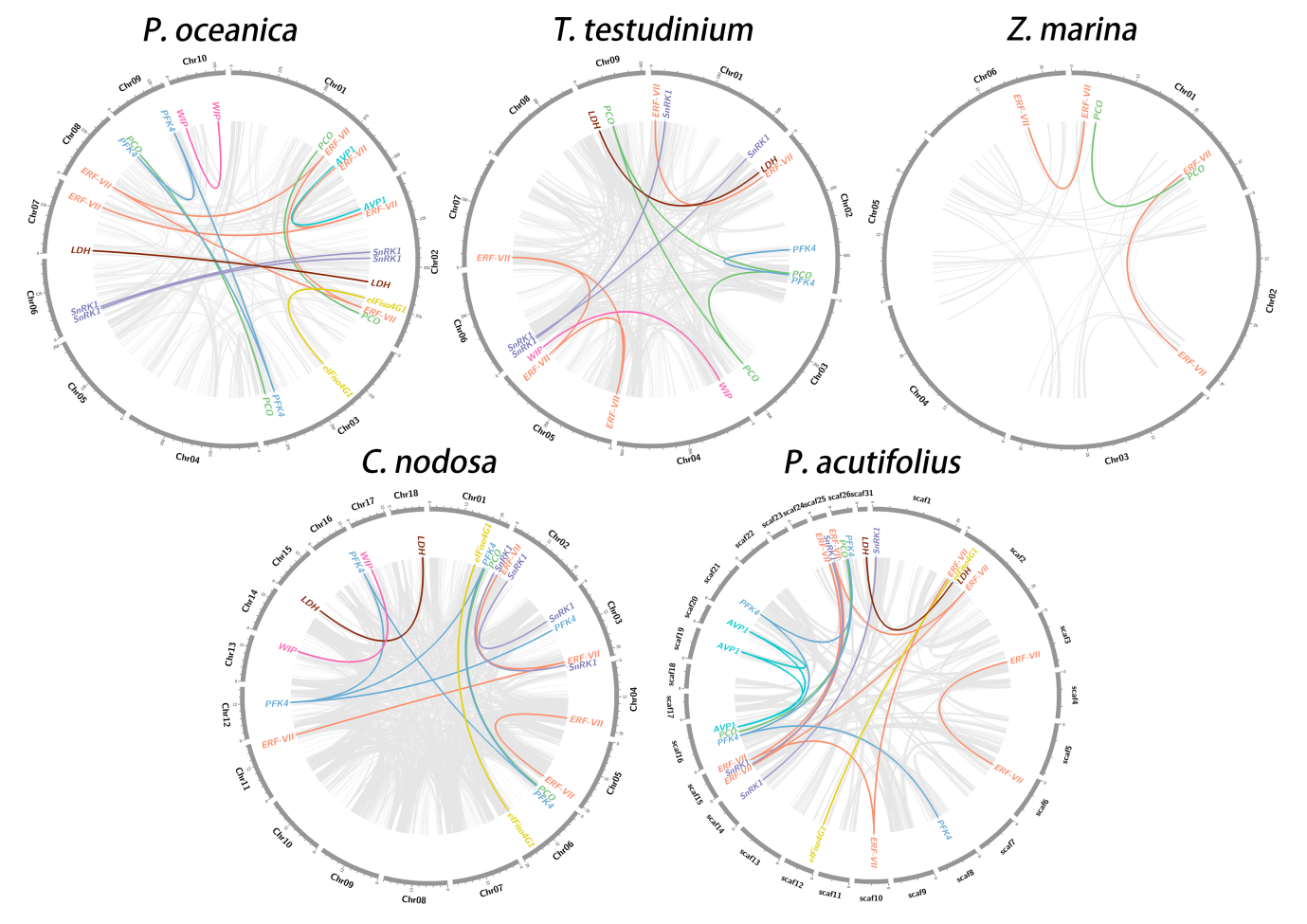

#### 5.5 Light perception, circadian clock, and photosynthetic carbon acquisition

##### **Supplementary Note S5.5.1** CO_2_-concentrating mechanisms (CCMs) and photosynthetic carbon acquisition

One of the major challenges that face seagrasses is the acquisition of inorganic carbon (Ci) for photosynthesis. Photosynthetic carbon limitation in the marine environment results from several physicochemical factors that restrict the supply of Ci to the leaf surface of seagrasses. Of the DIC (dissolved inorganic carbon) pool in seawater, the bicarbonate ion (HCO_3_^−^) accounts for nearly 90%, while the primary Ci source for RuBisCO (CO_2(aq)_) is in limited supply (roughly 1% of the DIC pool) (Campbell and Fourqurean 2013). Many algae and terrestrial plants have thus evolved various carbon-concentrating mechanisms (CCMs) to enhance their photosynthetic capabilities under Ci limitation. The occurrence of true CCMs in seagrasses is currently still a matter of debate (Larkum et al. 2017; Larkum et al. 2018), although recent findings point to the existence of biophysical CCMs and demonstrate an evolutionary adaptation of RuBisCO kinetics across submerged angiosperms (Capó-Bauçà et al. 2022). This closely resembles what seen in eukaryotic algae rather than that of terrestrial C4 plants (biochemical CCMs). The acquisition of HCO_3_^−^ for photosynthesis can occur via two (non-exclusive) basic models: (a) apoplastic conversion of HCO_3_^−^ to CO_2_ and OH^−^, catalyzed by external carbonic anhydrases (*CA*); (b) direct uptake of HCO_3_^−^ by anion transporters or an H^+^-HCO_3_^−^ symport based on H^+^‐ATPase pumps (Larkum et al. 2018). In *P. oceanica*, a direct HCO_3_^−^ uptake via a fusicoccin‐sensitive H^+^‐ATPase pump has recently been demonstrated (Rubio et al. 2017).

The four seagrass species studied, as well as *Potamogeton*, possess genes encoding for all three carbonic anhydrase gene families present in higher plants (i.e., α, β and λ) (DiMario et al. 2017). Six orthogroups (OGs) encode for *α-CA*, the most abundant family, two are associated to *β-CA* and two to *λ-CA*, respectively (Extended Data Table 9). Overall, 87% of seagrass *α-CA* are identified as extracellular/soluble proteins (Suppl. Table S5.5), which could be excreted from epidermal cells for catalyzing the conversion of HCO_3_^−^ to CO_2_ and OH^−^, hence contributing to CCMs (Larkum et al. 2017). 64% of seagrass *β-CA* are targeted to the chloroplast, generally to the chloroplast stroma or thylakoid membranes, while 100% of *λ-CA* are targeted to the mitochondria (data not shown). This confirms that seagrasses do have a requirement for an extracellular *CA* activity for adequate photosynthesis, as previously demonstrated by using chemical inhibitors (Larkum et al. 2017; Larkum et al. 2018; Capó-Bauçà et al. 2022). Interestingly, *α-CA* OG0013954 is exclusive of seagrasses (except for *T. testidinum*) and *P. acutifolius* (Extended Data Table 9). All sequences within this OG possess the typical *α-CA* domain, but most of them have variable mutations of the canonical His at the active site (data not shown), with unknown effects. Almost all OG0013954 members, as well as other *α-CA* transcripts, are highly expressed in leaf tissue (Suppl. Fig. S5.5.1), which supports the hypothesis that they could constantly increase the CO_2_ concentration in the periplasmic space, thus enhancing its diffusive transfer to RuBisCO and the photosynthetic rate in seagrass leaves. *α-CA* OG0028785 appears unique to *Z. marina*. A slight expansion of *α-CA* genes is evident in seagrasses with respect to terrestrial and freshwater/brackish-water species, when considering the average number of genes per species (terrestrial: 7; freshwater/brackish: 6; marine: 8). This increase in copy number of *α-CA* in *P. oceanica* and *P. acutifolius* results from the WGT event as well as specific tandem duplications.

A C4 or C3-C4 intermediate photosynthetic metabolism could also contribute to CCMs in seagrasses. We screened the components of the C4 photosynthesis pathway as outlined in Rao et al. (2016). The analysis of C4-pathway related genes revealed that all genes typical of terrestrial plants are present in seagrasses (Extended Data Table 9). However, this is not diagnostic of a functional C4 biochemical pathway, as they have functions other than C4 photosynthesis. In addition, none of the studied species possesses the Ser residue characteristic of C4 Phosphoenolpyruvate carboxylase (PEPC) (data not shown). Interestingly, in *C. nodosa*, which has been previously hypothesized to be a C4 species (Koch et al. 2013), there were 15 C4-related genes retained specifically after WGT or WGD events, including two encoding for PEPC (retained after WGD). Similarly, in *P. acutifolius*, 17 C4-related genes were retained following WGT or WGD (Extended Data Table 9). We cannot exclude the presence of some kind of C3-C4 intermediate metabolism at least in these species, similarly to what observed in the freshwater hydrophyte *Hydrilla verticillata* (Hydrochariraceae), where facultative single cell C4 type photosynthesis is known to occur (Rao et al. 2002).

Four orthogroups are annotated as boron transporters (HCO_3_^-^ transporter family) in the studied species (Extended Data Table 9). One orthogroup is completely absent in aquatic species (OG0012730). On average, aquatic and marine species have less genes encoding for boron transporters than terrestrial species, which seems to rule out their potential role for Ci acquisition at sea and CCMs. Regarding proton pumps (H^+^-ATPases), only three OGs encoded for plasma membrane *H^+^-ATPases* (Extended Data Table 9), whose role could be associated to bicarbonate symport for photosynthesis. Overall, seagrasses have less genes, on average per species, than terrestrial ones, but several of them were retained following WGT or WGD events (7 in *C. nodosa*, 6 in *P. oceanica* and 3 in *P. acutifolius*) (Extended Data Table 9). This confirms that *ATPase* pumps could still play a role in seagrasses for driving symport of different substances across the plasma membrane, as demonstrated by Rubio et al. (2017) in *P. oceanica* for bicarbonate uptake.

##### **Supplementary Table S5.5** Prediction of sub-cellular localizations of α-Carbonic Anhydrases (*α-CA*) in the studied seagrass species and *P. acutifolius*.

The presence of transit and signal peptides for specific cellular compartments and the determination of the protein type (soluble vs. membrane) was done by using multiple bioinformatics approaches.

| **Orthogroup** | ***C. nodosa*** | | ***P. oceanica*** | | ***T. testudinum*** | | ***Z. marina*** | | ***P. acutifolius*** | |
| --- | --- | --- | --- | --- | --- | --- | --- | --- | --- | --- |
| OG0000299 | Cymno02g02020 | Endoplasmic reticulum, Membrane | Posoc02g30740 | Extracellular, Soluble | Thate05g03940 | Sec/SPI; Extracellular, Soluble | Zosma01g08990 | Sec/SPI; Extracellular, Soluble | Potac_scaffold2.170 | Sec/SPI; Extracellular, Soluble |
|  | Cymno02g02030 | Sec/SPI; Extracellular, Soluble | Posoc02g30790 | Sec/SPI; Extracellular, Soluble | Thate05g03970 | Sec/SPI; Extracellular, Soluble | Zosma01g09000 | Sec/SPI; Extracellular, Soluble | Potac_scaffold2.171 | Sec/SPI; Extracellular, Soluble |
|  | Cymno02g02040 | Sec/SPI; Extracellular, Soluble | Posoc02g35020 | Sec/SPI; Extracellular, Soluble | Thate08g22080 | Sec/SPI; Extracellular, Soluble | Zosma06g13140 | Sec/SPI; Extracellular, Soluble | Potac_scaffold2.173 | Sec/SPI; Extracellular, Soluble |
|  | Cymno02g02050 | Extracellular, Soluble | Posoc06g06700 | Endoplasmic reticulum, Membrane | Thate08g22090 | Sec/SPI; Extracellular, Soluble | Zosma06g16770 | Sec/SPI; Extracellular, Soluble |  |  |
|  | Cymno02g02060 | Lysosome/ Vacuole, Soluble |  |  | Thate08g22100 | Sec/SPI; Extracellular, Soluble |  |  |  |  |
|  | Cymno09g06350 | Sec/SPI; Extracellular, Soluble |  |  | Thate08g22270 | Sec/SPI; Extracellular, Soluble |  |  |  |  |
| OG0002316 | Cymno04g00210 | Extracellular, Soluble | Posoc01g00410 | Sec/SPI; Extracellular, Soluble | Thate01g08320 | Cytoplasm, Soluble |  |  | Potac_scaffold2.352 | Cytoplasm, Soluble |
|  |  |  |  |  | Thate01g08350 | Sec/SPI; Extracellular, Soluble |  |  | Potac_scaffold9.158 | Sec/SPI; Extracellular, Soluble |
|  |  |  |  |  | Thate06g11790 | Sec/SPI; Extracellular, Soluble |  |  |  |  |
| OG0013954 | Cymno09g03270 | Sec/SPI; Extracellular, Soluble | Posoc08g07760 | Sec/SPI; Extracellular, Soluble |  |  | Zosma06g19600 | Sec/SPI; Extracellular, Soluble | Potac_scaffold17.661 | Sec/SPI; Extracellular, Soluble |
|  |  |  | Posoc08g07770 | Sec/SPI; Extracellular, Soluble |  |  | Zosma06g20930 | Sec/SPI; Extracellular, Soluble | Potac_scaffold17.663 | Sec/SPI; Extracellular, Soluble |
|  |  |  |  |  |  |  |  |  | Potac_scaffold18.302 | Sec/SPI; Extracellular, Soluble |
| OG0028785 |  |  |  |  |  |  | Zosma01g03190 | Sec/SPI; Extracellular, Soluble |  |  |

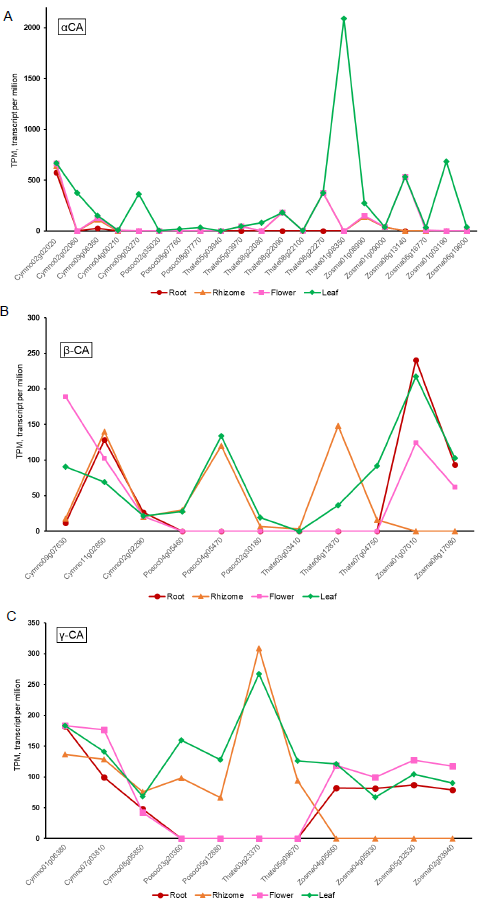

##### **Supplementary Fig. S5.5.1** Differential expression of *α-CA*, *β-CA* and *λ-CA* in root, rhizome, flower, and leaf tissues of the studied seagrass species.

##### **Supplementary Note S5.5.2** Photosynthesis

Seagrasses face different light environments according to the depth and latitude where they live. Irradiance decreases with depth, and light quality is also altered along the water column. In order to investigate if the adaptation to the light environment experienced by seagrasses in their submerged marine life imposed evolutionary adaptations resulting in expansion/reduction of gene families, the following categories of genes have been analyzed (from KEGG Photosynthesis Proteins - *Arabidopsis thaliana*): Photosystem and electron transport system: Photosystem II (P680 chlorophyll a); Photosystem I (P700 chlorophyll a); Cytochrome b6/f complex; Photosynthetic electron transport; F-type ATPase; Antenna proteins: Light-harvesting chlorophyll-protein complex A and B (*LHCA*, *LHCB*). The total number of genes present in the orthology groups for the investigated gene families are presented in the Extended data table 10. In comparison with the other Alismatales, seagrasses have a noteworthy expansion in genes of the *psA*/*psB* complexes and of the *LHCB*. Looking at this group in particular, *Posidonia*, *Thalassia* and *Zostera* have a clear expansion in comparison to *Cymodocea*, which could be related to the larger depth gradient experienced by the formers. Genes for the *psA*/*psB* complexes are coded by the chloroplastic genome, as well as for the Cytochrome b6/f complex and the F-type ATPase. Nevertheless, in most of case for seagrass species and for some of the other species included in the analysis there are also extra-copies in the nuclear genome (numbers in red in the Extended data Table 10). *T. testudinum*, for example, has 6 nuclear copies of *psbA*. Nevertheless, nuclear copies of chloroplast genes are not expressed. See for example genes for the Cyt b6/f complex in *C. nodosa* and *Z. marina* leaf and flower tissues (Suppl. Fig. S5.5.2 A and B).

Seagrass species analyzed feature different floral morphology. For two species (*Z. marina* and *C. nodosa*) gene expression data have been obtained also from floral tissue. Results indicate that *Zostera* flowers are photosynthetically active, while *Cymodocea* flowers are not. *LHC*s seem to be active at the same levels of leaf tissue while electron transport genes express at a lower level (Suppl. Fig. S5.5.2 C and D).

Photosynthesis occurs mainly in female *Zostera* flowers. This could be due to the maturation timing of male and female flowers, as suggested for *P. oceanica*, where it was also found that only female flowers expressed photosynthetic genes (Entrambasaguas et al. 2017).

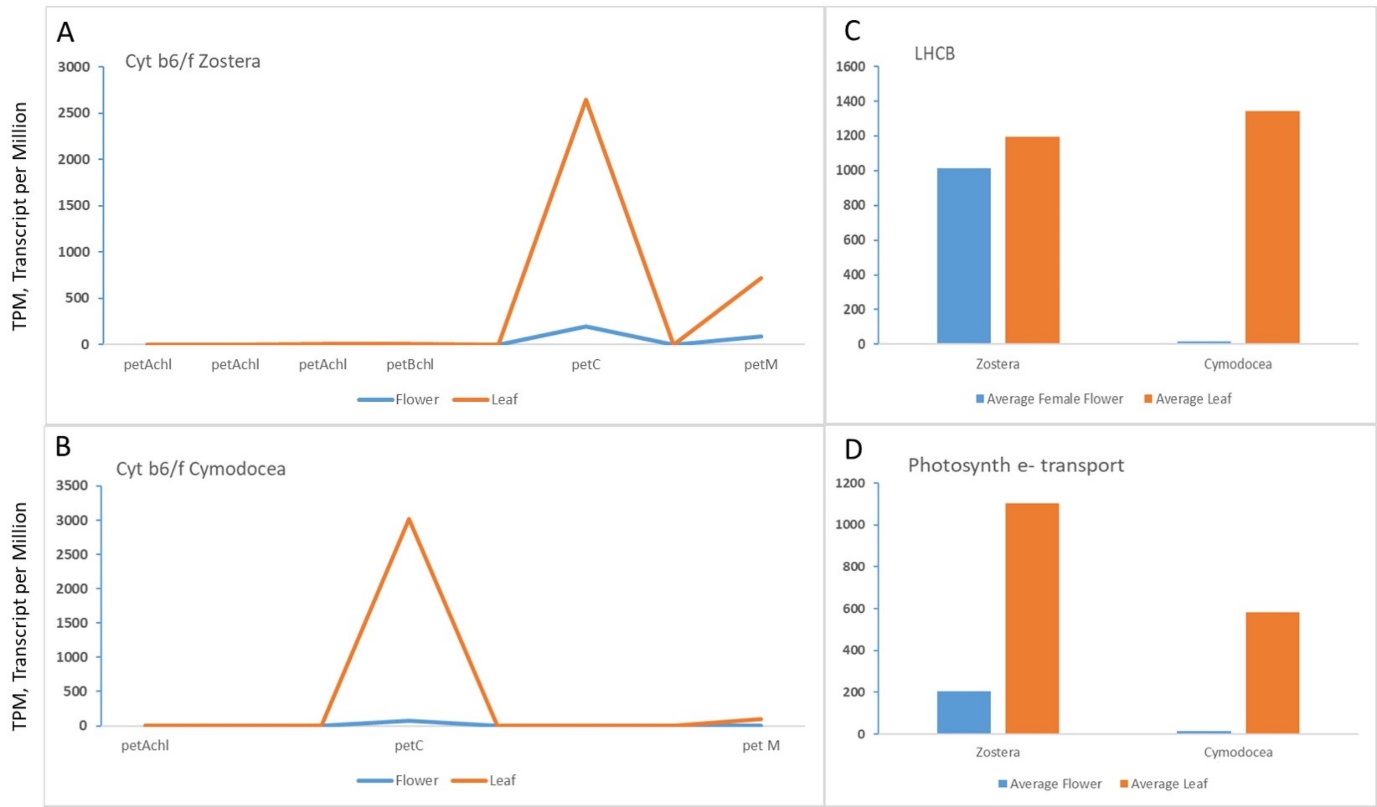

##### **Supplementary Fig. S5.5.2** Differential expression of Cyt b6/f complex, *LHCB* and electron transport genes in *Z. marina* and *C. nodosa*.

A, B: expression of nuclear copies of chloroplast genes (*chl*) and of nuclear genes in leaf and flower tissue. C, D: comparison between gene expression in leaf and flower tissue.

##### **Supplementary Note S5.5.3** Light Signaling & Circadian Clock

Seagrasses mostly conserved the full repertoire of orthologous genes for photosensory proteins and signalling systems, evolved in the green lineage during the different stages of plant terrestrialization (Extended data Table 11) (Han et al. 2019; Jing and Lin 2020). Seagrasses conserved genes for all the three classes of photoreceptors UV-A/Blue, UV-B and RED/FAR-RED typical of higher plants, with few exceptions (Extended data Table 11) as well as at least one ortholog for each of the three main classes of UVA/Blue photoreceptors, that are Phototropins (*PHO*), Cryptochromes (*CRY*) and the LOV/F-box protein (*FKF/LKP/ZTL*) (Extended data Table 11).

The UV-B receptor (UVB-Resistance 8) that is already known to be absent in *Z. marina* is still present in the other three species although the predicted protein sequence of *UVR8* in *P. oceanica* has a shorter N terminus compared to other species (data not shown) and lacks the C27 domain (Supplementary Fig. S5.5.3). This is a region of 27 amino acids from the C terminus that mediates the interaction with proteins repressor of photomorphogenesis 1, 2 (*RUP1* and *RUP2*) (Yin et al. 2015), two proteins belonging to UV-B signalling including UV-B acclimation and tolerance. *RUP1* and *RUP2* are missing in the *Z. marina* genome (Extended data Table 11). These observations indicate that UV-B tolerance and the downstream regulation signalling pathways vary among species and are related to the relative light habitat features.

The family of red receptors (Phytochromes), present in green plants and prokaryotes, was expanded in land plants after terrestrialization in response to a light environment enriched in red and far-red wavelengths (Smith 2000; Mathews 2006). The number of phytochromes varies among species: eudicots tend to have two or three phytochromes genes (up to five *PHYA*-*PHYE* in *A.thaliana*), while dicots species tend to have a lower number of genes, generally only one gene for *PHYA* and one for *PHYB*. The return to the sea did not lead to a general reduction of phytochrome genes (Extended data Table 11); indeed, all the four species investigated possess at least one gene for phytochome A (*PHYA*) and phytochrome B (*PHYB*) while *P. oceanica* and *T. testudinum* also possess an orthologous gene for *PHYC*. Furthermore, *P. oceanica* and *C. nodosa* have a unique orthologous cluster (OG0026441) for a Phytochome E (*PHYE*) (Suppl. Fig. S5.5.4) often absent in monocotyledonous plants (Smith 2000; Mathews 2006). After WDG and WGT, *P. oceanica* and *C. nodosa* as well as *P. acutifolius* retained duplicated genes for *CRY* or *PHOT*s (Extended data Table 11) while *P. oceanica* and *P. acutifolius* also for *ZTL/FKF1*.

Components of the downstream light signaling pathways such phytochrome interacting factors (*PIF*s), constitutive photomorphogenic protein 1 (*COP1*) and elongated hypocotyl 5 (*HY5*) are still present (Extended data table 11) with several orthologous genes comparable with other aquatic and land species. Also, the repertoire of transcription factors essential for photomorphogenesis and seed emergence, like the far-red elongated hypocotyl 1,3 (*FHY1/3*), far-red-impaired response1 (*FAR1*) and long after far-red light 1 (*LAF1*) is the one typical of land angiosperms. However, further functional studies must investigate if those genes have also conserved the same pattern of expression of land plants, especially during critical stages of seed setting and plant development.

Perception of surrounding light cues is critical also for the entrainment of the circadian clock system. The circadian clock regulates a plethora of processes that affect physiology and life cycle in plants, such as daily water and carbon availability and hormone signalling pathways (McClung 2019). All seagrass species, apart from *T. testudinum*, lost ortholog genes for *Timing of Cab* (*TOC1*) (Extended data Table 12). *TOC1* is one of the key clockwork components of the evening transcriptional-translational loop (Harmer 2009) belonging to the PSEUDO RESPONSE REGULATOR (*PRR*) family with a crucial function in the integration of light signals to the circadian control (Pokhilko et al. 2013). *TOC1* has also a central role in adapting plants’ physiology to drought (Legnaioli et al. 2009; Wang et al. 2020) and in regulating the day-night energy metabolism (Cervela-Cardona et al. 2021). Remarkably, *TOC1* is also lost in the freshwater *P. acutifolius* and *W. australiana,* the latter showing a reduced circadian time control of gene expression in comparison with *Arabidopsis* (Michael et al. 2021). The loss of some genes related to the circadian system in a large part of marine and freshwater species can suggest that, in the aquatic environment, the absence of some environmental stressors typical of land habitats, such as water deficit, has led to a reduction of the regulative constraints for daily management of some metabolic and developmental plant processes. Further functional studies could highlight changes in regulative networks mediated by circadian clock genes and their implication for seagrass adaptation to marine environments.

*C. nodosa*, *P. oceanica* and *T. testidinum* retained, after WGT and WGD events, one gene each related to the circadian clock and photoperiodism, respectively *LNK1*, *ZTL* and *GI* (Extended Data Table 12).

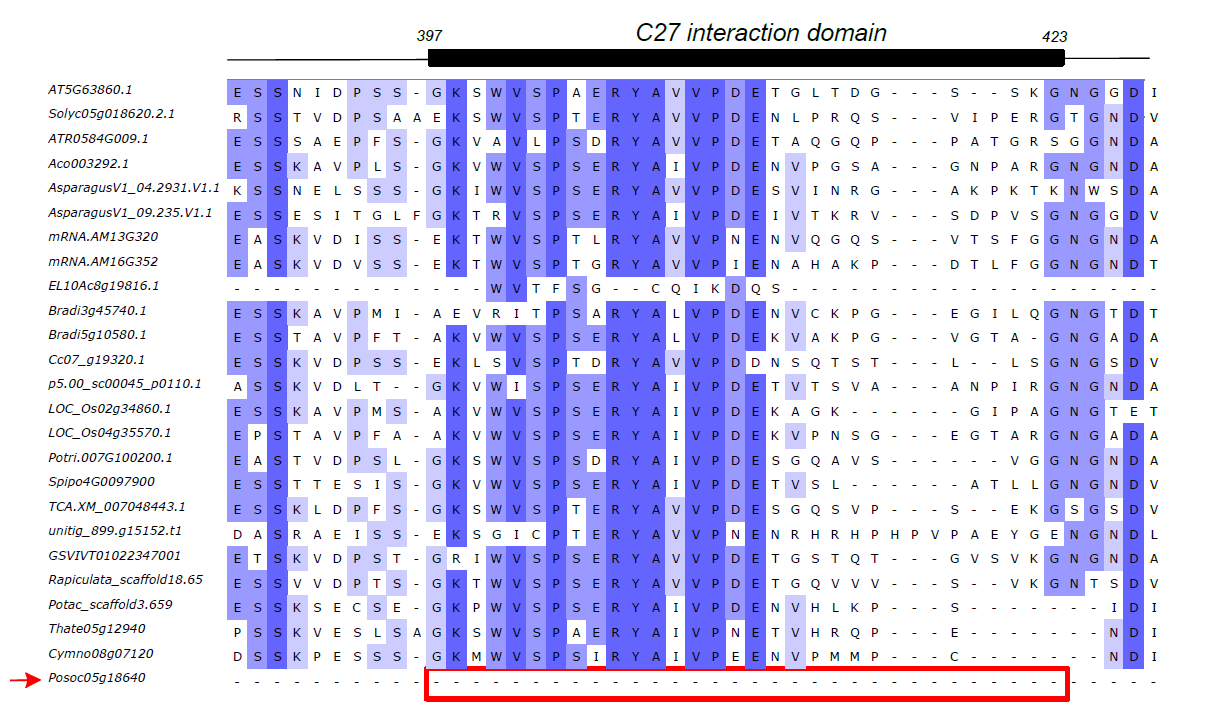

##### **Supplementary Fig. S5.5.3** The N terminus alignment of UVB-Resistance 8.

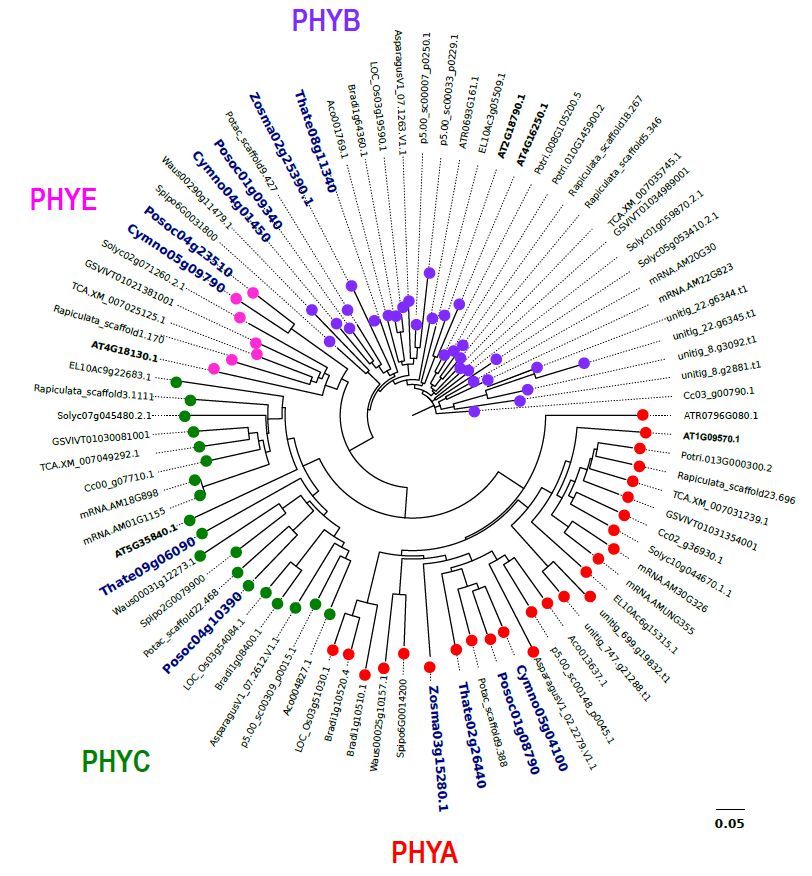

##### **Supplementary Fig. S5.5.4** Phylogenetic tree of seagrass phytochromes.

#### 5.6 NAC transcriptional factors

##### **Supplementary Note S5.6** NAC transcriptional factors

NAC proteins are one of the largest family of transcriptional factors that are involved in different developmental processes as well as in the regulation of signaling pathways in response to abiotic stressors, especially salt stress (Puranik et al. 2012). A comparable number of sequences were found in seagrasses with respect to land and aquatic species. However, specific orthogroups were found for aquatic plants, especially for *Z. marina* and *P. oceanica*. A specific functional analysis revealed that while *P. oceanica* retained JUNGBRUNNEN 1 (*JUB1*) genes in the inactive form (i.e., low expression values), *C. nodosa* which is phylogenetically more related to *P. oceanica* then the other species, expressed *JUB1* genes in leaves. *JUB1* is a central longevity regulator as well as a regulator of responses to abiotic stressors enhancing salt stress tolerance (Wu et al., 2012). In addition, other sequences annotated as *JUB1* were found across all species belonging to different orthogroups. A total of 67 sequences were found across all species included in OG0000816 orthogroup. Here too, *P. oceanica* sequences were not functionally active, only Posoc08g09120 and Posoc08g09110 were weakly expressed in leaves. Contrarily, sequences found for the other seagrasses (Cymno02g16010, Cymno02g16050, Zosma03g24220.1) showed higher expression values especially in leaves. In *T. testudinum*, one single gene copy was found (Thate04g03560) specifically expressed in root and leaf. Thus, a phylogenetic tree was built including these sequences to visualize relationships of the *JUB1* sequences between seagrasses (Suppl. Fig. S5.6). Sequences of *P. oceanica* and *C. nodosa* belonging to the same orthogroup (OG0000816) formed a single clade, apart from *T. testudinum*. In conclusion, the functional re-organization observed for *JUB1* could be related to the different ecological requirements of these species, modulating salt stress tolerance in seagrasses. *P. oceanica*, in fact, colonizes open coastal habitats with a very narrow range of salinity, contrary to *C. nodosa* which can also be present in estuarine systems with highly variable salinities.

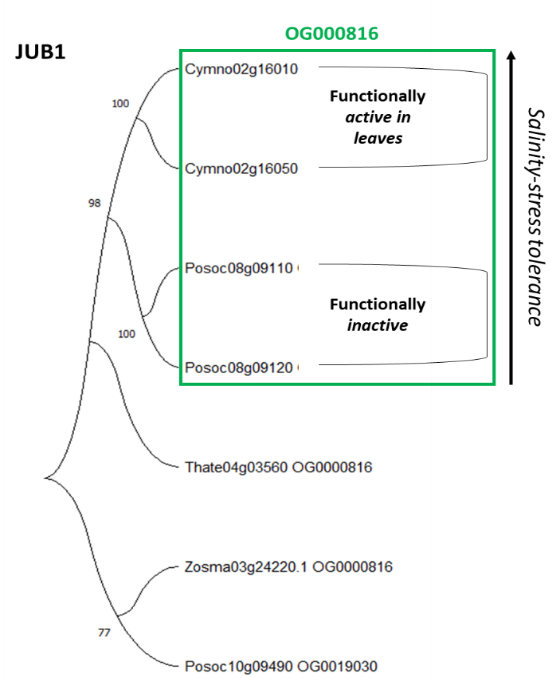

##### **Supplementary Fig. S5.6** Evolutionary analysis by Maximum Likelihood method of *JUB1* in seagrasses.

#### 5.7 Nitrogen metabolism

##### **Supplementary Note S5.7** Nitrogen metabolism

Seagrass meadows act as an important nitrogen (N) filter in the coastal environments. In this context, seagrasses assimilate large amounts of N, and exude oxygen and labile carbon into sediments, which stimulate other processes in the nitrogen metabolism pathway, including nitrification-denitrification process that counterbalance the net N loads through microbial transformation (Zarnoch et al. 2017; Aoki et al. 2020). The key genes linked to nitrogen uptake/transport and assimilation were retained in all of the plants examined (Extended table 13). This corresponds to at least 66 and 25 orthogroups which functions in uptake/transport and assimilation, respectively. Their existence is essential because an efficient N metabolism process is required to ensure normal growth and development in plants, regardless of their diversity, habitat or nature. Moreover, seagrasses may have acquired a more effective nitrogen metabolism in N-deficient marine environments through symbiotic N2-fixing bacteria, that could have facilitated the migration of flowering plants back to the sea some 100 million years ago (Mohr et al. 2021).

As compared to non-seagrass genomes, the nitrate transporter (*NRT*) gene families of seagrasses were contracted (40.71%), indicating that seagrasses may have evolved alternative mechanisms to utilize nitrogen sources more effectively (Suppl. Table S5.7 and Extended table 13). Other than nitrate, seagrasses rely on ammonium as a primary source of nitrogen (Touchette and Burkholder 2001; Xu et al. 2020), particularly when exposed to anoxic conditions in marine sediment where nitrate is scarce due to disrupted ammonium-to-nitrate oxidation. In this case, ammonium is metabolized directly via GOGAT pathway, which catalyse the formation of glutamine from glutamate and ammonium, instead of converting nitrate to ammonium prior to glutamine formation (Wang et al. 2021).

Nitrate reductase (*NR*) was expressed in all parts of seagrasses (flower, root, vegetative, rhizome), with *NR* robustly expressed in the root of *Zostera marina* and the leaves of *Cymodocea nodosa* and *Thalassia testudinum*. *NR* activity is widely influenced by light (Touchette and Burkholder 2001) and given that *NR* activity is highest during photosynthetic periods and lowest in the dark, nitrite reduction occurs more frequently in leaf tissues than in root tissues in most seagrasses, suggesting the importance of light on *NR* response (Manassa et al. 2017; Wang et al. 2021; Jiménez-Ramos et al. 2022). *Z. marina* is unique from the other seagrasses as its *NR* activity can be maintained in the dark, provided that the environment is nitrate-enriched and tissue carbohydrate levels are high(Touchette and Burkholder 2001). The intensity and duration of *NR* activity is directly parallel with the soluble carbohydrate supplies (Touchette and Burkholder 2007). On the other hand, seagrass leaves such as those in *T. testudinum* tend to have higher efficiency of nitrogen assimilation compared to root, given that the NH4+ in the water column is relatively lower than the sediments where seagrass inhabit (Lee and Dunton 1999; Cornelisen and Thomas 2004).

##### **Supplementary Table S5.7** Comparison of nitrate transporter (*NRT*) gene families between seagrass and non-seagrass species.

| Orthogroups | Number of gene copy in four seagrass species in this study | Number of gene copy in non-seagrass species |
| --- | --- | --- |
| OG0000120 | 19 | 133 |
| OG0000306 | 14 | 81 |
| OG0000367 | 11 | 78 |
| OG0000480 | 4 | 76 |
| OG0000886 | 0 | 64 |
| OG0000925 | 6 | 54 |
| OG0002990 | 5 | 34 |
| OG0003549 | 4 | 33 |
| OG0003600 | 4 | 31 |
| OG0008152 | 5 | 19 |
| OG0010200 | 4 | 18 |
| OG0013783 | 0 | 9 |
| OG0016342 | 1 | 3 |
| OG0016485 | 2 | 3 |
| OG0017868 | 0 | 4 |
| OG0023822 | 1 | 1 |
| Total | 80 | 641 |
| Average | 20 | 33.7 |
